# Novel lipid biomarkers for algal resistance to viral infection in the ocean

**DOI:** 10.1101/2022.09.14.507897

**Authors:** Guy Schleyer, Constanze Kuhlisch, Carmit Ziv, Shifra Ben-Dor, Sergey Malitsky, Daniella Schatz, Assaf Vardi

## Abstract

Marine viruses play a key role in regulating phytoplankton populations, greatly affecting the biogeochemical cycling of major nutrients in the ocean. Resistance to viral infection has been reported for various phytoplankton species under laboratory conditions. Nevertheless, the occurrence of resistant cells in natural populations is underexplored due to the lack of sensitive tools to detect these rare phenotypes. Consequently, our current understanding of the ecological importance of resistance and its underlying mechanisms is limited. Here, we sought to discover lipid biomarkers for the resistance of the bloom-forming alga *Emiliania huxleyi* to its specific virus, *E. huxleyi* virus (EhV). We identified novel glycosphingolipids (GSLs) that characterize resistant *E. huxleyi* strains by applying an untargeted lipidomics approach. Further, we detected these lipid biomarkers in *E. huxleyi* isolates that were recently collected from *E. huxleyi* blooms and used them to detect resistant cells in the demise phase of an open ocean *E. huxleyi* bloom. Lastly, we show that the GSL composition of *E. huxleyi* cultures that recover following infection and gain resistance to the virus resembles that of resistant strains. These findings highlight the metabolic plasticity and co-evolution of the GSL biosynthetic pathway and underscore its central part in this host-virus arms race.

## Introduction

Viruses are the most abundant biological entities in the marine environment and serve as major evolutionary and biogeochemical drivers in the oceans^1–4^. Algae-infecting viruses are estimated to turn over a substantial portion of the photosynthetically-fixed carbon, thus fueling microbial food webs, short-circuiting carbon transfer to higher trophic levels and promoting its export to the deep sea^5–7^. Recent developments allow to better quantify infected cells in the natural environment^8–13^, yet studying host-virus dynamics in natural populations^14^ remains a major challenge for our understanding of the possible phenotypic outcomes of viral infection.

The ongoing evolutionary arms race between algae and their viruses leads to diverse defense strategies, supported by continuous genetic and phenotypic adaptations of the algal cells^15–17^. Resistance to viral infection has been reported for several algal species, both as isolates from natural populations and as sub-populations that emerge following infection under laboratory conditions^16, 18–22^. Nevertheless, the prevalence of resistant phenotypes in nature is currently unknown, as we lack sensitive tools to detect resistant cells in mixed populations, hindering our understanding of their ecological importance.

The cosmopolitan alga *E. huxleyi* and its specific virus, *E. huxleyi* virus (EhV), are an attractive model system to study host-virus interactions. *E. huxleyi* forms vast annual blooms in the ocean that play an important role in regulating the global biogeochemical cycling of carbon and sulfur^23–26^ and are routinely infected and terminated by EhV^27–30^. Laboratory-based studies revealed that viral infection leads to profound rewiring of the *E. huxleyi* metabolism, including changes in glycolysis, elevated fatty acid (FA) synthesis and alterations in the cellular lipid content and composition^31–34^. Particularly, EhV is the only virus known to date to encode almost a complete pathway for sphingolipid (SL) biosynthesis, resulting in the production of structurally distinct virus-derived glycosphingolipids (vGSLs) by infected cells^35–37^. vGSLs were found to trigger host programmed cell death and are central components of the EhV membranes^37, 38^. In addition, *E. huxleyi* cells produce host-derived GSLs (hGSLs), which are found in all *E. huxleyi* strains and serve as a proxy for healthy cells^38, 39^, and sialic acid GSLs (sGSLs), which characterize susceptible *E. huxleyi* strains and were suggested to be involved in viral attachment and entry^38^. Given their structural variability and diverse roles, SLs are key players in the arms race between *E. huxleyi* and its virus.

Resistance to infection by EhV has been described in several *E. huxleyi* strains and was previously attributed to ploidy level, genome and transcriptome variations between the strains^16, 40^, to expression and activity of specific enzymes, such as DMSP-lyase, and to metacaspase expression^22, 41^. Resistant cells were also identified in low numbers (<1%) in infected *E. huxleyi* cultures^42^, revealing that resistance can also be triggered by viral infection. These resistant cells were found to be morphologically distinct from their susceptible progenitors, indicating the involvement of a life-phase transition and highlighting the phenotypic plasticity within *E. huxleyi* populations during infection^16, 42^. Nevertheless, the metabolic basis of *E. huxleyi*’s resistance to viral infection is unknown, as is the prevalence of resistant *E. huxleyi* cells in natural populations. In this study, we aimed at addressing this conundrum by identifying specific lipid biomarkers for resistant *E. huxleyi* cells and applying them to natural mixed populations.

## Results

### Untargeted lipidomics profiling of virus-resistant and susceptible *E. huxleyi* strains

To identify lipids that are characteristic of resistant strains, we compared the lipidome of four *E. huxleyi* strains that differ in their susceptibility to viral infection by EhV201 (hereinafter, EhV): the resistant *E. huxleyi* strains CCMP373 and CCMP379 and the susceptible *E. huxleyi* strains CCMP2090 and CCMP374 (hereinafter, *E. huxleyi* strains 373, 379, 2090 and 374, respectively) ^22, 38, 40^. Previous studies reported that following infection of *E. huxleyi* cultures by EhV in the lab, a small proportion of the population (< 1%) can survive and acquire resistance to the virus^16, 42^. We were therefore interested to delineate possible correlations between the lipid profile of resistant strains and the evolving resistant cells within infected susceptible cultures.

The lipidome of the resistant and susceptible strains in the presence and absence of the lytic virus EhV201 was compared over a three-day time course using liquid chromatography-high resolution mass spectrometry (LC-HRMS)-based untargeted lipidomics. All untreated cultures grew throughout the experiment, reaching 1.0-2.4×10^6^ cells per mL (Fig. 1a). The resistant *E. huxleyi* strains 373 and 379 grew throughout the experiment regardless of the presence of EhV and with no accumulation of virions in the media (Fig. 1b). In contrast, upon addition of EhV, the susceptible *E. huxleyi* strains 2090 and 374 showed growth arrest one day post infection (dpi) and were subsequently lysed (Fig. 1b). Concomitantly, accumulation of virions was detected in the medium of the infected cultures starting from 1 dpi. In all cultures, cells were harvested at four different time points (0, 1, 2 and 3 days) for lipid extraction and untargeted lipidomics analysis.

**Figure 1:**
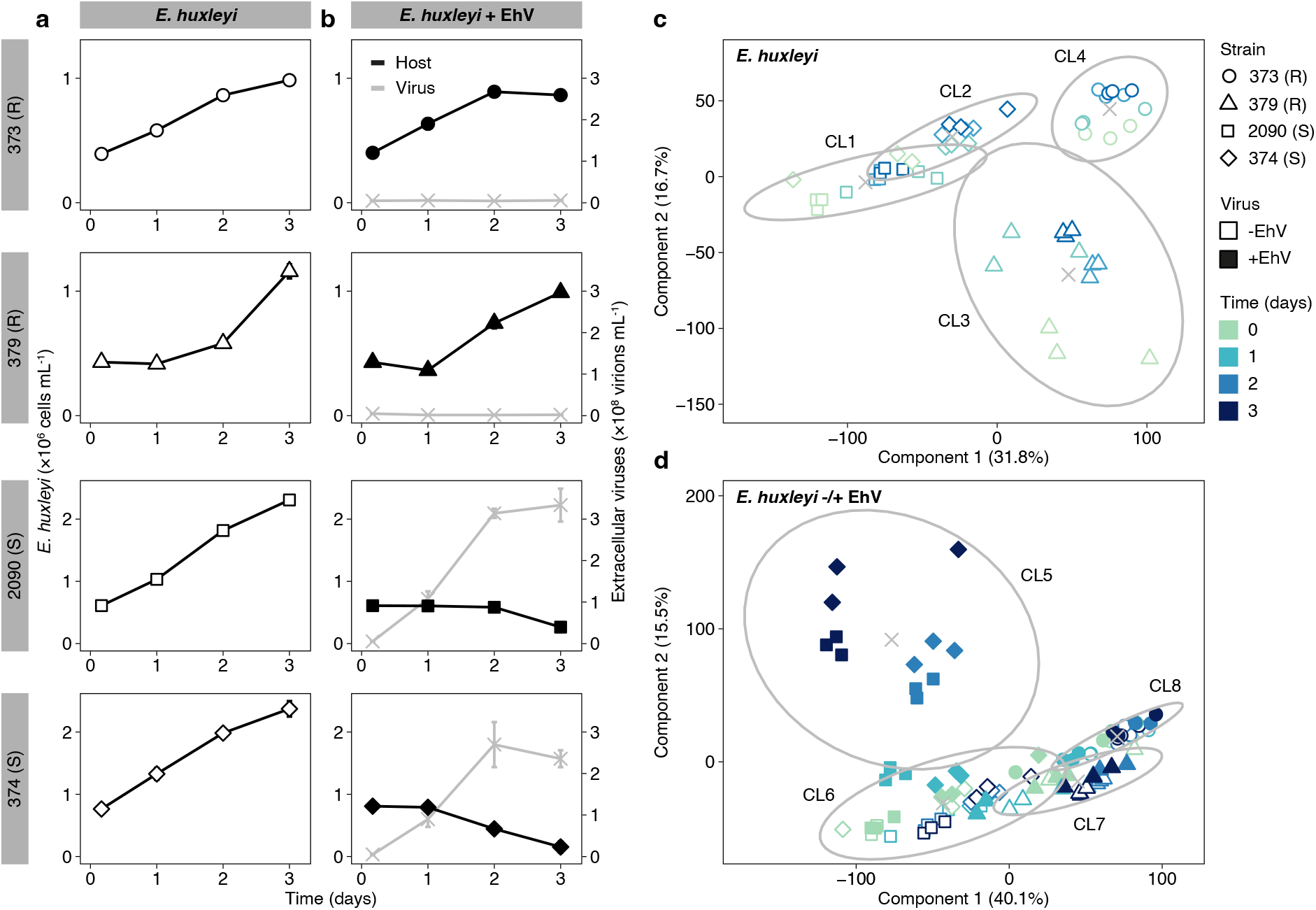
Untargeted LC-HRMS-based lipidomics analysis reveals differences between virus-resistant and susceptible *E. huxleyi* strains. (**a**) Cell abundance during growth of *E. huxleyi* strains that differ in their susceptibility to viral infection: the resistant (R) *E. huxleyi* strains 373 and 379 and the susceptible (S) *E. huxleyi* strains 2090 and 374. (**b**) Cell abundance (black lines) and production of virions (grey lines) following the addition of EhV. Values for (a) and (b) are presented as the mean ± SD (*n* = 3). (**c**) Clustering of resistant and susceptible *E. huxleyi* strains based on untargeted lipidomics (using 12,190 mass features) and *k*-means clustering (*k* = 4, Fig. S1a), as visualized by PCA. (**d**) Clustering of resistant and susceptible *E. huxleyi* strains in the presence and absence of EhV based on untargeted lipidomics (using 12,190 mass features) and *k*-means clustering (*k* = 4, Fig. S1b), as visualized by PCA. Percentage of explained variance is stated in parentheses. Each cluster (CL) is surrounded by an ellipse, with the mean marked by ‘×’. Next, we focused on mass features that were differential between the resistant and susceptible clusters in the cultures without EhV (resistant clusters 3 and 4 vs susceptible clusters 1 and 2, Fig. 1c) using a comparative analysis (one-way ANOVA with false discovery rate (FDR)-correction). By doing so, we could reduce the data to 173 differential mass features (*p* < 0.01). Following feature deconvolution and manual curation, these mass features were grouped into 43 putative lipid species (Table S1). We then applied two-dimensional hierarchical clustering to this subset of 43 putative lipid species using the complete dataset (that is, with and without addition of EhV; Fig. 2a and Fig. S2). This subset of lipid species recapitulated the previously observed separation (Fig. 1d) between the resistant and susceptible strains (two main clusters, separating *E. huxleyi* 379 and 373 from *E. huxleyi* 374 and 2090), and between each pair of strains. Similarly, while there was no clear separation between resistant strains in the presence and absence of EhV, the susceptible strains infected with EhV were clustered separately from the uninfected cultures as early as 1 dpi.

First, we compared the lipidome of the four strains in the absence of EhV. Unsupervised *k*-means clustering of the extracted data (*n* = 48; 12,190 mass features, *k* = 4, Fig. S1a), visualized by principal component analysis (PCA), separated the strains into four distinct clusters (clusters 1-4, Fig. 1c). The first PC axis (31.8%) revealed a clear separation between the susceptible and resistant strains (clusters 1 and 2 vs clusters 3 and 4, respectively), and the second PC axis (16.7%) highlighted further differences between the strains. Next, we applied *k*-means clustering to the combined dataset of cultures with and without addition of EhV (*n* = 96; 12,190 mass features, *k* = 4, Fig. S1b), which showed a clear separation between susceptible and resistant strains (clusters 5 and 6 vs clusters 7 and 8, respectively) along the first PC axis (40.1%, Fig. 1d). The second PC axis (15.5%) further separated the susceptible strains at late infection stages (2 and 3 dpi; cluster 5) from early infection stages (0 and 1 dpi) and the uninfected cultures (cluster 6).

The 43 putative lipid species were grouped into two main clusters, each further divided into two sub-clusters (Fig. 2a): (i) lipids with higher intensity in the resistant strains (especially *E. huxleyi* 379), of which most had higher intensity also in infected *E. huxleyi* 2090 cultures; (ii) lipids with higher intensity in both resistant strains; (iii) lipids with higher intensity in the resistant *E. huxleyi* strain 373 and the susceptible *E. huxleyi* strain 374; and (iv) lipids with higher intensity in *E. huxleyi* strain 374 or in both susceptible strains. Out of the 43 putative lipid species, 21 were higher in one or both resistant strains (sub-clusters i and ii). Some of these species were elevated in the resistant *E. huxleyi* strain 379 compared to *E. huxleyi* strain 373, shedding light on possible metabolic differences between these two resistant strains. Seven putative lipid species were higher in the susceptible *E. huxleyi* strains 374 and 2090 (sub-cluster iv), one of which was identified as the known sGSL d18:2/c22:0^38^ and three of which were higher in *E. huxleyi* 374 compared to *E. huxleyi* 2090.

**Figure 2:**
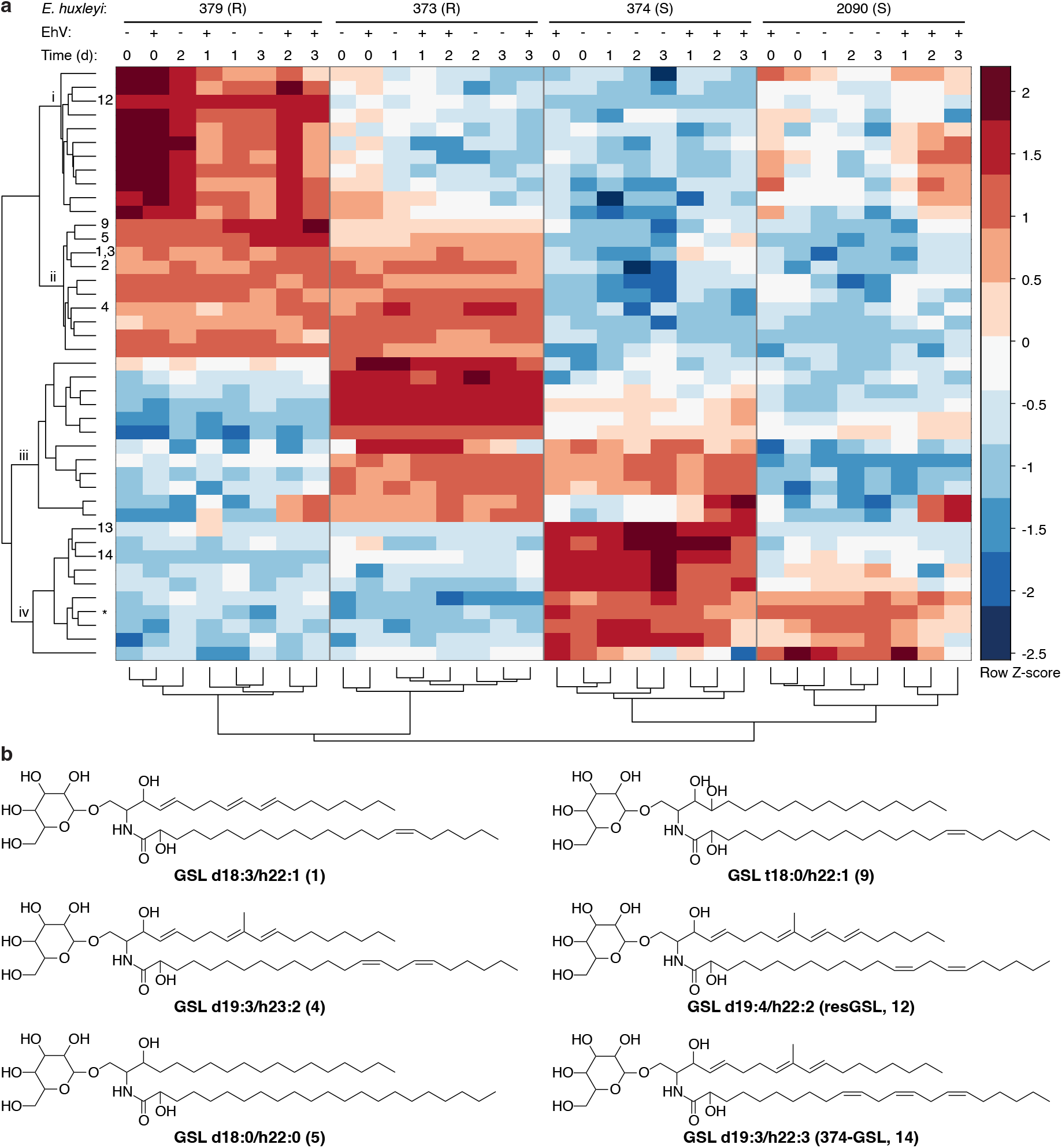
Putative lipid biomarkers for *E. huxleyi* strains differing in their susceptibility to viral infection. (**a**) Two-dimensional hierarchical clustering of 43 putative lipid species (Table S1) in four *E. huxleyi* strains in the presence and absence of EhV throughout a time course of four days (*n* = 32). Clustering was performed on log-transformed and standardized mean peak areas (*n* = 3) of the adduct ion with the highest intensity (see Fig. S2 for the non-averaged data). Samples are grouped into two main clusters that separate the resistant (R) strains from the susceptible (S) ones. Each cluster forms two sub-clusters that further separate the strains. The putative lipid species are divided into four sub-clusters (i-iv). Nine identified GSL species are marked by numbers (Table 1). The peak areas of GSLs **1** and **3** (structural isomers with a similar retention time, see Table 1), were integrated together. *sGSL d18:2/c22:0. (**b**) Putative structures of six of the previously undescribed GSL species in the *E. huxleyi*-EhV model system, which are differential between the resistant and susceptible *E. huxleyi* strains. See Fig. S3 for putative structures of all GSL species identified in this study. The structures, including LCB and FA composition, were determined based on LC-MS/MS analysis (Fig. S4-S12). The positions of the double bonds and functional groups were assigned based on the most common structures in the Lipid Maps Structure Database (LMSD) ^44^.

**Table 1:** Putative annotation and identification of GSL species that differ between resistant and susceptible strains and are previously undescribed in the *E. huxleyi*-EhV.

| Group | # | GSL species<br>LCB/FA | RT<br>(min) | Measured *m/z*<br>([M+H]<sup>+</sup>) | Predicted<br>formula |
| --- | --- | --- | --- | --- | --- |
| A<br>Higher in resistant | 1 | d18:3/h22:1 | 13.12 | 794.6107 | C<sub>46</sub>H<sub>83</sub>NO<sub>9</sub> |
|  | 2 | d18:3/h22:2 | 12.47 | 792.5980 | C<sub>46</sub>H<sub>81</sub>NO<sub>9</sub> |
|  | 3 | d19:3/h21:1 | 13.03 | 794.6107 | C<sub>46</sub>H<sub>83</sub>NO<sub>9</sub> |
|  | 4 | d19:3/h23:2 | 13.18 | 820.6278 | C<sub>48</sub>H<sub>85</sub>NO<sub>9</sub> |
| B<br>Only in resistant<br>and during infection | 5 | d18:0/h22:0* | 14.44 | 802.6722 | C<sub>46</sub>H<sub>91</sub>NO<sub>9</sub> |
|  | 6 | d18:0/h22:1<sup>†</sup> | 14.22 | 800.6600 | C<sub>46</sub>H<sub>89</sub>NO<sub>9</sub> |
|  | 7 | d18:1/h22:1* | 14.01 | 798.6440 | C<sub>46</sub>H<sub>87</sub>NO<sub>9</sub> |
|  | 8 | t18:0/h22:0* | 14.00 | 818.6702 | C<sub>46</sub>H<sub>91</sub>NO<sub>10</sub> |
|  | 9 | t18:0/h22:1 | 13.77 | 816.6531 | C<sub>46</sub>H<sub>89</sub>NO<sub>10</sub> |
|  | 10 | t18:0/h22:2<sup>‡</sup> | 13.18 | 814.6346 | C<sub>46</sub>H<sub>87</sub>NO<sub>10</sub> |
| C<br>Only in resistant | 11 | d19:4/h22:1 (resGSL) | 12.92 | 806.6127 | C<sub>47</sub>H<sub>83</sub>NO<sub>9</sub> |
|  | 12 | d19:4/h22:2 (resGSL) | 12.25 | 804.5975 | C<sub>47</sub>H<sub>81</sub>NO<sub>9</sub> |
| D<br>Only in the susceptible<br>*E. huxleyi* 374 | 13 | d19:3/h22:2 (374-GSL)** | 12.98 | 806.6143 | C<sub>47</sub>H<sub>83</sub>NO<sub>9</sub> |
|  | 14 | d19:3/h22:3 (374-GSL)** | 12.34 | 804.5981 | C<sub>47</sub>H<sub>81</sub>NO<sub>9</sub> |

We putatively annotated nine lipid species as GSLs using characteristic neutral losses and fragments of long-chain bases (LCBs) and amino fatty acids (FAs, based on MS/MS spectra, **1-5**, **9**, **12-14**, see Table 1, Fig. S3-S12 and Table S1). These GSL species varied in their LCB composition, including dihydroxylated LCBs d18:0, d18:3, d19:3 and d19:4, and the trihydroxylated LCB t18:0 (Fig. 2b). We manually identified five additional GSL species with the same LCB composition that had higher intensity in the resistant strains (**6-8**, **10-11**, see Table 1, Fig. S3, Fig. S13-S17 and Table S2; these GSL species were filtered out in the initial data preprocessing). We classified these GSL species into four groups based on their abundance in the different strains (Table 1, Fig. S18 and Fig. S19): (A) GSL species that are highly abundant in the resistant strains compared to susceptible strains (**1-4**, difference of >1 order of magnitude). These GSL species contain LCB d18:3 and d19:3; (B) GSL species that are found in resistant strains and in infected susceptible strains (**5-10**). These contain LCB d18:0, d18:1, and t18:0. (C) GSL species that are found only in the two resistant strains, with higher abundance in *E. huxleyi* 379 compared to *E. huxleyi* 373 (**11-12**, Fig. S19). These contain LCB d19:4 and were termed resistance-specific GSLs (resGSLs) due to their detection in resistant strains and their absence in susceptible strains. (D) GSL species that are found only in *E. huxleyi* 374 (**13-14**). These contain LCB d19:3 and were termed *E. huxleyi* 374-specific GSLs (374-GSLs).

GSL species containing LCBs d18:1, d18:3 and d19:4 were not detected thus far in the *E. huxleyi*-EhV system. LCBs d18:0, d19:3 and t18:0 were previously reported in the *E. huxleyi*-EhV system: LCB d19:3 in hGSL species and LCB d18:0 and t18:0 in infection-derived GSL and ceramide species (Table S3) ^36, 43^. Intriguingly, GSL species containing LCB t18:0 (**8-10**), which were detected in resistant strains and in infected cultures (Fig. S18), varied in their FA composition: resistant strains produce GSL species with a clear preference for mono- and di-unsaturated FAs over saturated ones (h22:1 and h22:2 vs h22:0, Fig. S18). Infected cultures, on the other hand, produce GSL species with saturated and mono-unsaturated FAs, as was previously described for t17:0-based vGSL species^36, 37^. Importantly, trihydroxylated LCBs were previously found only in vGSL species and were considered a unique attribute of viral infection, derived from the virus-encoded biosynthetic pathway. The tetra-unsaturated LCB d19:4, on the other hand, appears only in resGSLs found resistant strains, and therefore, we suggest that these unique resGSLs can be used as a biomarker for resistant cells in natural populations. Detection of GSL species with tetra-unsaturated LCB and trihydroxylated LCB in resistant strains suggests the involvement of specific modifying enzymes in these strains.

### Potential enzymes involved in modulating GSL composition in resistant strains

The detection of resGSL species with LCB d19:4 (**11-12**), which contains an additional double bond compared to the LCB d19:3 found in hGSL species (Table S3), indicates the involvement of an additional sphingolipid desaturase (SLD) in resistant strains, which would be responsible for the fourth double bond. A gene encoding a putative SLD was previously identified in *E. huxleyi* (*sld2*) ^31, 47^, and we identified four additional genes based on the *E. huxleyi* genome and expressed sequences (*sld1*, *sld3*-*sld5*, Table S4). Phylogenetic analysis of the conserved domain of the SLD proteins revealed three distinct clades (I-III, Fig. 3a and Table S5), each consisting of diverse taxonomic groups. Out of the five putative *E. huxleyi* SLDs, SLD1 clustered together with a viral SLD (EhV201 SLD, AET97947.1, clade I). We further examined the expression of these genes using previous transcriptomics experiments with *E. huxleyi* strains 373, 379, 2090 and 374^40, 48^. *sld1* was expressed in the resistant *E. huxleyi* strains 373 and 379 and not in the susceptible strains (Fig. 3c and Fig. S20a), suggesting that the viral and resistant-host enzymes share a similar role in the GSL biosynthetic pathway. Notably, *sld4* was also differentially expressed in the resistant strains, however, the protein falls into a different clade than the viral SLD (clade III). Therefore, *sld4* is a possible candidate for the formation of the fourth double bond in resGSLs (**11-12**), which were detected only in resistant strains. The other genes (*sld2*, *sld3* and *sld5*) were expressed in all strains (Fig. S20a and Fig. S21a). We were further intrigued to identify possible similarities between the viral and the host biosynthetic pathways that are responsible for the production of GSL species with trihydroxylated LCBs in infected and resistant cells. Previous studies suggested that the characteristic trihydroxylation of the LCB in infection-derived vGSL species is facilitated by a viral sphingoid base hydroxylase (EhV201 SBH, AET97919.1) ^36, 49^, which is highly expressed at early stages of infection (Fig. S22). LCB t17:0 is the major LCB in vGSL species, while LCB t16:0 and t18:0 are found in lower abundances^36^. In GSL species of resistant strains (**8-10**), on the other hand, only LCB t18:0 was detected. A gene encoding a putative SBH was previously identified in *E. huxleyi* (*sbh1*) ^31, 40^, and we identified six additional genes based on the *E. huxleyi* genome and expressed sequences (*sbh2*-*sbh7*, Table S4). Phylogenetic analysis of the conserved domain of the SBH proteins revealed that the *E. huxleyi* SBHs do not form a clade together but rather show similarities to diverse phyla, indicating different evolutionary origins (Fig. 3b and Table S6). Interestingly, SBH4 and SBH5 clustered together with the viral SBH, indicating a possible host-virus co-evolution. Out of the seven SBHs, *sbh*4 and *sbh*5 were highly expressed in the resistant *E. huxleyi* strains 373 and 379 and not in the susceptible *E. huxleyi* strains 2090 and 374 (Fig. 3d and Fig. S20b). Concomitantly, *sbh2* was differentially expressed in the susceptible strains, while *sbh1* and *sbh6* were expressed in all four strains. *sbh7* was detected in all four strains, with higher expression in infected *E. huxleyi* 2090 cultures. The expression of *sbh3* was not detected in all strains and conditions tested (Fig. S20b and Fig. S21b). Future functional analysis of these SLDs and SBHs will allow to determine their role in the biosynthetic pathway of GSL species in different *E. huxleyi* strains and during viral infection.

**Figure 3:**
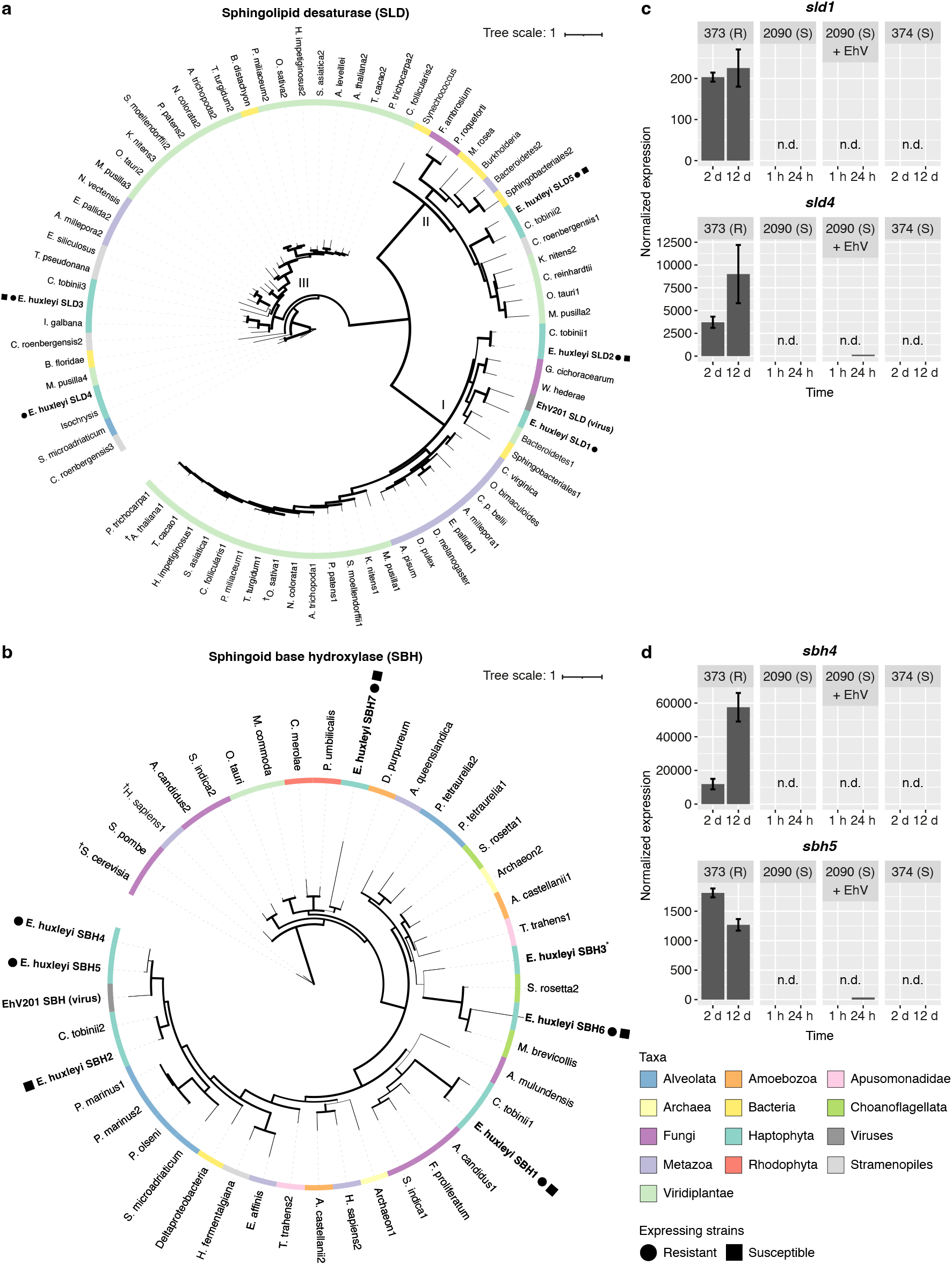
Phylogenetic analysis and gene expression patterns of SLDs and SBHs in resistant and susceptible *E. huxleyi* strains. Phylogenetic trees of (**a**) SLD and (**b**) SBH proteins based on the conserved domains. Protein domain sequences were aligned using Mafft (for SLD) and ClustalW (for SBH). Maximum Likelihood trees (PhyML) are shown. Colors represent different taxonomic groups and shapes indicate the expression in the resistant *E. huxleyi* strains 373 and 379 (circles) and in the susceptible *E. huxleyi* strains 2090 and 374 (rectangles; legend at the bottom right side). ^†^Functionally characterized protein. *Expression of *sbh3* was not detected in the *E. huxleyi* strains and conditions tested. Bootstrap values are represented by the line width. Expression patterns of (**c**) *sld*1 and *sld*4, and (**d**) *sbh*4 and *sbh*5 in the resistant *E. huxleyi* strain 373 and in the susceptible *E. huxleyi* strains 2090 (uninfected and infected cultures) and 374. Values for *E. huxleyi* strain 373 are presented as the mean ± SD (*n* = 2). Expression was not detected in *E. huxleyi* strains 2090 and 374 under the tested conditions.

### Detection of resistant algal cells in an open ocean bloom using lipid biomarkers

Since little is known about resistance to viral infection in algal blooms, we sought to utilize our new resistant metabolic biomarker (resGSL) to assess the occurrence of resistant cells in an oceanic *E. huxleyi* bloom. To that end, biomass samples for lipidomics analysis were collected during the ‘Tara Breizh Bloom’ cruise in the Celtic Sea, capturing the demise phase of an *E. huxleyi* bloom (Fig. 4a) ^50^. The occurrence of hGSL species (Fig. 4b), which are known lipid biomarkers for *E. huxleyi* and are present in all strains^37, 39^, confirmed the presence of *E. huxleyi* cells, as was also visible using scanning electron microscopy^50^. sGSL species, which characterize susceptible strains^38^, were also detected (Fig. 4c), indicating the presence of virus-susceptible *E. huxleyi* cells in the water. We could also detect 374-GSL species (group D, **13-14**) at a similar intensity as the hGSL species (Fig. 4d), indicating that some *E. huxleyi* cells share similarity to the susceptible *E. huxleyi* strain 374. Importantly, we detected resGSL d19:4/h22:2 (**12**) in four out of the five days of sampling (Fig. 4e). This is the first demonstration of the presence of resistant *E. huxleyi* cells during bloom succession of *E. huxleyi*. The occurrence of hGSL, sGSL, 374-GSL and esGSL species during the demise phase of the bloom suggests a complex population composition towards the end of the bloom.

**Figure 4:**
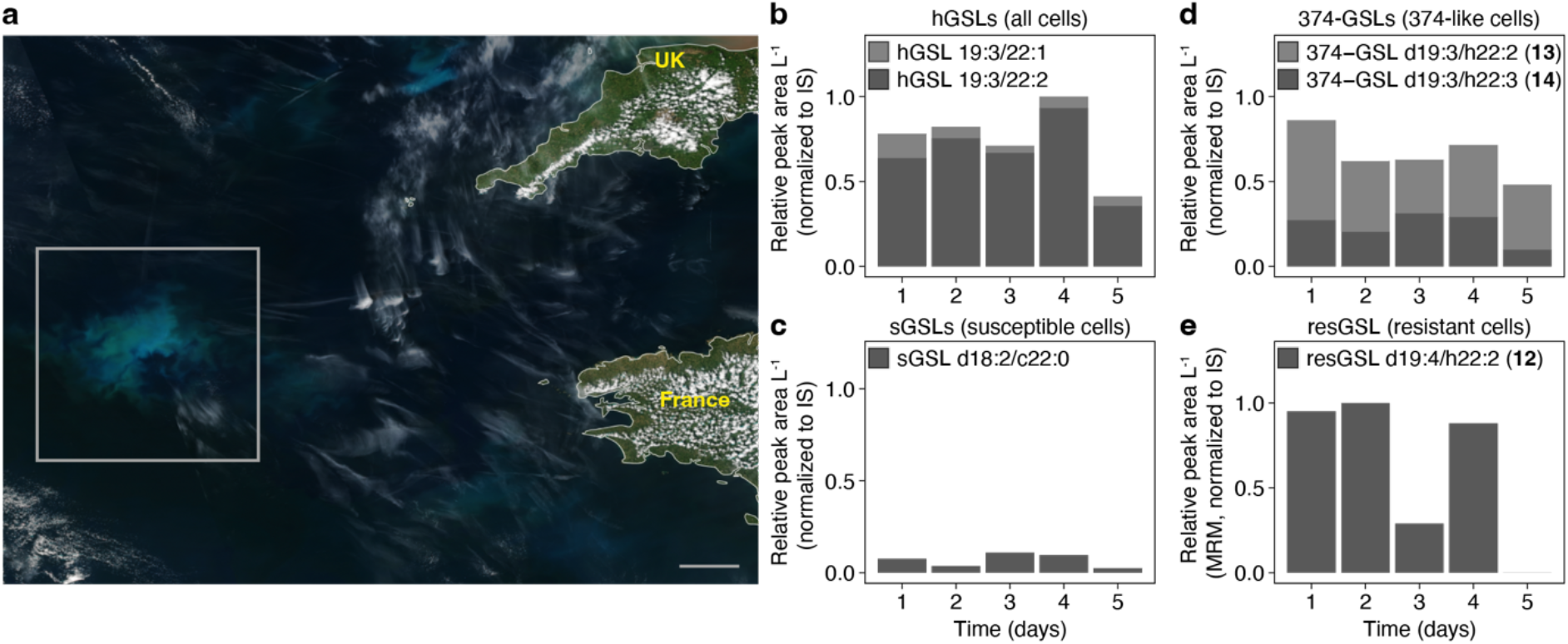
Detection of resGSL in an open ocean *E. huxleyi* bloom. (**a**) Satellite ocean true-color image from the Visible Infrared Imaging Radiometer Suite (VIIRS) onboard the Suomi National Polar-orbiting Partnership (SNPP) depicting the bloom area on May 21, 2019 (marked by a rectangle, source: https://www.star.nesdis.noaa.gov/sod/mecb/color/ocview/ocview.html). Scale bar, 50 km. Relative intensity of (**b**) hGSL (all *E. huxleyi* cells), (**c**) sGSL (susceptible *E. huxleyi* cells) and (**d**) 374-GSL (susceptible, 374-like *E. huxleyi* cells, **13-14**) species during five days of sampling. (**e**) Relative intensity of resGSL d19:4/h22:2 (resistant *E. huxleyi* cells, **12**) analyzed using high-sensitivity multiple reaction monitoring (MRM) mode during five days of sampling.

### Lipidomics profiling during *E. huxleyi* bloom succession and virus-induced demise

Sampling open ocean bloom provides only a snapshot of the bloom dynamics. Therefore, we sought to gain a detailed temporal resolution for our suite of biomarkers in order to assess the various phenotypes the occur during *E. huxleyi* bloom succession. Therefore, we conducted an *in situ* mesocosm experiment in the coastal waters of southern Norway^51, 52^, where annual blooms and viral infection of *E. huxleyi* occur naturally^27^. Briefly, the experiment included seven mesocosm bags that were filled with natural marine microbial communities and monitored daily over 24 days. Four bags (bags 1-4) were sampled for lipidomics analysis and are discussed hereinafter. All bags were supplemented with nutrients at a nitrogen to phosphorous ratio of 16:1 to favor the growth and induce a bloom of *E. huxleyi*^53^. *E. huxleyi* blooms were observed starting from day 10 in all bags, reaching a concentration of up to 8×10^7^ cells per L at day 17, followed by bloom demise starting from day 18 (Fig. 5a). Viral infection varied between the bags, as was visible by measurement of biomass-associated EhV by quantitative PCR (qPCR) using the major capsid protein (*mcp*) gene. Bag 4 showed the strongest increase in EhV starting from day 17, followed by bag 2 and bag 1. No viral proliferation was observed in bag 3 (Fig. 5b). Concomitantly, the extent of bloom demise also varied between the bags, reaching the lowest cell abundance in bag 4 (Fig. 5a).

**Figure 5:**
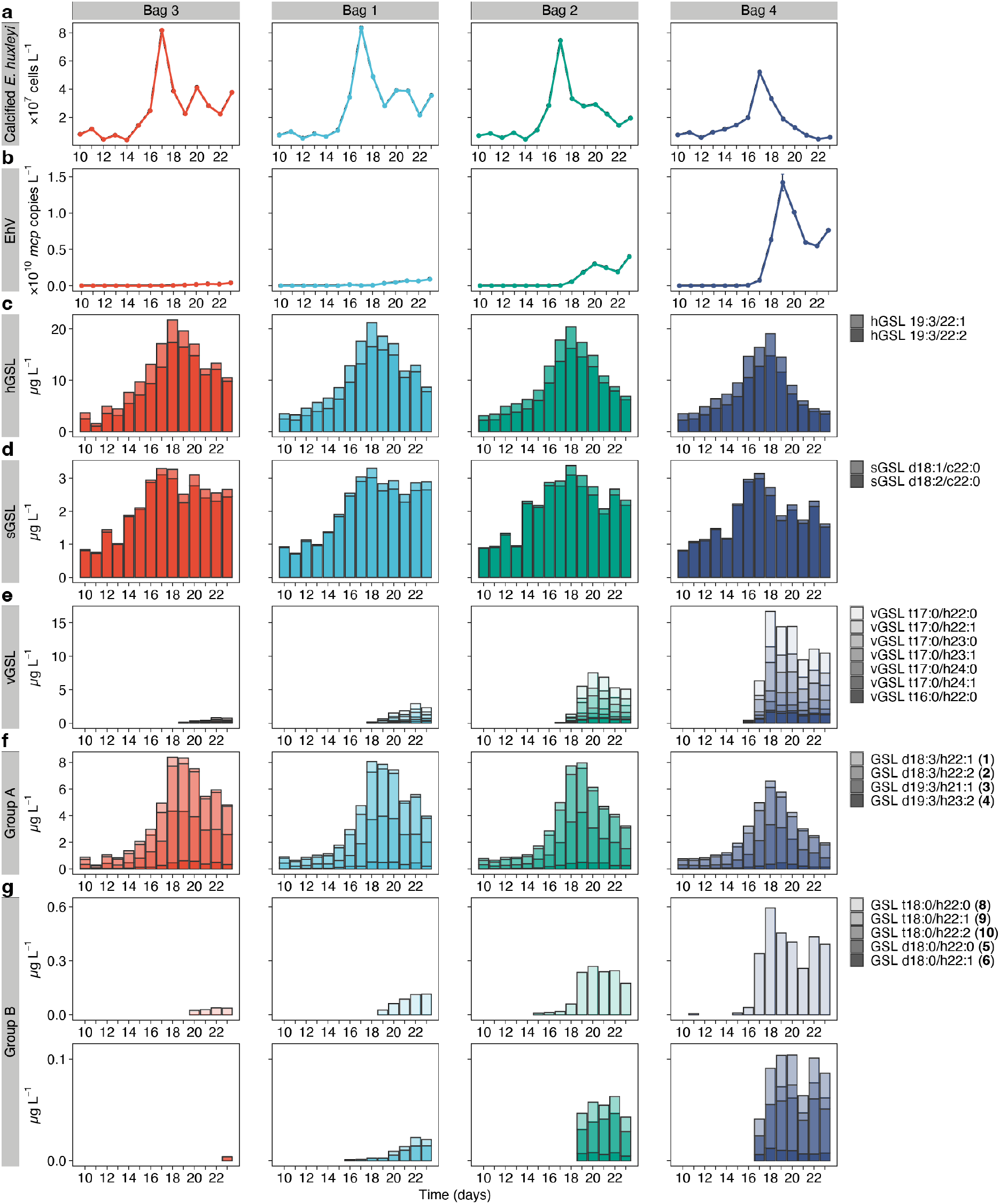
Changes in cellular content of GSL species in response to viral infection of natural *E. huxleyi* populations. Variable growth and infection dynamics of *E. huxleyi* across four mesocosm bags based on (**a**) the abundance of calcified *E. huxleyi* cells and (**b**) biomass-associated EhV, starting from day 10 of the experiment. Bags are ordered by increasing EhV abundance, with the lowest in bag 3 and the highest in bag 4. Abundance of calcified *E. huxleyi* cells is based on flow cytometry analysis and abundance of biomass-associated EhV is based on the quantification of the EhV major capsid protein (*mcp*) gene by qPCR. *mcp* copy values are presented as the mean ± SD (*n* = 3, technical replicates). Concentration of (**c**) hGSL, (**d**) sGSL, (**e**) vGSL, (**f**) group A GSL, and (**g**) group B GSL species are presented. GSL d18:1/h22:21 (**7**) was not detected to due technical reasons. Group B was divided into two rows due to difference in concentrations of the different species.

We followed changes in the lipid composition of the particulate fraction (1.6-25 µm) during the bloom and demise of *E. huxleyi* (days 10-23) by LC-HRMS. Known lipid biomarkers of *E. huxleyi* were used to describe changes that occur during the bloom: hGSL species, present in all *E. huxleyi* strains^37, 39^, correlated with *E. huxleyi* abundance (Fig. 5c, Pearson correlation, *r* = 0.66-0.73, Table S7); sGSL species, which characterize susceptible strains^38^, also correlated with *E. huxleyi* abundance, primarily in the bloom phase (Fig. 5d, *r* = 0.58-0.72, Table S7).

To detect active viral infection of *E. huxleyi* cells, we monitored the production of vGSL species that are produced only by infected cells^36, 37^. Six t17:0-based vGSL species and one t16:0-based vGSL species were positively correlated with the varying degree of infection between the bags, as measured by the abundance of biomass-associated EhV (Fig. 5e, *r* = 0.87-0.95, Table S7). In bag 4, the sum concentration of vGSL species was similar to that of hGSL species (∼15 µg per L and ∼20 µg per L on day 18, respectively), which exemplifies the pronounced metabolic remodeling in infected cells. Moreover, several vGSL species were detected in bag 4 as early as day 16, one day before the first detection of biomass-associated EhV (Fig. 5b) or extracellular EhV^52^. To our surprise, we detected low levels of vGSL species also in bag 3 starting from day 20, although viral abundance (as measured by qPCR, Fig. 5b) was below the detection limit. This indicates that a small number of *E. huxleyi* cells in bag 3 were infected following bloom demise, however, it is not clear whether production of virions or abortive infection occurred. Altogether, these observations suggest that vGSL species can serve as a more sensitive biomarker for the occurrence of infected cells in the natural environment than the quantification of virions by gene biomarkers.

Interestingly, we could not detect resGSL species (**11-12**, group C), which are characteristic of resistant cells, suggesting that the abundance of resistant *E. huxleyi* cells throughout the bloom and demise phases (and within our sampling period) was below the level of detection. GSL species of group A (**1-4**), which are found in higher intensity in resistant strains compared to susceptible strains in the laboratory (Fig. 2a and Fig. S18), appeared from the beginning of the bloom and were highly correlated to hGSL species and to a lesser extent to *E. huxleyi* abundance and sGSL species (Fig. 5f, *r* = 0.76-0.96, 0.53-0.75 and 0.70-0.81, respectively, Table S7). Accordingly, the detection of these GSL species is most probably derived from susceptible cells that dominated the bloom rather than rare resistant cells. 374-GSL species (group D, **13-14**) appeared in a similar pattern to group A (Fig. S23 and Table S7), indicating a high abundance of susceptible cells that share some similarity to *E. huxleyi* strain 374. GSL species of group B (**5-6**, **8-10**), which are found in resistant strains and infected susceptible strains, appeared mostly from day 17 onwards and were highly correlated to the abundance of EhV and of the main vGSL species (vGSL t17:0/h22:0, Fig. 5g, *r* = 0.69-0.99, Table S7), as was also observed in the laboratory (Fig. S18). The amount of these GSL species was ∼20 times lower than that of vGSL species, and might be a result of enzyme promiscuity in infected cells^36^. Nevertheless, the infection-related occurrence of these GSL species, which are characteristic of resistant strains and are also induced during infection in the laboratory (Fig. S18), might suggest an infection-derived initiation of cellular processes that eventually lead to resistance.

### Remodeling of GSL composition and induction of resistance in infected cultures

To assess whether resistant *E. huxleyi* cells appear in low numbers during bloom succession, as detected in the open ocean bloom (Fig. 4e), we isolated numerous *E. huxleyi* clones during the mesocosm experiment and determined their susceptibility to infection by EhV strain M1 (EhVM1), which was isolated during the same mesocosm experiment^54^. Most isolates were found to be susceptible to EhVM1, among them isolates RCC6918, RCC6936 and RCC6912 (Fig. 6a, b and Fig. S24a, b), which were isolated during the bloom phase of *E. huxleyi*^51^. However, we also isolated a few resistant *E. huxleyi* strains, among them isolate RCC6961 (Fig. S24c). This isolate, along with additional resistant isolates, was isolated during the virus-induced demise phase of the *E. huxleyi* bloom^51^. Interestingly, some of the isolated susceptible strains showed rapid recovery 1-2 weeks after viral infection (RCC6918 and RCC6912, Fig. 6b and Fig. S24b, respectively). The recovered populations were resistant to the virus, as was validated by re-exposing the cultures to viral infection (Fig. S25a). To examine whether the newly identified GSL markers for resistant cells can differentiate between the *E. huxleyi* isolates with different phenotypes, we compared the GSL composition of the isolates and the recovered cultures (Fig. 6c, d and Fig. S25b). All isolates had similar amounts of hGSL species. The susceptible mesocosm isolates RCC6936, RCC6918 and RCC6912 had a similar GSL composition to *E. huxleyi* 374, having high intensity of sGSL and 374-GSL species, and a lower intensity of group A GSL species (Fig. 6c, d and Fig. S25b). GSL species from groups B and C were not detected in these susceptible isolates. The resistant isolate RCC6961 had a similar GSL composition to the resistant laboratory strains 373 and 379, with higher intensity of GSL species from group A (compared to the susceptible isolates) and presence of GSL species from groups B and resGSL species from C (Fig. 6c). The distinct occurrence of resGSLs species in a resistant isolate further supports its use as a biomarker for resistant cells. As predicted based on the GSL biomarkers, sGSL species were not detected in this isolate, and, surprisingly, neither were 374-GSL species. Remarkably, the cultures that recovered following infection of isolates RCC6918 and RCC6912 and acquired resistance to the viral infection had a similar GSL composition to the resistant isolate RCC6961 and the resistant laboratory strains 373 and 379, including the presence of resGSLs (Fig. 6d and Fig. S25b). These results indicate a metabolic plasticity in GSL metabolism, which corresponds to the change in phenotype from susceptibility to resistance towards viral infection. Furthermore, the detection of resGSL species (group C) in resistant isolates from the mesocosm and recovered resistant cultures suggests that these GSL species might have been produced during the mesocosm experiment by these rare populations, albeit in concentrations below our detection limit.

**Figure 6:**
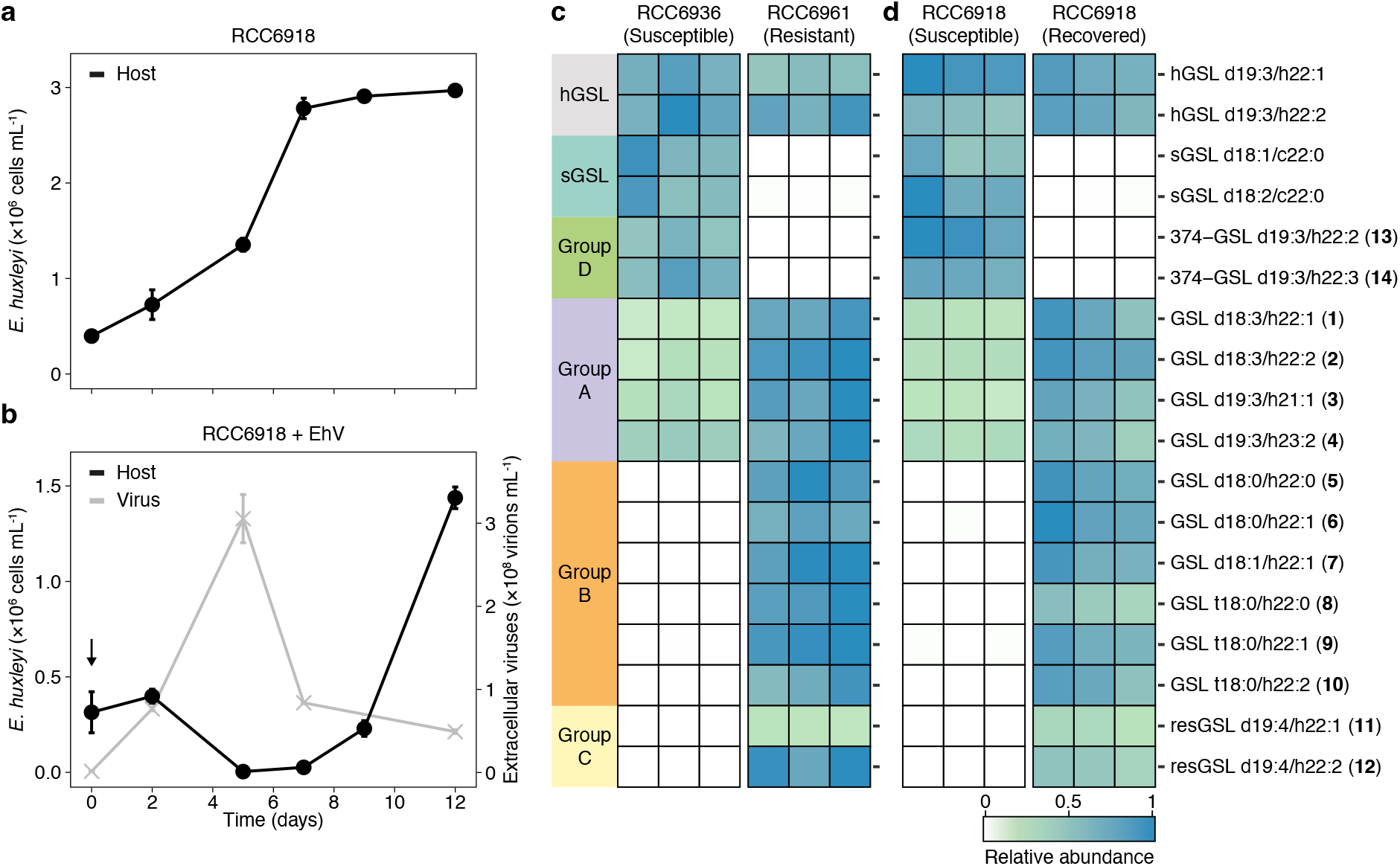
Plasticity in the GSL composition of *E. huxleyi* cultures that recover following viral infection. (**a**) Cell abundance in cultures of the susceptible *E. huxleyi* isolate RCC6918, isolated from the mesocosm experiment. (**b**) Cell abundance (black) and production of virions (grey) following addition of EhVM1 to *E. huxleyi* isolate RCC6918. A recovered resistant population emerged a week after infection. Values of (a) and (**b**) are presented as the mean ± SD (*n* = 2). The black arrow indicates the addition of EhVM1 to the cultures. (**c**) GSL composition of the susceptible *E. huxleyi* isolate RCC6936 and the resistant *E. huxleyi* isolate RCC6961, both isolated from the mesocosm experiment (*n* = 3). (**d**) GSL composition of the susceptible *E. huxleyi* isolate RCC6918, isolated from the mesocosm experiment, and of the culture that recovered following infection and was resistant to the virus (*n* = 3). Values for each lipid species (row) in (c) and (d) are shown after normalization. GSL species are grouped and numbered based on Table 1.

## Discussion

Resistance to viral infection has been described in various phytoplankton cultures under laboratory conditions^15, 16, 55, 56^. Nevertheless, the extent of resistance in natural algal populations is unknown as we lack the tools to detect resistant cells, hindering our ability to understand the metabolic basis of resistance to viral infection and its ecological significance. In the *E. huxleyi*-EhV model system, the difference in susceptibility of *E. huxleyi* strains to viral infection has been previously associated to ploidy level during life cycle changes, as well as to genome and transcriptome variations between the strains^16, 22, 40, 42, 57^. Resistant cells were also identified as a small sub-population in infected cultures^42^. Yet, to date there exists no specific metabolic biomarker for algal resistance to viral infection, and the mechanisms underlying resistance are largely unknown.

### Proposed functional role of resistance-specific LCBs

resGSL species found in resistant cells are characterized by an uncommon tetra-unsaturated LCB 19:4, which has been previously identified only in a few dinoflagellates and other haptophytes (e.g. GSL d19:4/h22:1, which was detected in *Isochrysis galbana*) ^58^. This LCB has an additional double bond compared to the LCB d19:3, which is found in GSL species in *E. huxleyi* (hGSL, group A and 374-GSL species, Table S3), other haptophytes and dinoflagellates, and in SLs of fungi and marine invertebrates^39, 58–61^. Interestingly, resistant *E. huxleyi* strains are also characterized by GSL species containing the trihydroxylated LCB t18:0 (Fig 2b and Table 1), which is highly abundant in plants and fungi^62^ and was thus far found only in vGSL species produced by infected cells (in addition to t16:0-and t17:0-based vGSL species, Table S3) ^36, 38^.

Both LCB unsaturation and hydroxylation were found to affect the biophysical properties of membranes: LCB unsaturation hinders the ability of SLs to form ordered domains within lipid bilayers, known as ‘lipid rafts’^63^, while additional hydroxyl groups facilitate the formation of more hydrogen bonds, leading to an increased stability and decreased permeability of the membrane and to lateral diffusion of membrane proteins^62^. Such changes in SL composition can also initiate signal transduction within the cells, as was found in plants, yeast, and mammals^62, 64^. Subsequently, they allow organisms to cope with environmental stress, such as low temperature^65, 66^, and can alter the susceptibility of cells to viral infection^67–70^. Specifically, GSL-rich lipid rafts in host cells were shown to serve as cellular entry or egress points in diverse systems^71^, suggesting that membrane lipids are under strong selection pressure during host-virus co-evolution, possibly driving the plasticity in lipid composition.

In the *E. huxleyi*-EhV model system, the lipid envelope of EhV and the *E. huxleyi* plasma membrane seem to play an important role at the onset of the infection process, mediating the entry of the virus to *E. huxleyi* cells by endocytosis or membrane fusion mechanism^72^. It was previously suggested that sGSLs, which characterize susceptible cells, mediate viral adsorption to host cells^38^. The occurrence of resGSLs and t18:0-based GSLs in resistant cells, in addition to the absence of sGSLs, might therefore hinder viral adsorption to the cells by impeding membrane fusion. Nevertheless, further structural and biochemical analyses are needed to determine the role of resGSLs and t18:0-based GSLs in modulating the resistance of *E. huxleyi* cells to viral infection.

### Plasticity in GSL composition during *E. huxleyi*-EhV interactions

The lipidome of *E. huxleyi* has been identified as a sensitive metabolic indicator for environmental stress conditions, such as nutrient limitation and viral infection, reflecting the physiological state of the cells^32, 45, 73, 74^. In particular, GSLs were found to play a distinct role in the *E. huxleyi*-EhV system due to their involvement in cell signaling during infection as well as in viral assembly and egress^36–39^. The identification of resGSLs and other GSL species characteristic of resistant *E. huxleyi* cells (Fig 2b and Table 1) broadens our view of the GSL diversity in the *E. huxleyi*-EhV model system and adds valuable biomarkers that were thus far missing (Fig. 7a and Table S3). While hGSL and group A GSL species are shared among all *E. huxleyi* cell types (that is, the different strains and phenotypes), most GSL species are produced only by some: sGSL and 374-GSL species by susceptible cells; vGSL species by infected cells; group B GSL species by both infected and resistant cells; and resGSL species by resistant cells (Table S3). A recent study further found that resistant *E. huxleyi* strains have a more diverse GSL composition than susceptible ones under nutrient-replete conditions^75^. GSL species vary in their sugar headgroup, FA and LCB^76^. In *E. huxleyi*, except for sGSL species that contain a sialic acid headgroup^38^, all other known GSL species contain a hexose-based sugar headgroup (Table S3). Additionally, most species have a highly similar FA composition (with the hydroxylated h22:0, h22:1 and h22:2 FAs being the most common), except for vGSL species that also contain longer FAs of 23-24 carbons, group A GSL species that contain FAs with 21 and 23 carbons (Table 1), and sGSL species that contain non-hydroxylated FAs (c22:0, c22:1, see Table S3) ^38^. LCB composition, on the other hand, seems to be the main factor that differentiates between the various GSL groups and, consequently, between the cell types, thus driving the phenotypic plasticity in the *E. huxleyi*-EhV model system (Fig. 7a and b). Some LCBs are shared among several GSL species and cell types (LCB d18:1, d18:3 and d19:3), while others appear only in specific GSL species and cell types (LCB d18:2 in sGSLs of susceptible cells, LCB d19:4 in resGSLs of resistant cells, LCB d18:0 and t18:0 in group B GSLs of resistant and infected cells, and LCB t16:0 and t17:0 in vGSLs of infected cells), leading to a unique LCB profile for each cell type (Fig. 7b). Biosynthetic genes at various steps of the GSL pathway determine LCB composition, from the formation of the LCB to its hydroxylation and unsaturation^77^. The presence of these genes and their differential expression under various biotic and abiotic conditions determine the GSL composition of the cells^66^. In infected *E. huxleyi* cells, virus-encoded SL biosynthetic enzymes lead to the production of t17:0-based vGSLs^36^. In resistant *E. huxleyi* strains, our results suggest that the differential expression of specific *sld* and *sbh* genes (Fig 6c, Fig. 3c and d) accounts for the biosynthesis of d19:4-based resGSL and t18:0-based GSL species. Remarkably, resistant *E. huxleyi* cells that emerge from infected susceptible cultures as early as one week post infection (Fig. 6b) produce resGSL and group B GSL species that are characteristic of resistant strains, consisting of LCBs that are not found in the parent susceptible strains (Fig. 6c). This striking difference between the parent cells and the derived resistant cultures delineates the plasticity of the *E. huxleyi* lipidome. Such a rapid modulation of GSL composition following viral infection is therefore not restricted only to infected cells but might occur also in cells that evade infection or survive and become resistant to the virus. If so, viral infection might directly induce changes in host LCB biosynthesis and lead to the formation of GSL species that facilitate resistance. Alternatively, resistant cells may already exist as a rare sub-population in cultures of susceptible strains. Such cultures can recover from infection following the death of susceptible cells due to viral infection, which allows the resistant cells to proliferate. The phylogenetic similarity between the enzymes expressed by resistant strains (SLD1, SBH4 and SBH5) and their viral analogues (EhV201 SLD and EhV201 SBH, Fig. 7c, Fig. 3a and b) may further indicate competing biosynthetic pathways that are co-expressed during infection and affect its outcome, shedding light on the ongoing co-evolution between *E. huxleyi* and its virus.

**Figure 7:**
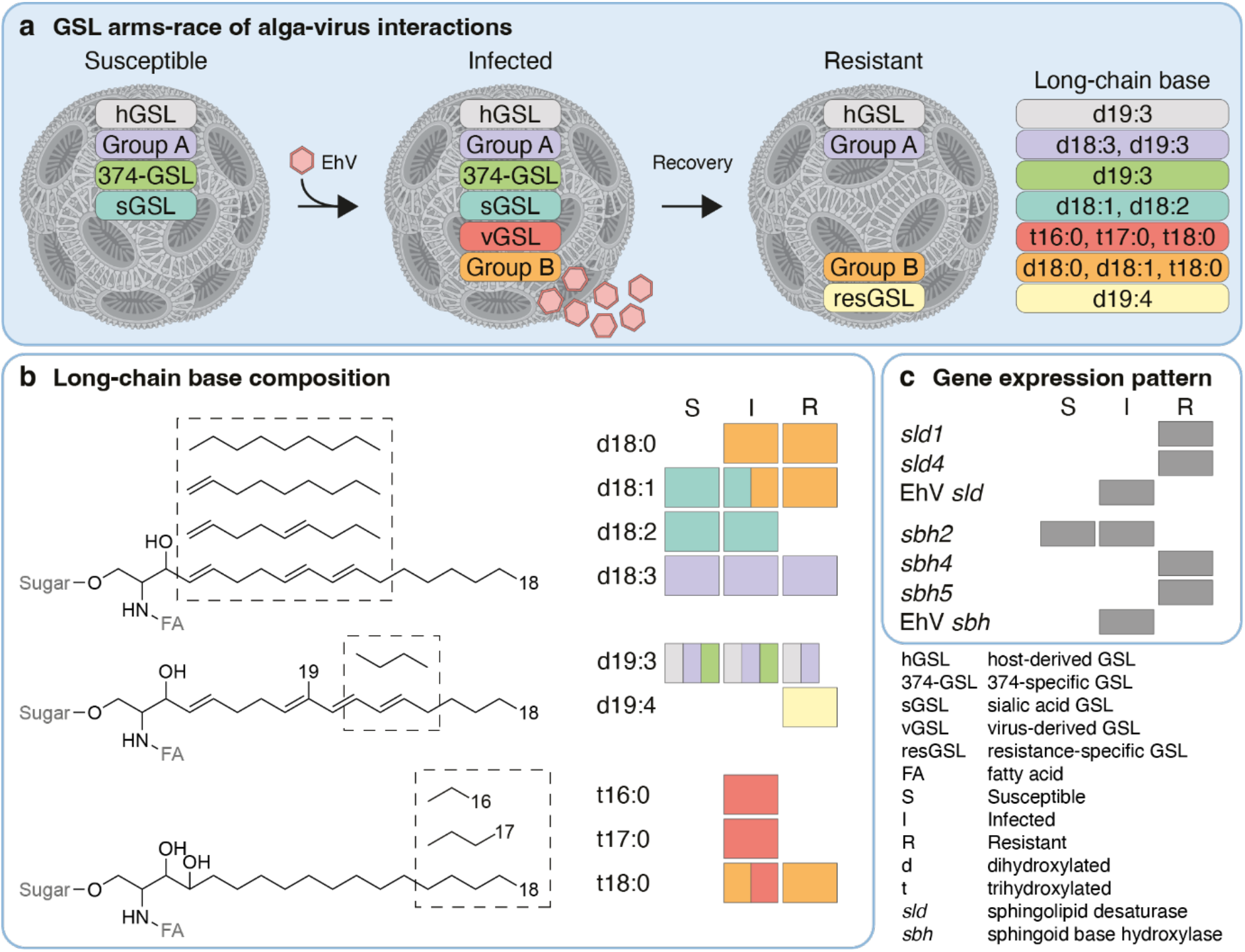
The GSL-based arms race between *E. huxleyi* and EhV. (**a**) The GSL composition of susceptible, infected, and resistant *E. huxleyi* cells. Each GSL group is marked with a different color and consists of different LCBs. Infection by EhV leads to the production of vGSL and group B GSL species, while recovered cells and resistant strains present a unique GSL composition, consisting of group B and resGSL species. Scheme created with BioRender.com. (**b**) LCB composition of GSLs in the *E. huxleyi*-EhV system. Presented are the structure of the different LCBs (left) and the LCB profile of susceptible (S), infected (I) and resistant (R) cells (right). Infected cells produce trihydroxylated LCBs (found in vGSL and group B GSL species), while resistant cells produce both trihydroxylated and tetra-unsaturated LCBs (found in group B and resGSL species, respectively). Colors mark the GSL group in which the LCB is found, as in (a). The position of the double bonds and functional groups were assigned based on the most common structure in the Lipid Maps Structure Database (LMSD) ^44^. (**c**) Expression pattern of *sld* and *sbh* genes which are differentially expressed in susceptible (S), infected (I) and resistant (R) *E. huxleyi* strains. *sld* and *sbh* genes are involved in LCB modification as part of the GSL biosynthetic pathway. EhV *sld* and EhV *sbh* are encoded by the EhV genome.

### Detecting resistant *E. huxleyi* cells in natural populations

Resistance to viral infection has been long studied using model systems in laboratory settings, describing a wide array of *E. huxleyi* strains that vary in their susceptibility to viral infection, and of EhV strains that vary in their level of infectivity^22, 42, 78^. *E. huxleyi* strains can recover from infection and gain resistance to the virus, highlighting their phenotypic plasticity and the rapid change in the dominating phenotypic state in the host cell population^42, 79^. Nevertheless, although we are able to detect susceptible and infected *E. huxleyi* cells in natural samples using GSL biomarkers (Fig. 5d,e) ^38, 39^, we still lack the tools to detect resistant cells in nature and to monitor their dynamics in natural heterogeneous populations.

In this study, we were able to detect resGSL species during the demise of an open ocean *E. huxleyi* bloom (Fig. 4e), indicating, for the first time, the occurrence of virus-resistant *E. huxleyi* cells in natural populations. The absence of resGSLs in samples from the mesocosm experiment stresses the scarcity of resistant *E. huxleyi* cells during the bloom phase and the early phase of the virus-induced bloom demise. This is further supported by the detection of resGSLs in resistant *E. huxleyi* isolates that originate from the mesocosm experiment. Additionally, the emergence of resistant cells 1-2 weeks after viral infection of some susceptible isolates in the laboratory (Fig. 6b) suggests that these cells can be detected during late and post-bloom phases in nature, as observed in the open ocean samples (Fig. 4e). Thus, the sampling time of the mesocosm experiment might not have been long enough to see such an emergence of resistant sub-populations.

In the future, combining the GSL biomarkers for the different cell types with advanced methods, such as single-cell lipid profiling and single-cell RNA sequencing^13, 80, 81^, could allow us to deconstruct the metabolic and phenotypic outcome of viral infection. Studying and identifying the various cell types that constitute algal blooms and the metabolites they use to communicate will provide valuable insights into the host-virus arms race during bloom succession.

## Materials and Methods

### Strains of *E. huxleyi* and EhV used in this study

Four *E. huxleyi* strains were used for the untargeted lipidomics profiling: CCMP2090, CCMP373, CCMP374 and CCMP379 (hereinafter, *E. huxleyi* 2090, 373, 374 and 379), all are non-calcifying. *E. huxleyi* 2090 and 374 are susceptible to viral infection, e.g., by EhV201, while *E. huxleyi* 373 and 379 are resistant. Transcriptomics data of all four strains are publicly available^40, 48, 82^. The *E. huxleyi* cultures were supplemented with the lytic virus EhV201^22^, whose genome data is publicly available^83^. Additionally, four *E. huxleyi* isolates, which were obtained during a mesocosm experiment (see below), were used for a targeted analysis of GSL composition: RCC6912, RCC6918, RCC6936 and RCC6961. Isolates RCC6912, RCC6918, RCC6936 are susceptible to viral infection by EhVM1, while isolate RCC6961 is resistant (Fig. 6b and Fig. S24). The lytic virus EhVM1 was isolated during the same mesocosm experiment and its genome data is publicly available^54^.

To isolate *E. huxleyi* strains from the mesocosm experiment, water samples were collected, and single *E. huxleyi* cells were sorted within two weeks of collection at the Roscoff Culture Collection (RCC) laboratories (https://roscoff-culture-collection.org). *E. huxleyi* RCC6912 and RCC6918 were isolated from bag 1 at day 10 of the experiment (June 3, 2018), during the bloom phase of *E. huxleyi*. *E. huxleyi* RCC6936 was isolated from bag 4 at day 13 of the experiment (June 6, 2018), also during the bloom phase of *E. huxleyi*. The resistant isolate RCC6961 was isolated from bag 7 at day 16 of the experiment (June 9, 2018), during the virus-induced demise of *E. huxleyi*^51^.

### Culture maintenance and viral infection experiments

Cells were cultured in modified K/2 medium (including replacement of organic phosphate with 18 µM KH_2_PO_4_) ^84^ in filtered and autoclaved seawater (FSW) supplemented with ampicillin (100 µg per mL) and kanamycin (50 µg per mL), and incubated at 18°C with a 16:8 h light:dark illumination cycle. A light intensity of 100 μmol photons m^-2^ s^-1^ was provided by cool white light-emitting diode lights. In all infection experiments, EhV was added to the cultures at the exponential phase (5×10^5^ to 1×10^6^ cells per mL) 2 h after the onset of the light period, at a ratio of 5:1 viral particles to *E. huxleyi* cells using a viral lysate derived from an infected *E. huxleyi* 374 culture. Growing cultures in the presence of antibiotics maintained a low basal abundance of bacteria throughout the experiments (Fig. S26). As previously shown, the lipid profile of *E. huxleyi* cultures does not change significantly in the presence of low levels of bacteria^33^.

### Enumeration of algae, virions and bacteria by flow cytometery

Algal cells were quantified using an Eclipse (iCyt) flow cytometer (Sony Biotechnology, Champaign, IL, USA, using ec800 version 1.3.7 software) equipped with a 488 nm solid state-air cooled laser (25 mW on the flow cell) and a standard filter setup. Algal cells were identified by plotting chlorophyll autofluorescence (em: 663-737 nm, see Fig. S27a). Virions and bacteria were quantified by flow cytometry (Fig. S27b), as described previously^36, 85^. Briefly, samples were fixed with a final concentration of 0.5% glutaraldehyde for 30 min at 4°C, plunged into liquid nitrogen, and stored at −80°C until analysis. After thawing, 5 µL of fixed sample were stained with 195 µL SYBR gold (Invitrogen, Paisley, UK) prepared in Tris-EDTA buffer as instructed by the manufacturer (5 μL SYBR gold in 50 mL Tris-EDTA), then incubated for 20 min at 80°C and cooled down to room temperature. Flow cytometric analysis was performed with excitation at 488 nm and emission at 525 nm. A threshold was applied based on the forward scatter signal to reduce the background noise. The gates ‘EhV’ and ‘Bacteria’ were set by comparing to reference samples containing either EhV201 or bacteria.

### Chemicals and internal standards

All solvents and metabolite standards were obtained at high purity. Methanol (Ultra Gradient high-performance liquid chromatography (HPLC) Grade) was purchased from J.T. Baker (Norway). Acetic acid (ULC/MS), acetonitrile (ULC/MS), isopropanol (ULC/MS) and methyl tert-butyl ether (MTBE, HPLC) were purchased from Bio-Lab (Jerusalem, Israel). Ammonium acetate (≥98%, Optima LC/MS) was purchased from Fisher Scientific (Fair Lawn, NJ, USA). Water was purified by a Milli-Q system (resistivity 18.2 MΩ cm at 25°C, TOC < 5 ppb, Merck Millipore, Molsheim, France). For laboratory culture samples, a SL standard mixture containing ten SL species (Cer/Sph Mixture I, Avanti Polar Lipids, Alabaster, AL, USA, LM6002) was used as extraction standard mixture. For mesocosm samples, glycosylceramide (soy) d18:2/C16:0 (>98%, Avanti Polar Lipids, 131304) was used as extraction standard, and isotopically-labeled d9-PC P-36:1 (P-18:0/18:1, >99%, Sigma, 852475C) and d4-palmitic acid (d4-C16:0, 98%, Cambridge Isotope Laboratories, DLM-2893) were used as injection standards for ultra-performance LC-HRMS (UPLC-HRMS) analysis.

### Extraction of cellular lipids in *E. huxleyi* cultures

Cultures of *E. huxleyi* strains 2090, 373, 374 and 379 with and without addition of EhV were analyzed for cellular lipid composition at days 0, 1, 2 and 3 of the experiment in three biological replicates. At day 0, samples were collected 4 hours after the addition of EhV. The samples (30-150 mL of each culture, equivalent to ∼5×10^7^ cells per sample) were collected by vacuum filtration onto glass microfiber filters (grade GF/C, 47 mm in diameter, pre-combusted at 460°C for >8 h, GE Healthcare Whatman, Buckinghamshire, UK), immediately plunged into liquid nitrogen, and stored at −80°C until extraction^32^. In total, 96 biological samples were collected.

Lipid extraction was performed as previously described^86^ with slight modifications. Briefly, biological triplicates were divided into three batches, with 32 samples in each batch. filters were placed in 15 mL glass tubes and extracted with 3 mL of a pre-cooled (−20°C) methanol:MTBE (1:3, ν:ν) solution containing sphingolipid standard mixture (∼150 nM of each species). The samples were shaken for 30 min at 4°C and sonicated for 30 min. The samples were then supplemented with 1.5 mL water:methanol (3:1, ν:ν) solution, vortexed for 1 min, and centrifuged for 10 min at 3,200×g and 4°C. The upper organic phase (1.5 mL) was transferred to 2 mL centrifuge tubes and dried under a flow of nitrogen (TurboVap LV, Biotage, Uppsala, Sweden). The polar phase was re-extracted with 1.5 mL of MTBE. The upper organic phase (2.25 mL) was combined with the organic phase from the first extraction and dried under a flow of nitrogen (TurboVap LV). The samples were stored at −80°C until UPLC-HRMS analysis. Two extraction blanks were collected following the same procedure using blank filters.

An additional analysis was performed to quantify several GSL species with higher sensitivity (Fig. S19). The samples (250 mL of each culture at the exponential phase, 1-1.5×10^6^ cells per mL, equivalent to ∼4×10^8^ cells per sample) were extracted as described above.

### Untargeted lipid profiling using UPLC-HRMS

Per batch, samples were thawed, re-dissolved in 300 μL mobile phase B (see below), vortexed, sonicated for 10 min and centrifuged at 20,800×g for 10 min at 10°C. The supernatants were transferred to 200 μL glass inserts in autosampler vials and directly used for LC-MS analysis. A pooled quality control (QC) sample was generated by combining aliquots of 10 μL from all biological samples. An aliquot of 1 µL was analyzed using UPLC coupled to a photodiode detector (ACQUITY UPLC I-Class, Waters, Milford, MA, USA) and a quadrupole time-of-flight (QToF) mass spectrometer (SYNAPT G2 HDMS, Waters), as described previously^86^ with slight modifications. Briefly, the chromatographic separation was performed on an ACQUITY UPLC BEH C8 column (2.1×100 mm, i.d., 1.7 μm, Waters). Mobile phase A consisted of water with 1% 1 M ammonium acetate and 0.1% acetic acid. Mobile phase B consisted of acetonitrile:isopropanol (7:3) with 1% 1 M ammonium acetate and 0.1% acetic acid. The column was maintained at 40°C and the flow rate of the mobile phase was 0.4 mL per min. The chromatographic gradient was set as follows: 1 min 45% mobile phase A, linear decrease from 45% to 35% mobile phase A over 3 min, from 35% to 11% mobile phase A over 8 min and from 11% to 0% mobile phase A over 3 min, after which the column was first washed with 100% mobile phase B for 4 min and then returned to initial conditions over 0.5 min and equilibrated for 2.5 min (22 min total run time).

The PDA detector was set to 210-800 nm. A divert valve (Rheodyne) excluded 0-1 min and 20-22 min from injection to the mass spectrometer. The ESI source was set to 120°C source and 400°C desolvation temperature, 1.0 kV capillary voltage, and 40 eV cone voltage, using nitrogen as desolvation gas (800 L/h) and cone gas (20 L/h). The mass spectrometer was operated in full scan MS^E^ resolution position ionization mode (25,000 at *m*/*z* 556) over a mass range of 50-1200 Da alternating with 0.1 min scan time between low-(1 eV collision energy) and high-energy scan function (collision energy ramp of 15-35 eV). Leucine-enkephalin was used as lock-mass reference standard. Pooled QC samples were injected at the beginning, middle and end of each batch.

### Comparative analysis of untargeted lipid profiling data

Raw LC-MS files were converted from the vendor’s format to the open-format ‘netCDF’ using a ‘DataBridge’ (MassLynx version 4.1). Pre-processing of the CDF files was done using the R^87^ packages ‘xcms’^88^ and ‘CAMERA’^89^ obtained from the Bioconductor repository (www.bioconductor.org). This yielded a matrix of 12,190 aligned mass features across samples with corresponding peak intensity values. Parameters for mass feature detection, smoothing, alignment, binning and filtering were set according to the instrument’s mass measurement specifications and detailed manual inspection of known mass features in the raw data, as suggested by the software guidelines (Table S8 and Table S9). The feature matrix was normalized to the total ion current (TIC, per sample) and standardized. The elbow method was applied to determine the number of clusters for a subset of samples (without addition of EhV, 48 samples, *k* = 4, Fig. S1a) and for the whole dataset (with and without addition of EhV, 96 samples, *k* = 4, Fig. S1b), followed by *k*-Means clustering and PCA analysis using the R package ‘factoextra’^90^ for both the subset and the whole dataset. *k*-Means clustering (*k* = 5) and PCA analysis were performed also for the whole dataset with the pooled QC samples, resulting in the same separation to clusters, while the pooled QC samples were grouped together in a fifth cluster (Fig. S28).

Comparative analysis between clusters 3, 4 (containing samples of resistant strains) and clusters 1, 2 (containing samples of susceptible strains) in the subset without addition of EhV (Fig. 1c) was performed by one-way ANOVA with FDR-correction (*p* < 0.01) using the R package ‘qvalue’^91^, reducing the data to approximately 10,922 mass features. The mean intensity of each mass feature was then calculated for all clusters, followed by calculation of the fold change between the cluster with the maximum mean intensity and the other clusters. A fold change of > 20 between the first and third highest clusters was selected, yielding a list of 173 differential mass features, which underwent further manual annotation to obtain a smaller number of feature groups. The peak shape of the extracted ion chromatograms (EICs) from co-eluting mass features was compared using MassLynx (Version 4.1, Waters), and isotopes, adducts and apparent neutral losses (e.g. of water) were annotated, grouping the mass features into 43 feature groups (Table S1). Next, the adduct ion with the highest intensity was selected for each feature group and the corresponding peak area was extracted using MassLynx and QuanLynx (Version 4.1, Waters) across samples in the full dataset (that is, with and without addition of EhV). Peak areas above a signal-to-noise threshold of 10 (limit of quantification) were normalized to the TIC. Per feature, zero values were replaced with half of the minimal value. Hierarchical cluster analysis was then applied to the whole data set (Fig. S2) and to the dataset after averaging the peak areas of the biological replicates (Fig. 2a) following log-transformation using Matlab R2021a, with row-wise (per feature) scaling, row-and column-wise clustering using the default ‘Eucledian’ method and the ‘redblue’ colour panel. Out of the 43 feature groups, nine were putatively identified as GSLs following manual annotation, based on the accurate mass, adducts and apparent in-source fragments.

### Putative annotation and phenotypic grouping of GSL species

The annotation of GSL species that were previously undescribed in the *E. huxleyi*-EhV model system (listed in Table 1) was based on LC-MS/MS analysis and the Lipid Maps computationally-generated database of lipid classes and the Lipid Maps Structure Database (LMSD) ^44^, and carried out according to the Metabolomics Standards Initiative, ‘Level 2 – putatively annotated compounds’^46^. The annotation of previously described GSL species was performed according to the accurate mass and LC-MS/MS fragmentation pattern^36, 38, 45, 52^. LC-MS/MS analyses were performed in positive ionization mode for the protonated molecules using a collision energy ramp of 10-45 eV and a scan time of 0.5 sec. Analyses were performed on samples with high intensities, with injection volumes of 3-5 µL. The data were analysed and processed with MassLynx and QuanLynx (version 4.1, Waters). For MS/MS spectra and a list of fragments of the GSL species that were previously undescribed in the *E. huxleyi*-EhV model system, see Fig. S4-S17.

Quantification and phenotypic grouping of the GSL species (Table 1) was based on their abundance profiles in the different *E. huxleyi* strains in the presence and absence of EhV (Fig. S18 and Fig. S19). The abundance profiles were generated by normalizing the peak area of each GSL species (extracted as described above) to the extraction standard (glucosylceramide d18:1/c12:0) and to the total number of extracted cells. Differences in GSL abundance were tested for day 2 of the experiment by a one-way ANOVA followed by Tukey’s post-hoc test, *p* < 0.01 (Table S10 and Table S11). Day 2 was chosen since it was the first time point in which infected samples appeared as a separate cluster in the *k-*means clustering analysis (Fig. 1d).

### Definition of *sld* and *sbh* genes

The *sld* and *sbh* genes were predicted from *E. huxleyi* genome sequences^82^, and defined based on expressed sequences when available: expressed sequence tags (ESTs) and Illumina short read sequences^40^, as described in ^92^.

Five SLDs and seven SBHs and were defined. All SLDs have the fatty acid desaturase domain (PF00487), while the SBHs have the fatty acid hydroxylase domain (PF04116). *sld2* (KJ868223, called Dcd2) and *sbh1* (KJ868226, called sphingainine hydroxylase 1) were deposited in GenBank from our earlier definitions^31^. The genes were redefined based on PacBio RNASeq long read sequences, and *sld1*-*sld5* and *sbh1*, *sbh2*, *sbh4*-*sbh7* sequences from the susceptible strain *E. huxleyi* 2090 and the resistant strain *E. huxleyi* 373 (either one, the other or both strains, depending on expression) were deposited in GenBank, and given accession numbers: MZ152812-MZ152827 (Table S4). *sbh*3 was not expressed in any condition checked, and therefore was not submitted to GenBank. The sequence is available, with all others used for the phylogenetic analysis, in Figshare: 10.6084/m9.figshare.20448579. Expression patterns of *sld* and *sbh* genes were based on data from previously published studies^31, 40, 48^.

### Phylogenetic analysis of SLD and SBH

Database searches were performed to find similar proteins using BlastP at NCBI^93^ against the nr database, allowing 250 hits (to find more distantly related sequences). As the *E. huxleyi* SLD and SBH proteins differed greatly from each other within each protein family, sequences to represent each branch of the families were chosen, to give as wide an evolutionary spread as possible, while trying to keep consistency in the choice of species (if a species had hits to multiple family members, it was preferred, though species that only matched individual branches were also chosen, Table S5 and Table S6). All sequences were required to have the domains that define the family. Domain searches were performed using the Pfam (http://pfam.xfam.org/) and CDD (https://www.ncbi.nlm.nih.gov/Structure/bwrpsb/bwrpsb.cgi) databases^94, 95^ Multiple alignments were performed on both whole protein and domain only, using ClustalW2.1^96^, Muscle 3.8.31^97^ in local installations, and in the case of SLDs, Mafft V7 online (https://mafft.cbrc.jp/alignment/server/)^98^. Due to the differences in the overall lengths of the proteins, the alignments chosen for the final phylogenetic analyses are those of the domains, as found in CDD. For SLD, as the subfamilies differed strongly even within the domain, Mafft using the L-INS-i algorithm gave the best alignment (Fig. S29). For the SBHs, the alignments were very similar, and the ClustalW alignment was used (Fig. S30). Phylogenetic analysis was performed with ClustalW (Neighbor-joining) and Phylip 3.697 (ProML, maximum likelihood)^99^ in local installations and PhyML 3.0 (maximum likelihood) online (http://www.atgc-montpellier.fr/phyml/)^100^. The topologies were similar, and the PhyML trees are shown. Trees were visualized with iTol (https://itol.embl.de/)^101^. Details of the amino acid sequences are listed in Table S5 (SLD) and Table S6 (SBH). Full sequences, domain sequences and alignments are available in Figshare: 10.6084/m9.figshare.20448579.

### Extraction and lipid profiling of a *E. huxleyi* bloom in the Celtic Sea

Water samples of a natural *E. huxleyi* bloom were collected during the ‘Tara Breizh Bloom’ cruise in the Celtic Sea from May 29 to June 2, 2019^50^. Water samples of 50 L were first filtered through a 20 μm nylon net to remove large particles. Cells were then collected by vacuum filtration onto glass microfiber filters (grade GF/C, 125 mm in diameter, pre-combusted at 460°C for > 5 h, GE Healthcare Whatman). The filters were transferred to 50 mL centrifuge tubes (Sarstedt, Nümbrecht, Germany) and immediately plunged into liquid nitrogen. The filters were kept at −80°C, freeze-dried (Gamma 2-16 LSCplus, Martin Christ, Osterode am Harz, Germany) within 6 months after collection, and stored at −80°C until further processing. Lipid extraction was performed as described above, using different solution volumes: 20 mL of the pre-cooled (−20°C) methanol:MTBE (1:3, ν:ν) solution containing sphingolipid standard mixture (∼150 nM of each species), 15 mL of water:methanol (3:1, ν:ν) solution, and 11 mL of MTBE for re-extraction. The upper organic phase (11 mL for the first extraction, 15 mL for the second extraction) was dried under a flow of nitrogen (TurboVap LV). An extraction blank was collected following the same procedure using a blank filter.

Untargeted profiling of lipids using UPLC-HRMS was performed as described above. An aliquot of 2 µL was analyzed using LC-HRMS as described above. The chromatographic separation was performed on an ACQUITY UPLC BEH C8 column (2.1×100 mm, i.d., 1.7 μm, Waters) attached to a VanGuard pre-column (5 × 2.1 mm, 1.7 µm; Waters). Mobile phase A consisted of water:acetonitrile:isopropanol (4.50:3.85:1.65, ν:ν) with 1% 1 M ammonium acetate and 0.1% acetic acid. Mobile phase B consisted of acetonitrile:isopropanol (7:3, ν:ν) with 1% 1 M ammonium acetate and 0.1% acetic acid. The column was maintained at 40°C and the flow rate of the mobile phase was 0.4 mL per min. The chromatographic gradient was set as follows: 1 min 100% mobile phase A, linear from 100% to 25% mobile phase A over 11 min and from 25% to 0% mobile phase A over 3 min, after which the column was first washed with 100% mobile phase B for 6 min and then returned to initial conditions (100% mobile phase A) over 0.5 min and equilibrated for 3.5 min (25 min total run time). The mass spectrometer was operated as described above over a mass range of 50-1500 Da.

Identification of GSL species was based on characteristic neutral losses and fragments of LCBs and amino FAs following collision-induced dissociation in MS^E^ mode. Relative intensity of hGSL, sGSL and 374-GSL species was performed by extracting the peak area of the adduct ion with the highest intensity or an indicative fragment ion ([M+Na]^+^ for hGSL, [M+H-(Sialic acid-H)-H_2_O]^+^ for sGSL and [Amino FA+H-H_2_O]^+^ for 374-GSL species) following collision-induced dissociation in MS^E^ mode using QuanLynx. LC-MS/MS operating in multiple reaction monitoring (MRM) mode was used to quantify resGSL d19:4/h22:2 (**12**) with increased sensitivity. Data was acquired as described above using the MRM mode, incorporating the observed retention times and accurate masses of precursor and product ions, using a collision energy of 20 eV. The most intense product ion of the d19:4 LCB (*m*/*z* 272.2378) was selected for target enhancement and the product ion of the h22:2 amino FA (*m*/*z* 334.3110) was used for quantification. Peak areas above a signal-to-noise threshold of 10 (limit of quantification) were normalized to the internal standard and the filtered volume.

### Mesocosm experimental setup

A mesocosm experiment (AQUACOSM VIMS-Ehux) was carried out over 24 days (May 24 – June 17, 2018) in Raunefjorden at the University of Bergen’s Marine Biological Station Espegrend, Norway (60.38° N; 5.28° E), as previously described^51, 52^. Briefly, the experiment consisted of seven enclosure bags made of transparent polyethylene (11 m^3^, 4 m deep and 2 m wide, 90% photosynthetically active radiation) mounted on floating frames and moored to a raft stationed in the fjord. Each bag was filled with surrounding fjord water and supplemented with nutrients. Samples for flow cytometric counts were taken twice a day, in the morning (7 am) and evening (8-9 pm) using 50 mL centrifugal tubes and following filtration using a 40 µm cell strainer. Calcified *E. huxleyi* cells were enumerated using an Eclipse iCyt flow cytometer (Sony Biotechnology).

### Enumeration of biomass-associated EhV in the environment by qPCR

Water samples (1-2 L) were sequentially filtered by vacuum through hydrophilic polycarbonate filters with a pore size of 20 µm (47 mm; Sterlitech, Kent, WA, US) and then 2 µm (Isopore, 47 mm; Merck Millipore, Cork, Ireland). Filters were immediately plunged into liquid nitrogen and stored at −80°C until further processing. DNA was extracted from the 2 µm filters using the DNeasy PowerWater kit (QIAGEN, Hilden, Germany) according to the manufacturer’s instructions. Each DNA extract was diluted 100 times, and 1 µL was then used for qPCR analysis as described in ^52^. Briefly, EhV abundance was determined for the major capsid protein (*mcp*) gene^102^: 5′-acgcaccctcaatgtatggaagg-3′ (mcp1Fw^47^) and 5′-rtscrgccaactcagcagtcgt-3′ (mcp94Rv^52^). Results were calibrated against serial dilutions of EhV201 DNA at known concentrations, enabling exact enumeration of viruses. Data is available in ^103^.

### Sampling, extraction, and cellular lipid profiling of mesocosm samples

Water samples for cellular lipidomics analysis were collected daily from bags 1-4, as described previously^52^. Briefly, samples were collected daily at 7-8.30 am using 10 L carboys (pre-cleaned with 1% HCl for > 10 min and rinsed three times with tap water) using a peristaltic pump at a speed of ca. 5 L per min. The samples were pumped through a 200 µm pore-size Nitex nylon mesh screen to remove microzooplankton grazers and large particles. Carboys were kept at 10°C and processed < 1 h after collection.

Water samples of 1-6 L (depending on the biomass) were first gravity filtered through 25 µm pore-size stainless steel filters (47 mm in diameter, Sinun Tech) to remove large particles. Cells were then collected by gentle vacuum filtration of 1-2 L onto glass microfiber filters (grade GF/A, 47 mm in diameter, pre-combusted at 460°C for > 5 h, GE Healthcare Whatman). The filters were transferred to 2.0 mL centrifuge tubes (SafeLock, Eppendorf) using stainless steel tweezers (pre-combusted at 460°C for > 5 h), supplemented with 5 µL of glycosylceramide d18:2/C16:0 (3 µg per µL in chloroform:methanol, 1:1 ν:ν) and immediately plunged into liquid nitrogen. An extraction blank was taken by soaking a glass microfiber filter in FSW, after which it was transferred to a 2.0 mL centrifuge tube and immediately plunged into liquid nitrogen. The filters were kept at −80°C, freeze-dried (Gamma 2-16 LSCplus, Martin Christ) within 1.5 months after collection, and stored at −80°C until further processing. Lipid extraction was performed for bag samples in days 10-23 (56 biological samples in total) and for the extraction blank as described for the laboratory samples, without the addition of a sphingolipid internal standard mixture and including two solvent blanks.

Untargeted profiling of lipids using UPLC-HRMS was performed as described above, with the following modifications: samples were randomized and divided into two batches with 30 samples in each batch, including extraction and solvent blanks. Randomization was performed automatically using an in-house R^87^ script with the following constraints: every bag was equally represented in each analytical batch and each experimental sampling day was represented at least once. Per batch, samples were thawed, re-dissolved in 220 μL mobile phase B containing d9-PC P-36:1 (0.5 µg per mL) and d4-palmitic acid (0.7 µg per mL) as injection standards, vortexed, sonicated for 10 min, and centrifuged at 20,800×g for 10 min at 10°C. The supernatants were transferred to 200 μL glass inserts in autosampler vials and directly used for LC-MS analysis. A pooled QC sample was generated by combining aliquots of 10 μL from all biological samples. An aliquot of 1 µL was analyzed using LC-HRMS as described above. The chromatographic separation was performed as described above for the Celtic Sea samples.

Identification of GSL species was performed as described above using LC-MS/MS analyses (see Fig. S31-S33 for fragmentation patterns of representative hGSL, sGSL and vGSL species). Absolute quantification of most GSL species was performed by extracting the peak area of the adduct ion with the highest intensity or an indicative fragment ion ([M+Na]^+^ for most GSL species, [M+H-(Sialic acid-H)-H_2_O]^+^ for sGSL and [Amino FA+H-H_2_O]^+^ for 374-GSL species) following collision-induced dissociation in MS^E^ mode using QuanLynx. LC-MS/MS operating in MRM mode was used to quantify group B GSL species with increased sensitivity. Data was acquired as described above using the MRM mode, incorporating the observed retention times and accurate masses of precursor and product ions, using a collision energy of 20 eV. The most intense product ion of the LCB (*m*/*z* 284.2953 for d18:0-based GSL species and *m*/*z* 300.2903 for t18:0-based GSL species) was selected for target enhancement and used for quantification. Peak areas above a signal-to-noise threshold of 10 (limit of quantification) were normalized to the internal standard and the filtered volume.

### GSL profiling of naïve and recovered cultures of the mesocosm-derived *E. huxleyi* isolates

EhVM1 was added to cultures of *E. huxleyi* isolates RCC6912, RCC6918, RCC6936 and RCC6961 at a ratio of 1:1 viral particles to *E. huxleyi* cells, as described above. Of the three susceptible isolates, isolates RCC6912 and RCC6918 recovered about a week following infection (Fig. 6b and Fig. S24b). The recovered cultures were continuously refreshed in modified K/2 medium, until no EhV was detected using flow cytometry.

*E. huxleyi* cultures of mesocosm isolates were analyzed for cellular lipid content at the exponential phase in three biological replicates (1×10^6^ to 2.5×10^6^ cells per mL, see Fig. S25c). The samples (100-150 mL of each culture, equivalent to ∼2×10^8^ cells per sample) were collected by gentle vacuum filtration onto glass microfiber filters (grade GF/A, 47 mm in diameter, pre-combusted at 460°C for > 5 h, GE Healthcare Whatman), immediately plunged into liquid nitrogen, and stored at −80°C until extraction. In total, 18 biological samples were collected. Lipid extraction was performed as described for the laboratory strains, including three extractions blanks. Untargeted profiling of lipids using UPLC-HRMS was performed as described for the mesocosm samples, using 200 μL mobile phase B for re-dissolving samples.

The samples were injected in one batch, including an extraction blank and a pooled QC sample. Absolute quantification of GSL species was performed as described for the laboratory strains, by normalizing the peak area of each GSL species to the extraction standard (glucosylceramide d18:1/c12:0) and to the total number of extracted cells. Heatmaps were generated using R^87^ with column-wise (per GSL species) normalization and the ‘GnBu’ color panel of the package ‘RcolorBrewer’ (Fig. 6c,d and Fig. S25b).

## Acknowledgements

We thank all team members of the AQUACOSM VIMS-Ehux project for setting up and conducting the mescosom experiment, especially Jorun Egge, Aud Larsen, Tatiana Tsagaraki, Celia Marrasé and Rafel Simó. We thank Ian Probert and Martin Gachenot for isolating and maintaining the *E. huxleyi* strains during the mesocosm experiment. We are grateful to the Tara Ocean Foundation, led by Romain Troublé and Etienne Bourgois, for the sampling opportunity and facilities onboard Tara, and to all the scientific and logistic team involved in the Tara Breizh Bloom cruise, notably captain Martin Herteau and his crew, the chief scientist Christian Jeanthon, and Colomban de Vargas. We are grateful to Ilana Rogachev for technical support on the LC-MS instrument. We thank Roi Avraham, Noa Ben-Moshe and Ron Rotkopf for their help in data analysis.

## Author contributions

G.S., C.Z. and A.V. conceptualized the project. G.S., C.K. and A.V. wrote the manuscript. G.S., C.K., C.Z. and S.M. designed and performed the experiments. G.S., C.K. and C.Z. performed the LC-MS data analysis. S.B.-D and G.S. performed the bioinformatics analyses. D.S. collected the open ocean samples. All authors reviewed and edited the manuscript.

## Data availability

Data supporting the findings of this study are available in the paper and its Supplementary Materials. Flow cytometry, qPCR, nutrient and temperature data from the mesocosm experiment are available in Dryad (https://doi.org/10.5061/dryad.q573n5tfr). Mass spectral raw data was deposited to the EMBL-EBI MetaboLights repository with the identifier MTBLS3323 (www.ebi.ac.uk/metabolights/MTBLS3323). Raw data files of biological samples from the laboratory, ‘Tara Breizh Bloom’ cruise and mesocosm experiments include full MS and MS/MS analyses in positive ionization mode. Nucleotide sequences were deposited in GenBank and given accession numbers: MZ152812-MZ152827. Full sequences, domain sequences and alignments used for the phylogenetic analysis are available on Figshare: 10.6084/m9.figshare.20448579.

## Supplementary Information

**Figure S8:**
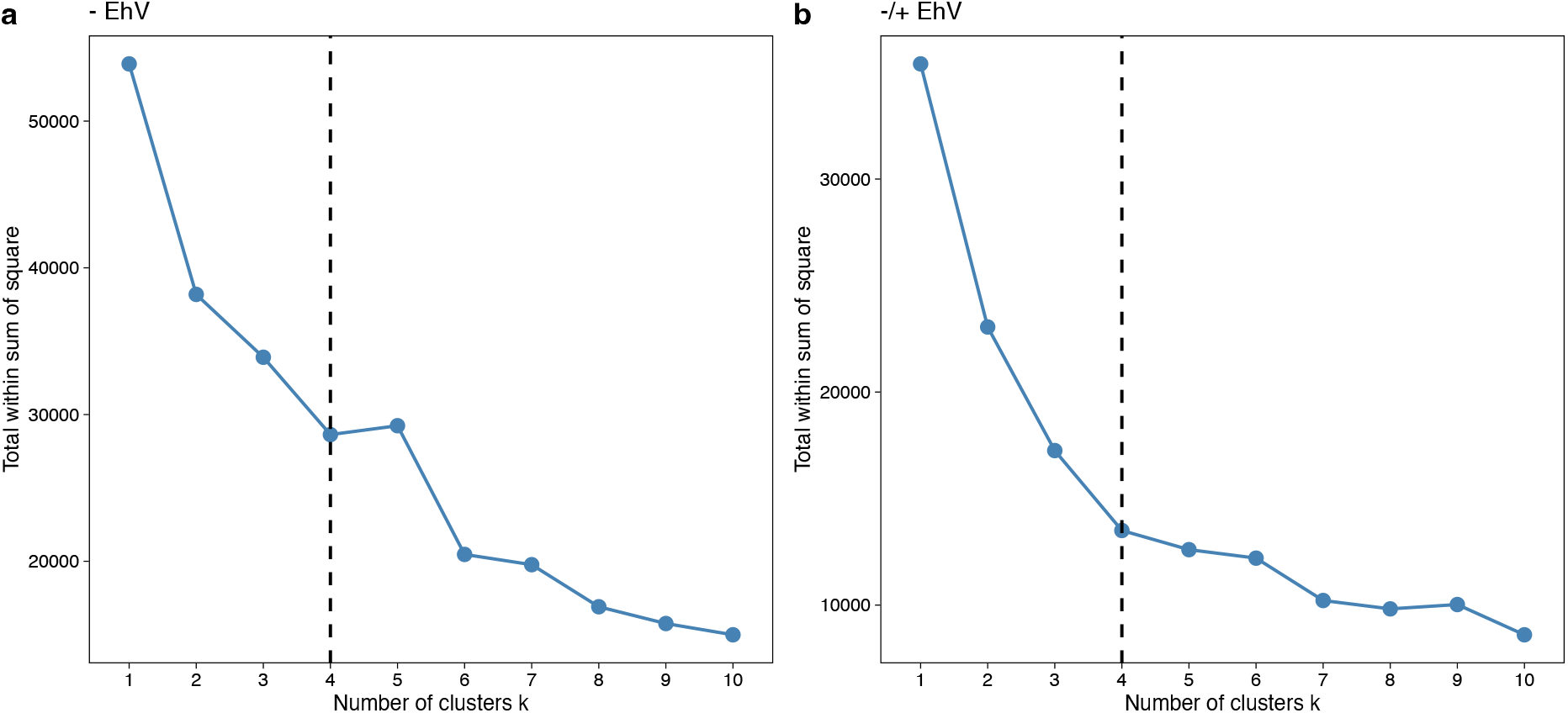
Elbow method for determining the best number of clusters. The method was applied on untargeted LC-MS-based lipidomics data (using 12,190 mass features) derived from cultures of two resistant and two susceptible *E. huxleyi* strains (**a**) without addition of EhV and (**b**) with and without addition of EhV. In both cases, *k* = 4 was chosen as the best number of clusters.

**Figure S9:**
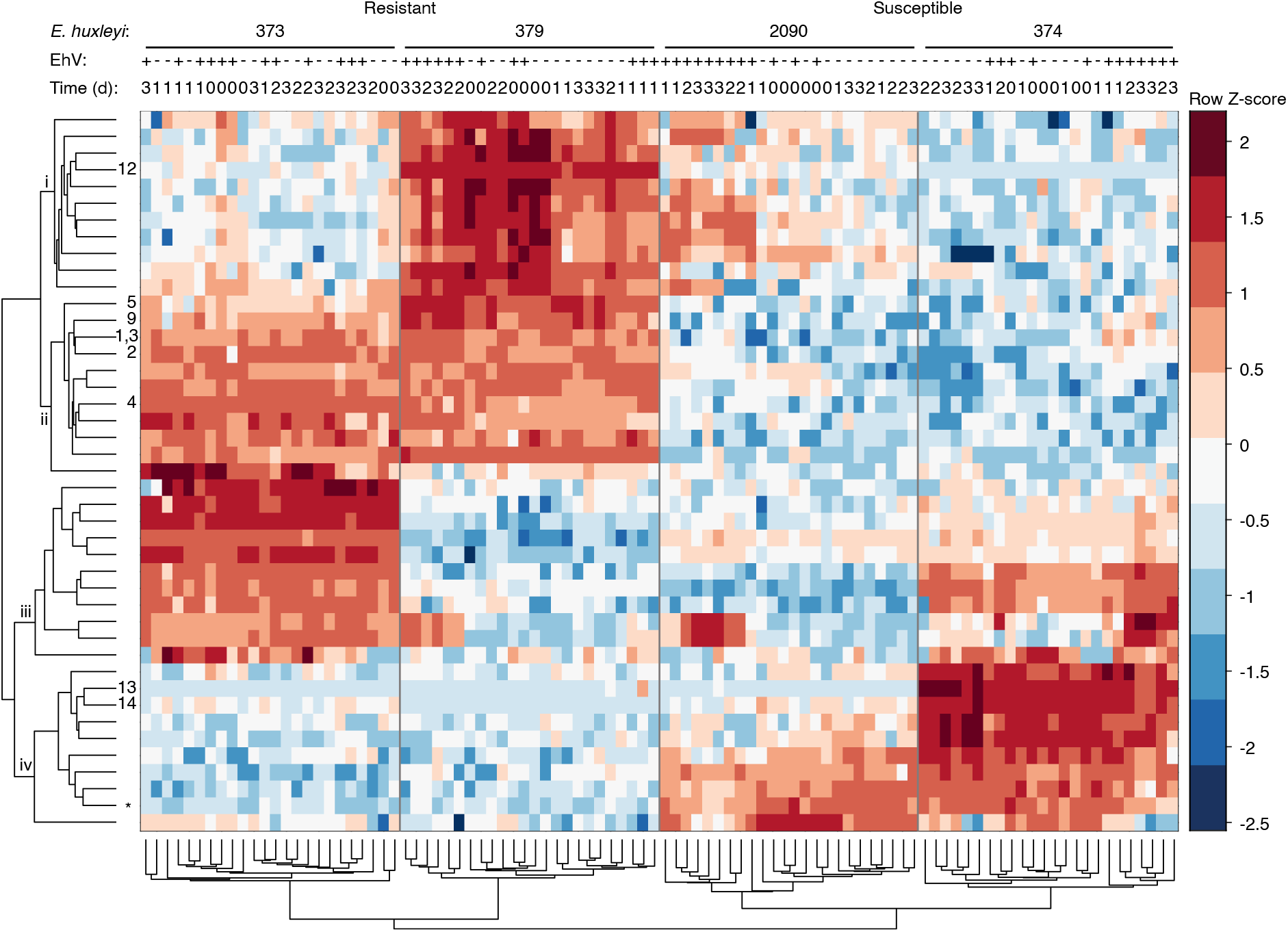
Two-dimensional hierarchical clustering of 43 differential lipid species (Table S1) in four *E. huxleyi* strains with and without EhV throughout a time course of four days. Clustering was performed on log-transformed and standardized peak areas of the adduct ion with the highest intensity. As in Fig. 2a, the samples are grouped into two main clusters that separate samples of the resistant (R) strains from the susceptible (S) ones (*n* = 96). Each cluster forms two sub-clusters that further separate the strains. The putative lipid species are also divided into four sub-clusters (i-iv), as in Fig. 2a. Nine GSL species were identified and are marked by numbers (Table 1). The peaks of GSLs **1** and **3**, structural isomers with a similar retention time (Table 1), were integrated together as they were not baseline separated. * sGSL d18:2/c22:0.

**Figure S10:**
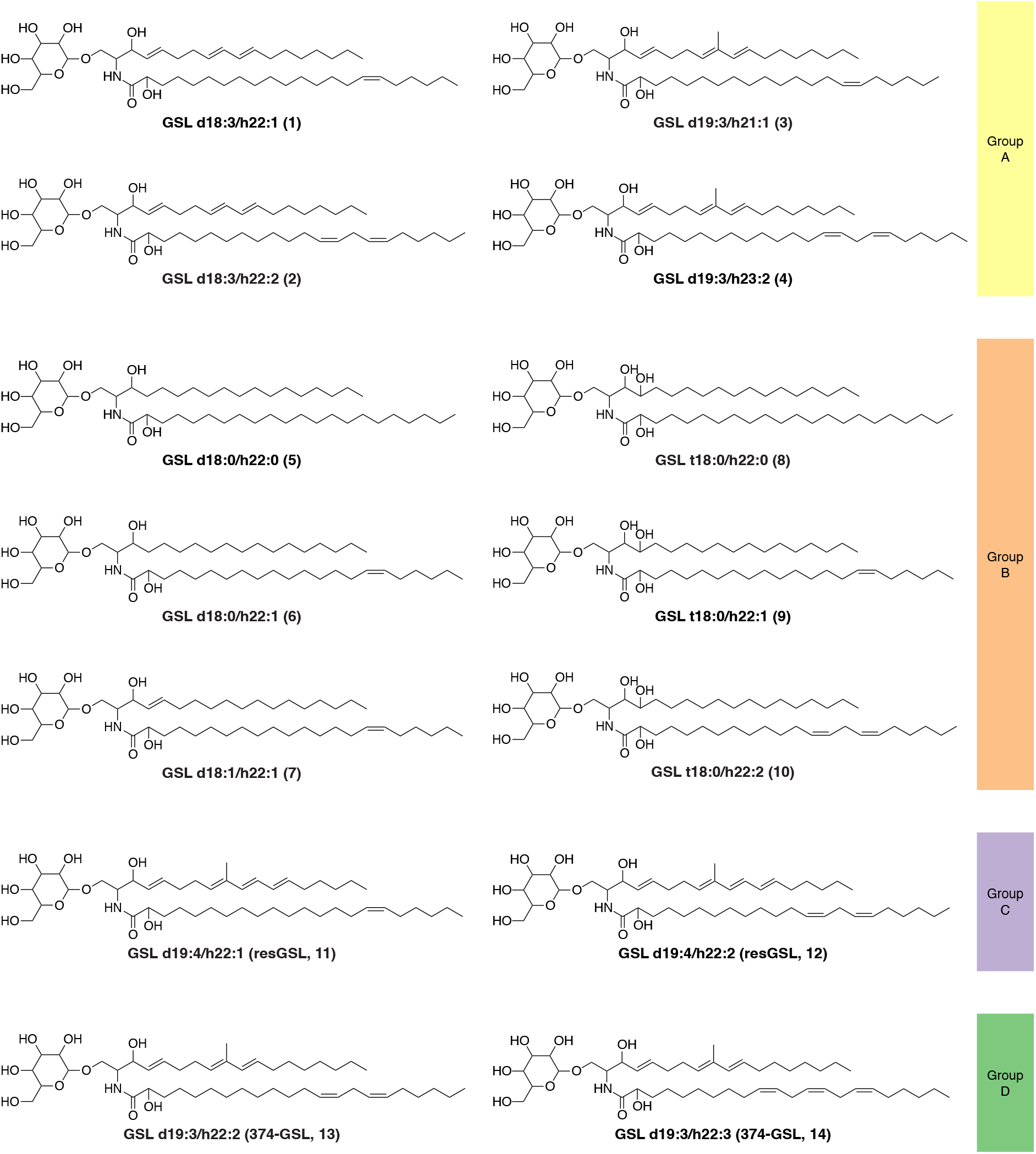
Structures of GSL species that differ between resistant and susceptible *E. huxleyi* strains and were identified in this study. Putative structures of previously undescribed GSL species within the *E. huxleyi*-EhV model system, which are differential between resistant and susceptible *E. huxleyi* strains (Table 1). The structures, including LCB and FA composition, were determined based on LC-MS/MS analysis (Fig. S4-S17). LCB composition, which varies in the amount of double bonds, hydroxyl groups and the alkyl chain branching, seems to be the main factor that differentiates between the various groups of GSLs, and, consequently, between the cell types. The positions of double bonds and functional groups were assigned based on the most common structures in the Lipid Maps Structure Database (LMSD) ^1^.

**Figure S11:**
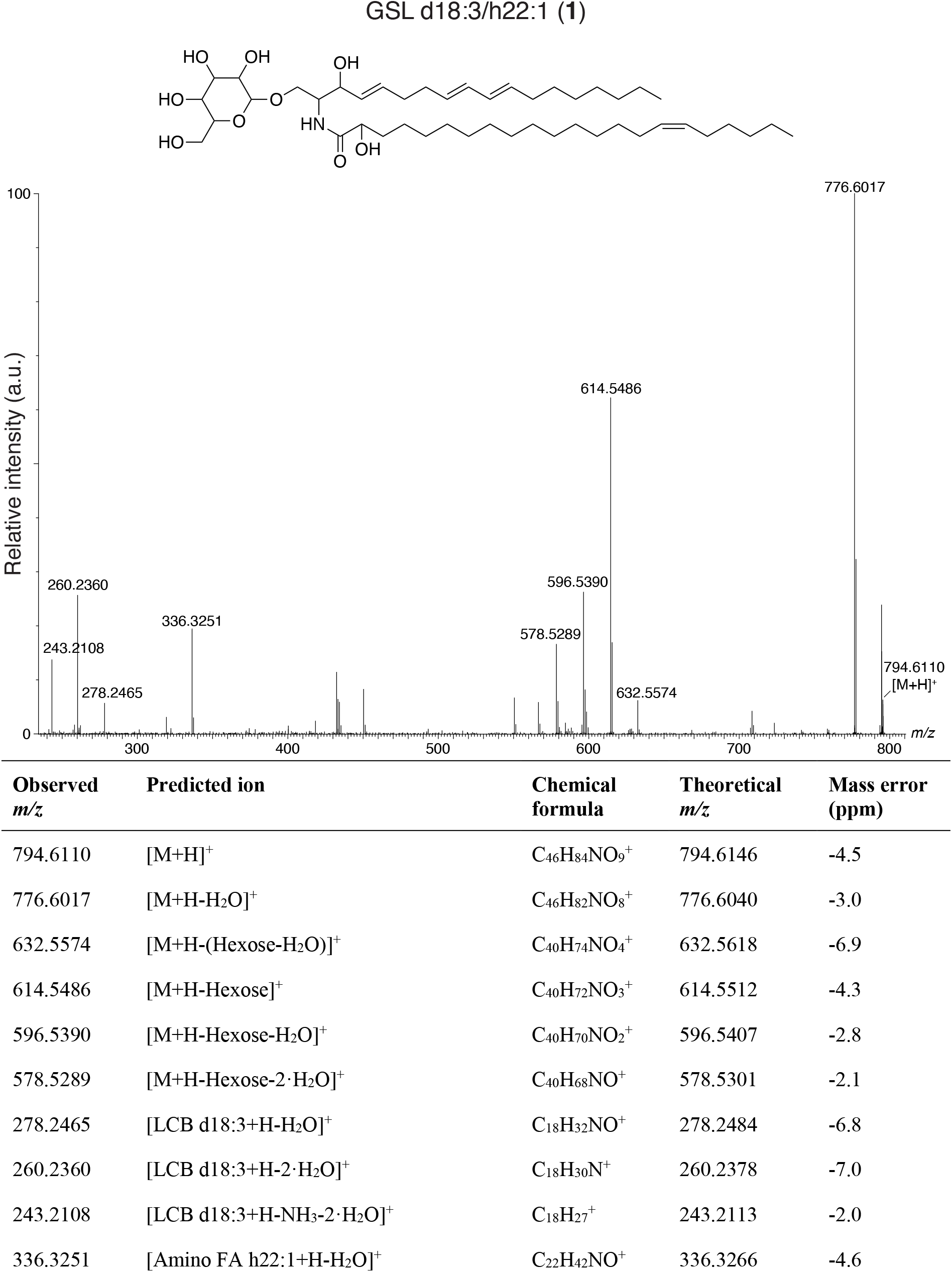
LC-MS/MS analysis of GSL d18:3/h22:1 (1). A putative structure is presented, supported by a list of fragments detected in MS/MS mode (Metabolomics Standards Initiative level 2 annotation^2^). Fragments were detected in positive ionization MS/MS mode using [M+H]^+^ = 794.6107 as the precursor ion (Table 1). The positions of the double bonds and functional groups were assigned based on the most common structures in the Lipid Maps Structure Database (LMSD) ^1^.

**Figure S12:**
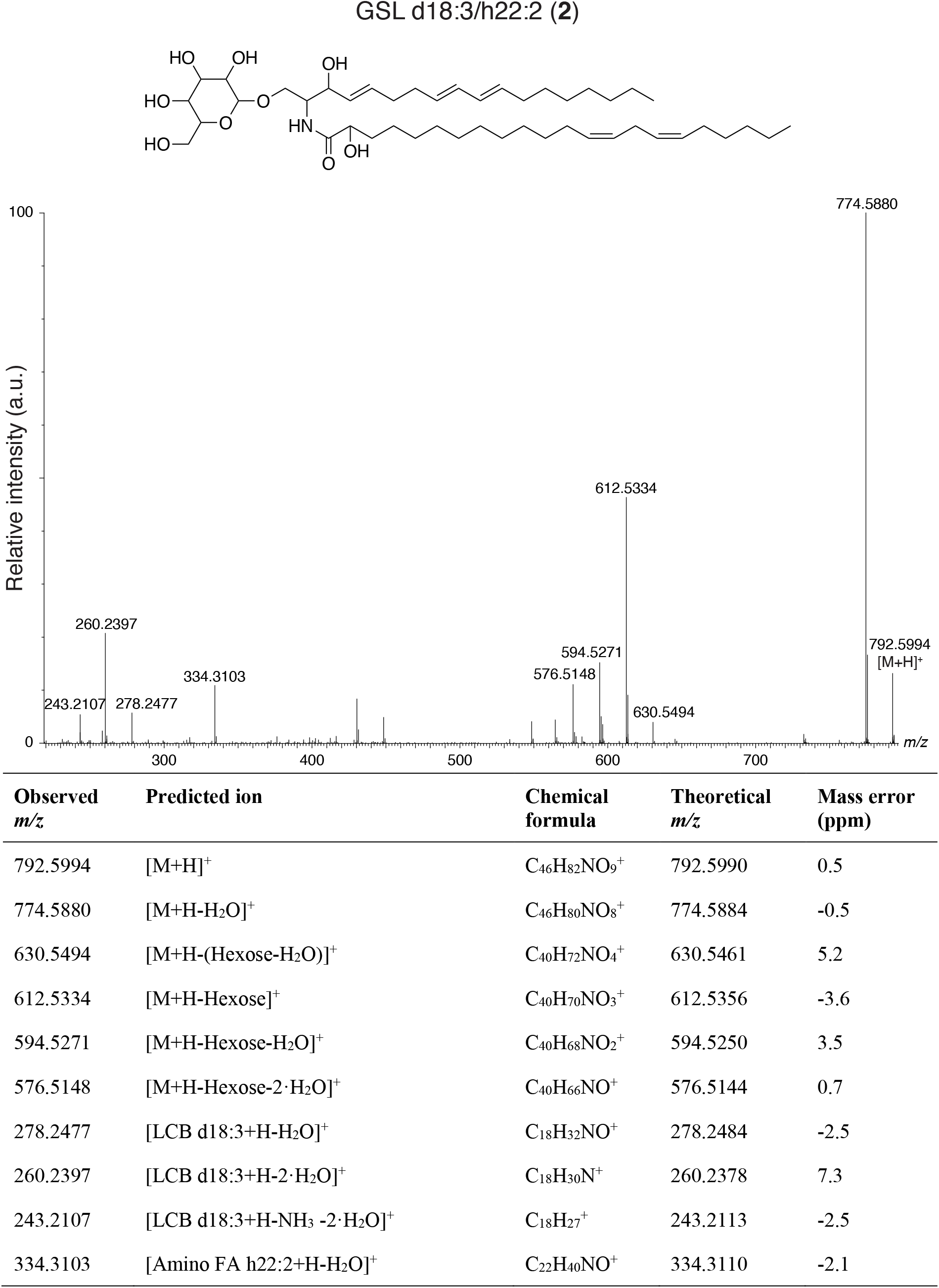
LC-MS/MS analysis of GSL d18:3/h22:2 (2). A putative structure is presented, supported by a list of fragments detected in MS/MS mode (Metabolomics Standards Initiative level 2^2^). Fragments were detected in positive ionization MS/MS mode using [M+H]^+^ = 792.5980 as the precursor ion (Table 1). The positions of the double bonds and functional groups were assigned based on the most common structures in the Lipid Maps Structure Database (LMSD) ^1^.

**Figure S13:**
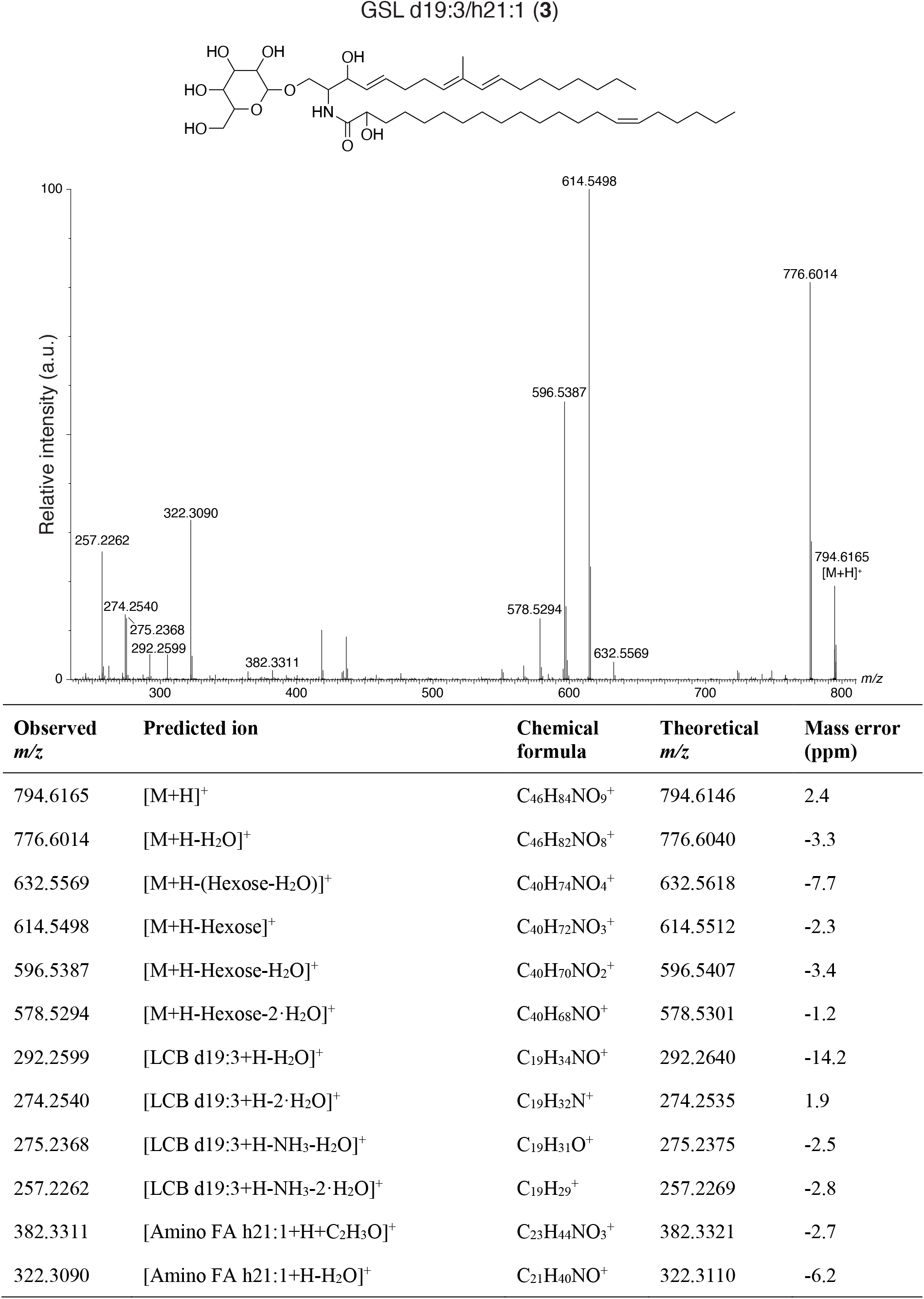
LC-MS/MS analysis of GSL d19:3/h21:1 (3). A putative structure is presented, supported by a list of fragments detected in MS/MS mode (Metabolomics Standards Initiative level 2 annotation^2^). Fragments were detected in positive ionization MS/MS mode using [M+H]^+^ = 794.6107 as the precursor ion (Table 1). The positions of the double bonds and functional groups were assigned based on the most common structures in the Lipid Maps Structure Database (LMSD) ^1^.

**Figure S14:**
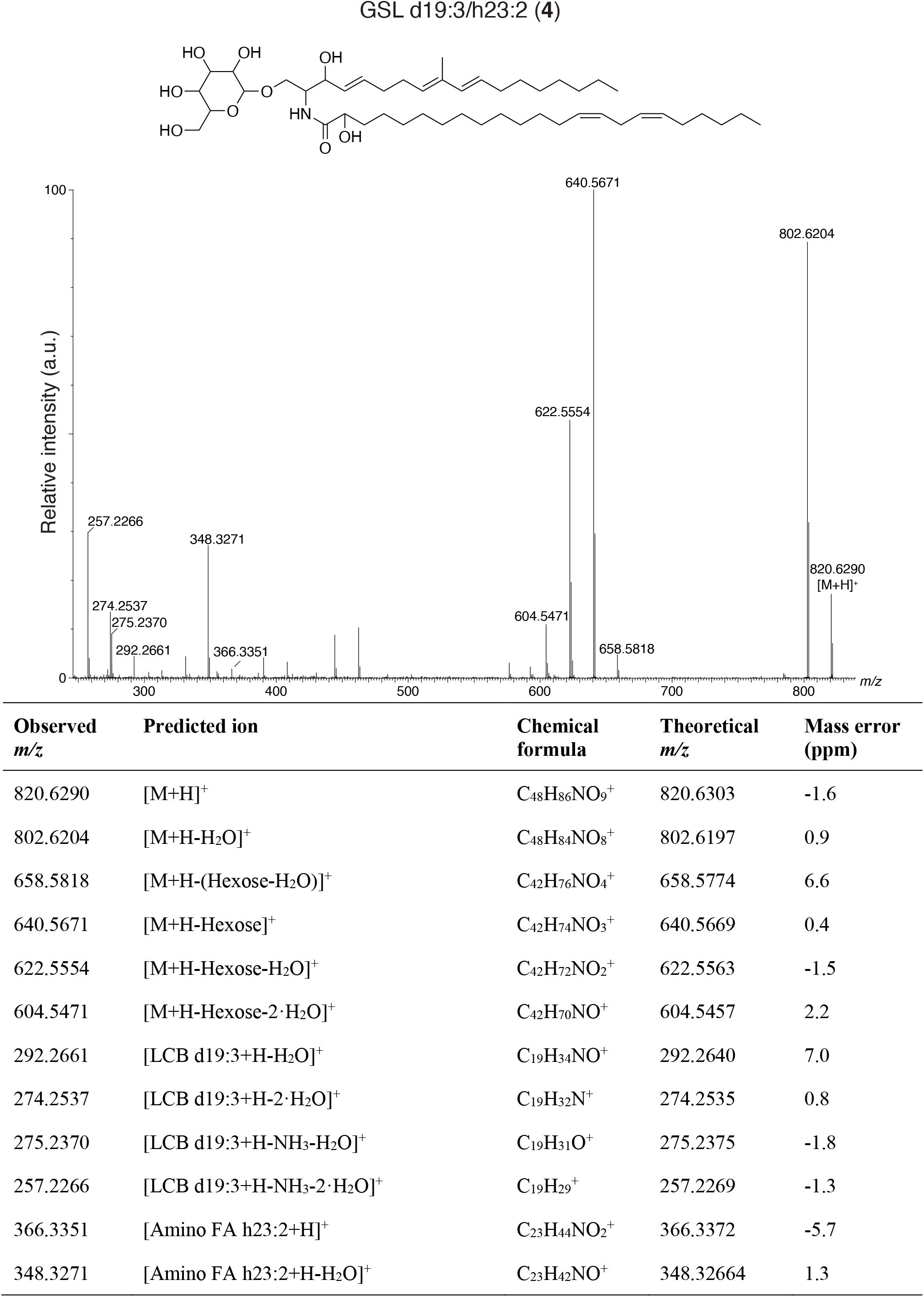
LC-MS/MS analysis of GSL d19:3/h23:2 (4). A putative structure is presented, supported by a list of fragments detected in MS/MS mode (Metabolomics Standards Initiative level 2 annotation^2^). Fragments were detected in positive ionization MS/MS mode using [M+H]^+^ = 820.6278 as the precursor ion (Table 1). The positions of the double bonds and functional groups were assigned based on the most common structures in the Lipid Maps Structure Database (LMSD) ^1^.

**Figure S15:**
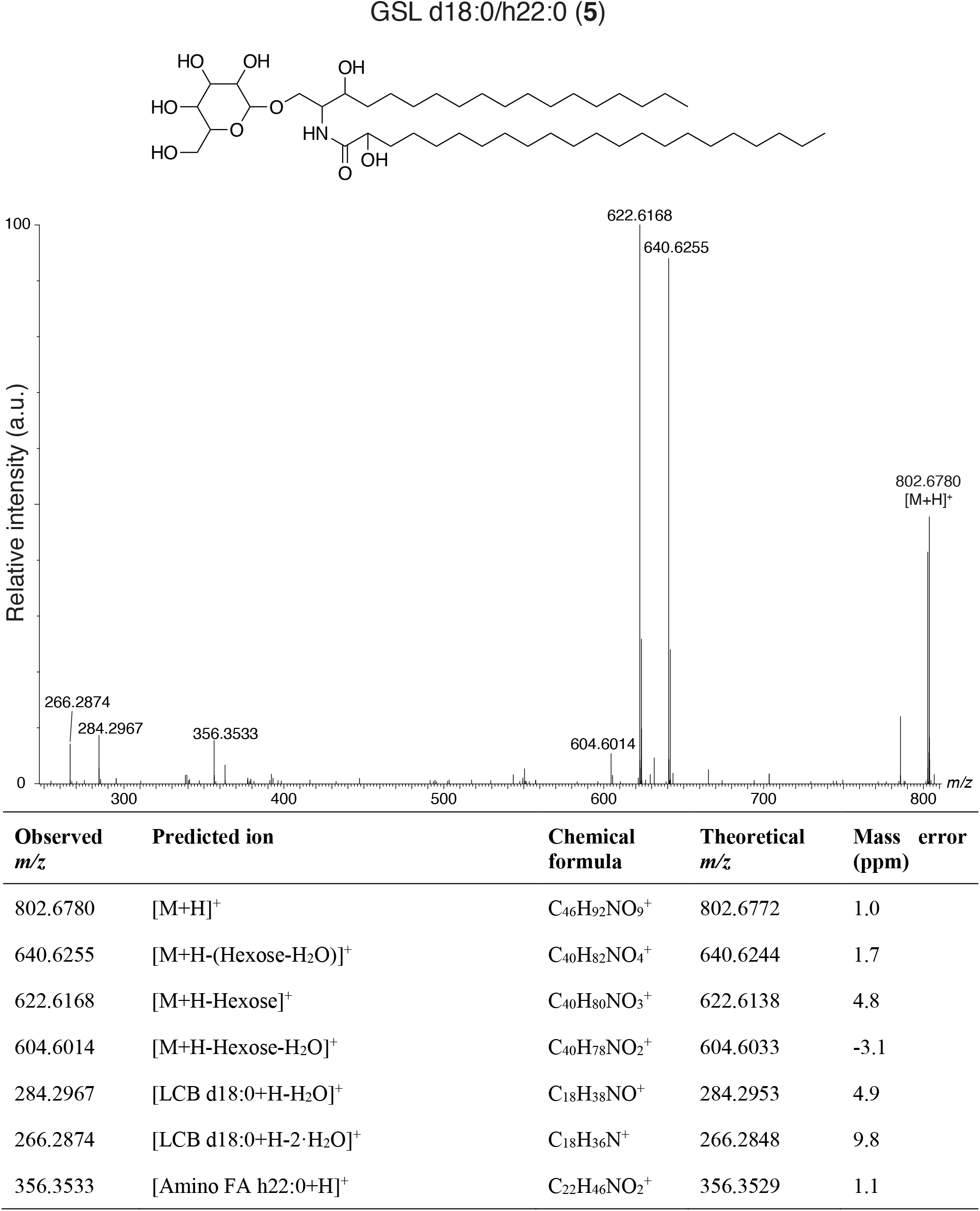
LC-MS/MS analysis of GSL d18:0/h22:0 (5). A putative structure is presented, supported by a list of fragments detected in MS/MS mode (Metabolomics Standards Initiative level 2 annotation^2^). Fragments were detected in positive ionization MS/MS mode using [M+H]^+^ = 802.6722 as the precursor ion (Table 1). The positions of the functional groups were assigned based on the most common structures in the Lipid Maps Structure Database (LMSD) ^1^.

**Figure S16:**
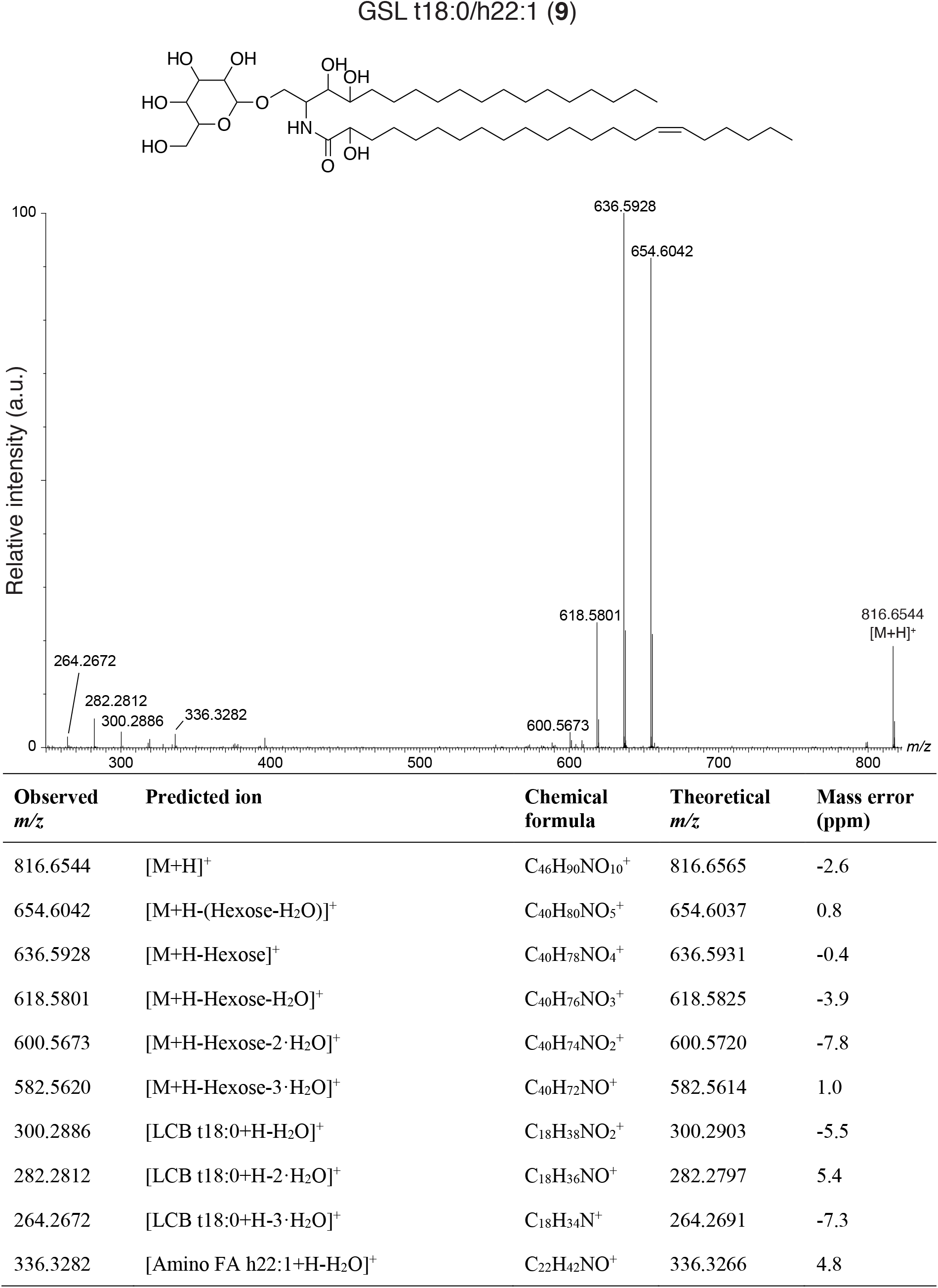
LC-MS/MS analysis of GSL t18:0/h22:1 (9). A putative structure is presented, supported by a list of fragments detected in MS/MS mode (Metabolomics Standards Initiative level 2 annotation^2^). Fragments were detected in positive ionization MS/MS mode using [M+H]^+^ = 816.6531 as the precursor ion (Table 1). The positions of the double bonds and functional groups were assigned based on the most common structures in the Lipid Maps Structure Database (LMSD) ^1^.

**Figure S17:**
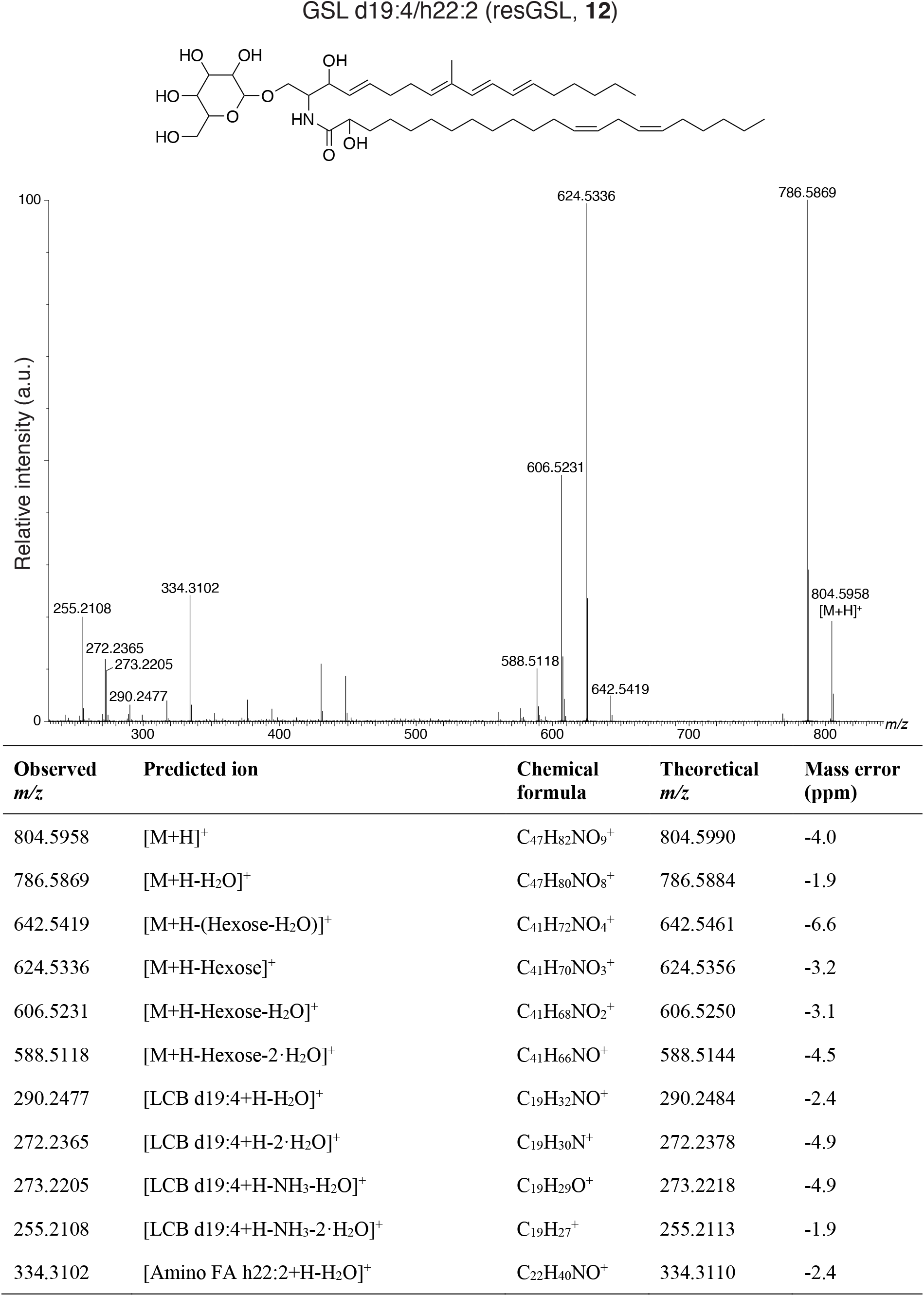
LC-MS/MS analysis of GSL d19:4/h22:2 (resGSL, 12). A putative structure is presented, supported by a list of fragments detected in MS/MS mode (Metabolomics Standards Initiative level 2 annotation^2^). Fragments were detected in positive ionization MS/MS mode using [M+H]^+^ = 804.5975 as the precursor ion (Table 1). The positions of the double bonds and functional groups were assigned based on the most common structures in the Lipid Maps Structure Database (LMSD) ^1^.

**Figure S18:**
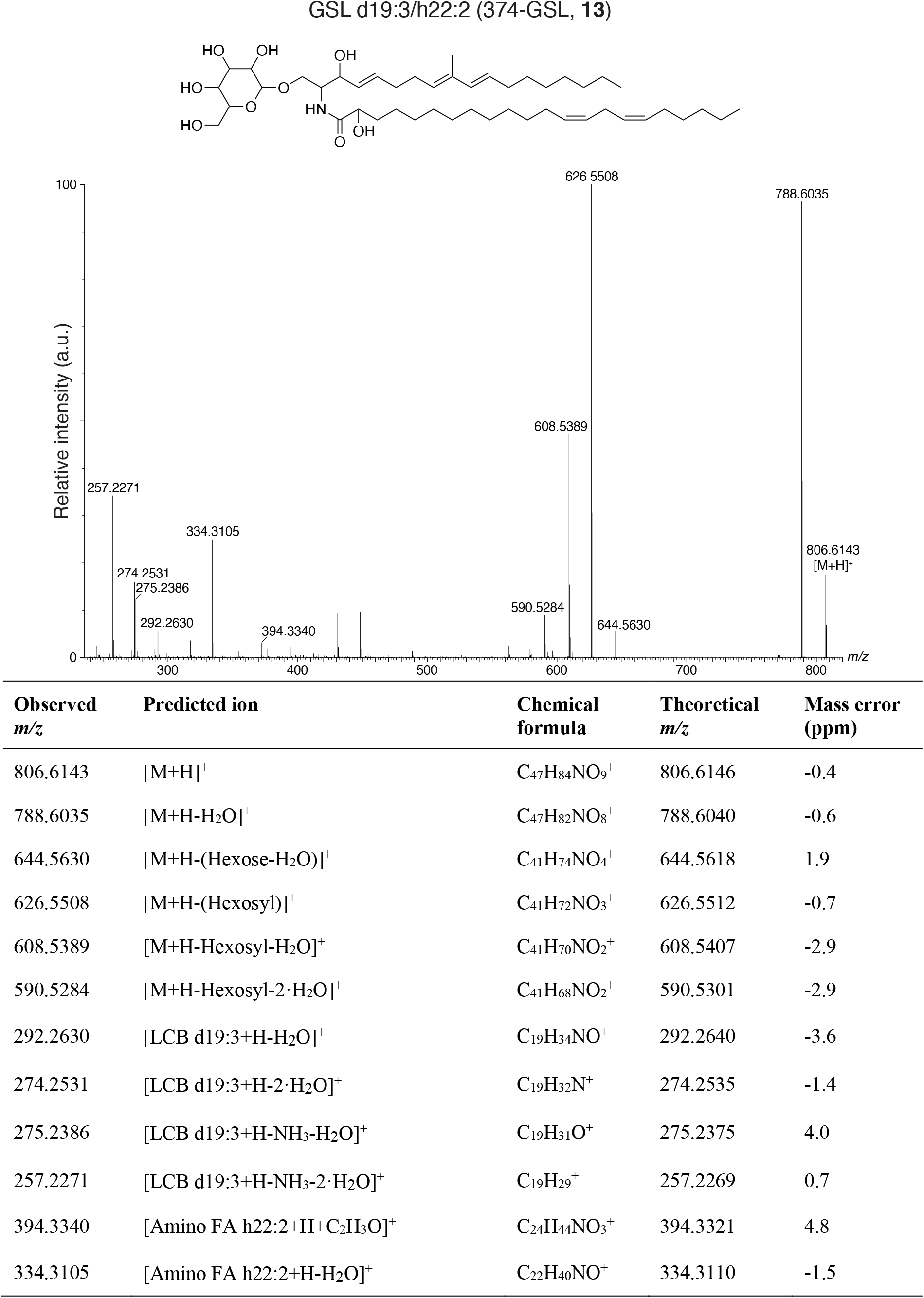
LC-MS/MS analysis of GSL d19:3/h22:2 (374-GSL, 13). A putative structure is presented, supported by a list of fragments detected in MS/MS mode (Metabolomics Standards Initiative level 2 annotation^2^). Fragments were detected in positive ionization MS/MS mode using [M+H]^+^ = 806.6143 as the precursor ion (Table 1). The positions of the double bonds and functional groups were assigned based on the most common structures in the Lipid Maps Structure Database (LMSD) ^1^.

**Figure S19:**
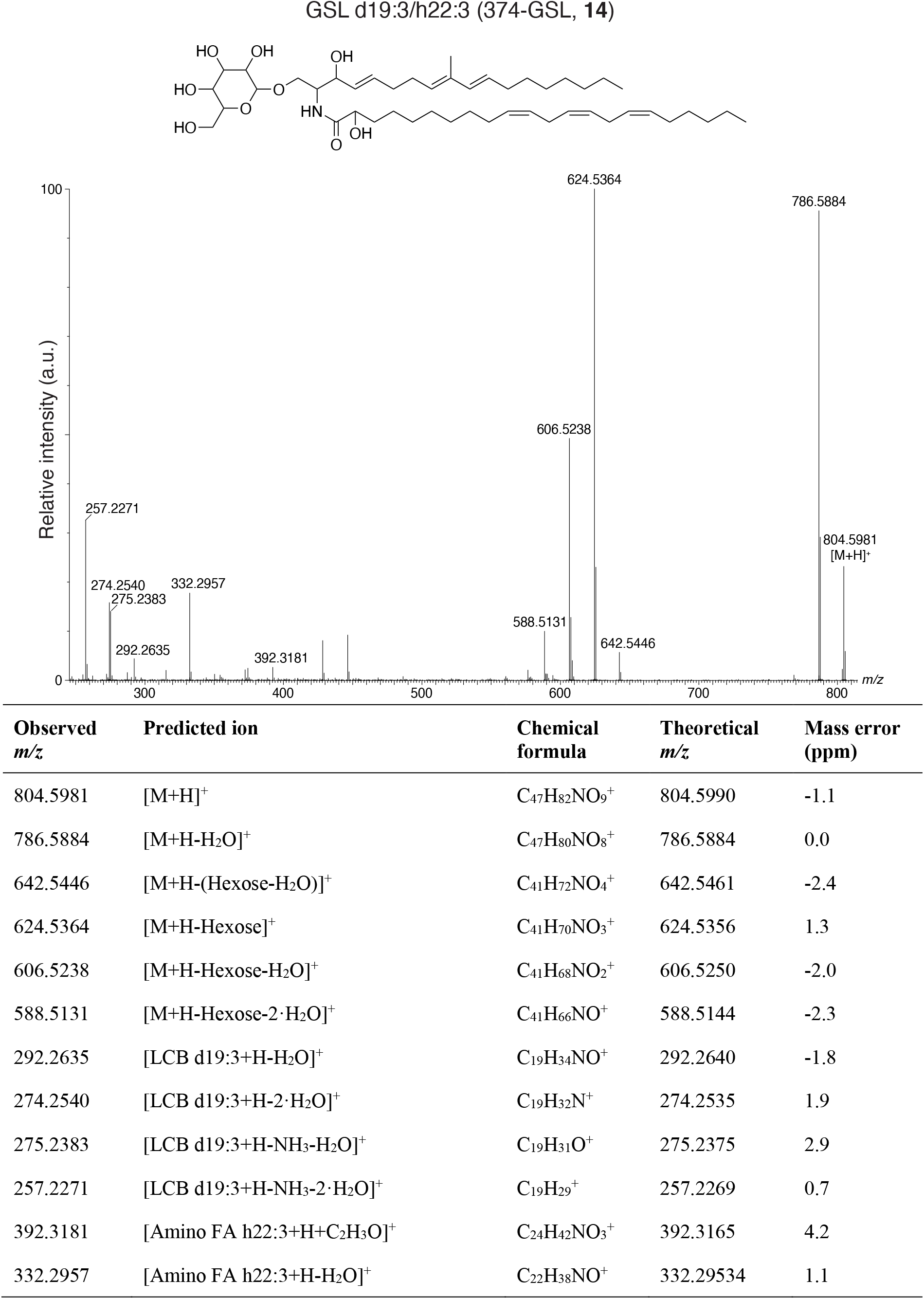
LC-MS/MS analysis of GSL d19:3/h22:3 (374-GSL, 14). A putative structure is presented, supported by a list of fragments detected in MS/MS mode (Metabolomics Standards Initiative level 2 annotation^2^). Fragments were detected in positive ionization MS/MS mode using [M+H]^+^ = 804.5981 as the precursor ion (Table 1). The positions of the double bonds and functional groups were assigned based on the most common structures in the Lipid Maps Structure Database (LMSD) ^1^.

**Figure S20:**
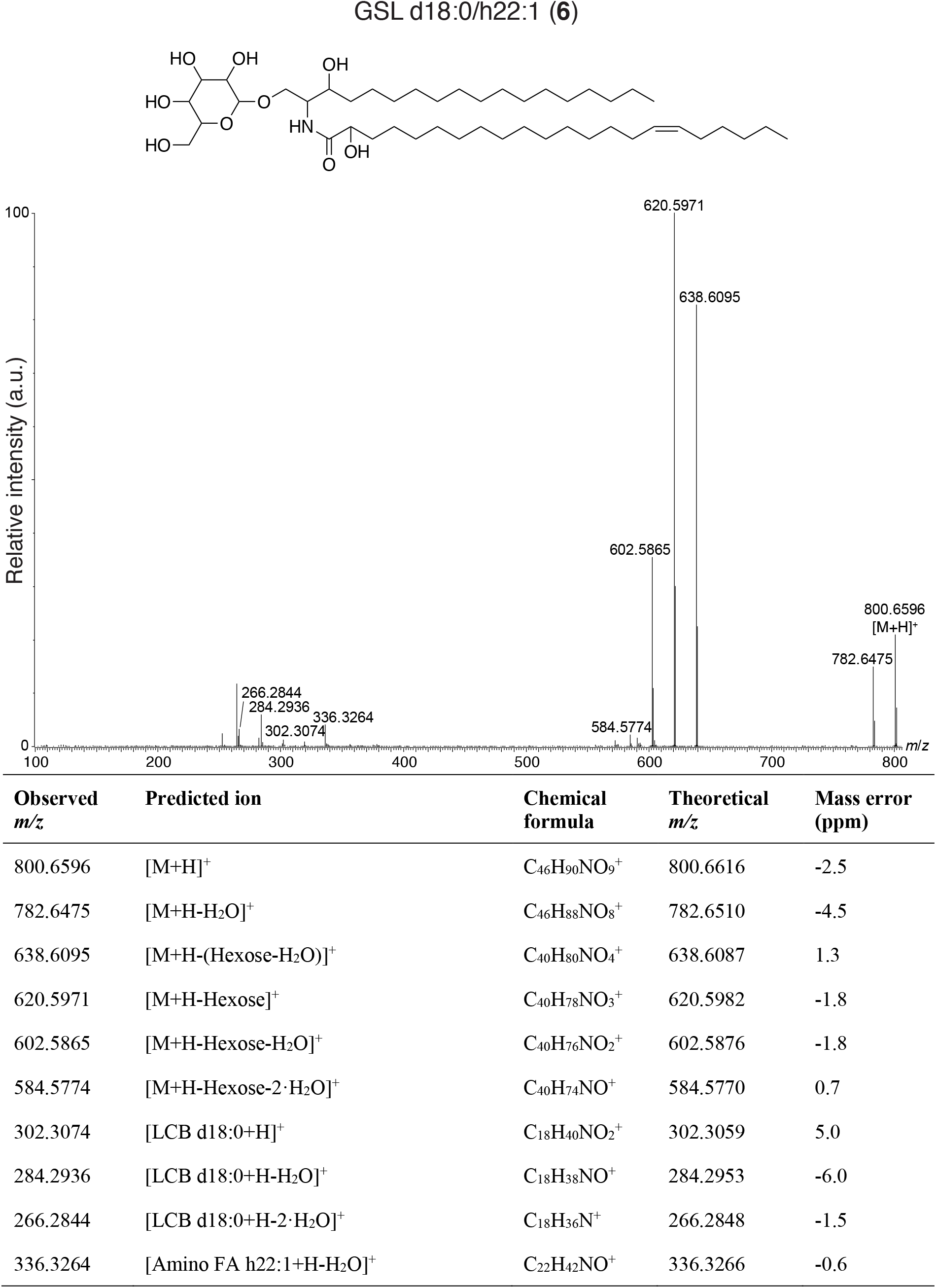
LC-MS/MS analysis of GSL d18:0/h22:1 (6). A putative structure is presented, supported by a list of fragments detected in MS/MS mode (Metabolomics Standards Initiative level 2 annotation^2^). Fragments were detected in positive ionization MS/MS mode using [M+H]^+^ = 800.6600 as the precursor ion (Table 1). The positions of the double bond and functional groups were assigned based on the most common structures in the

**Figure S21:**
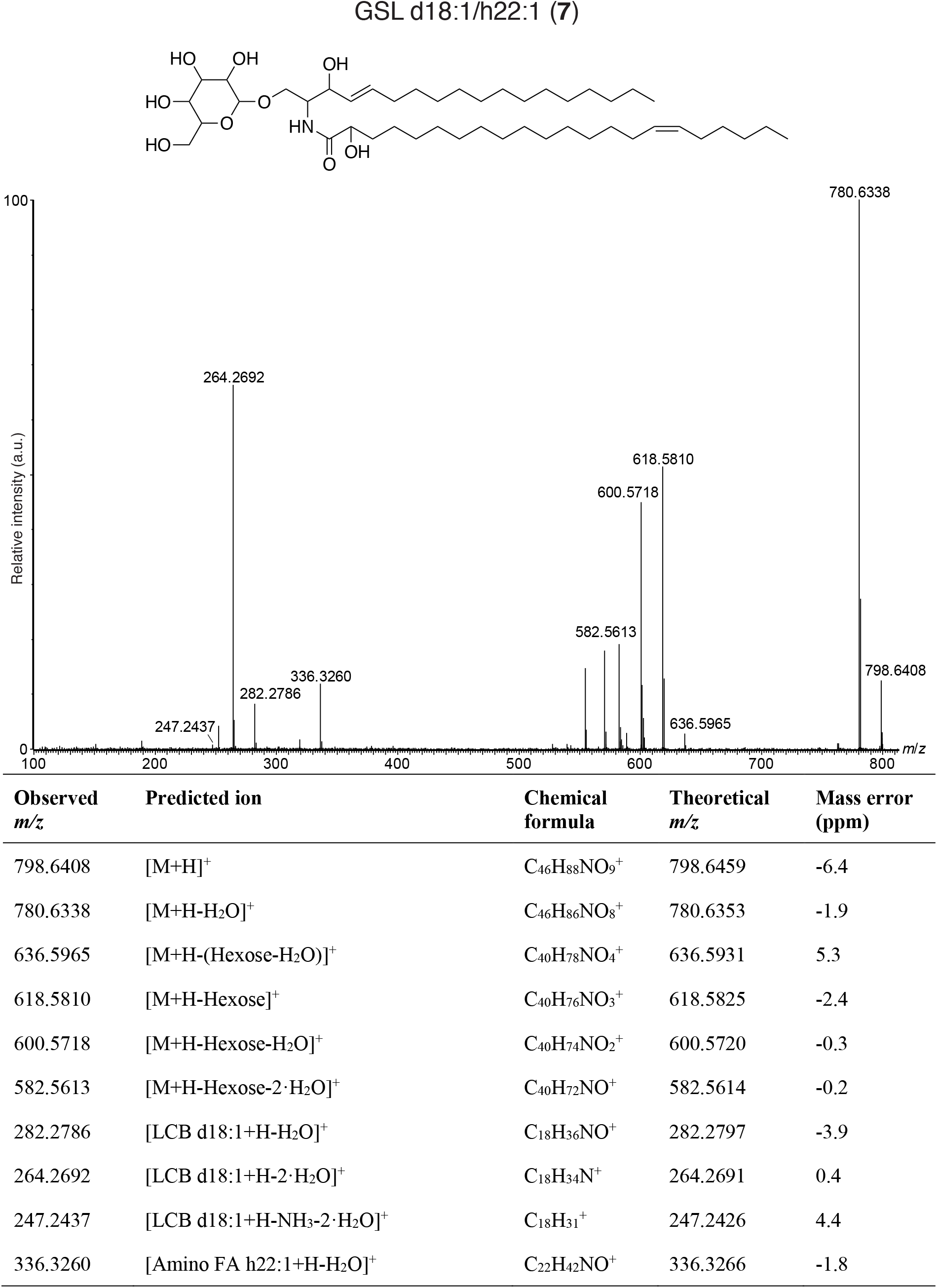
LC-MS/MS analysis of GSL d18:1/h22:1 (7). A putative structure is presented, supported by a list of fragments detected in MS/MS mode (Metabolomics Standards Initiative level 2 annotation^2^). Fragments were detected in positive ionization MS/MS mode using [M+H]^+^ = 798.6440 as the precursor ion (Table 1). The positions of the double bonds and functional groups were assigned based on the most common structures in the

**Figure S22:**
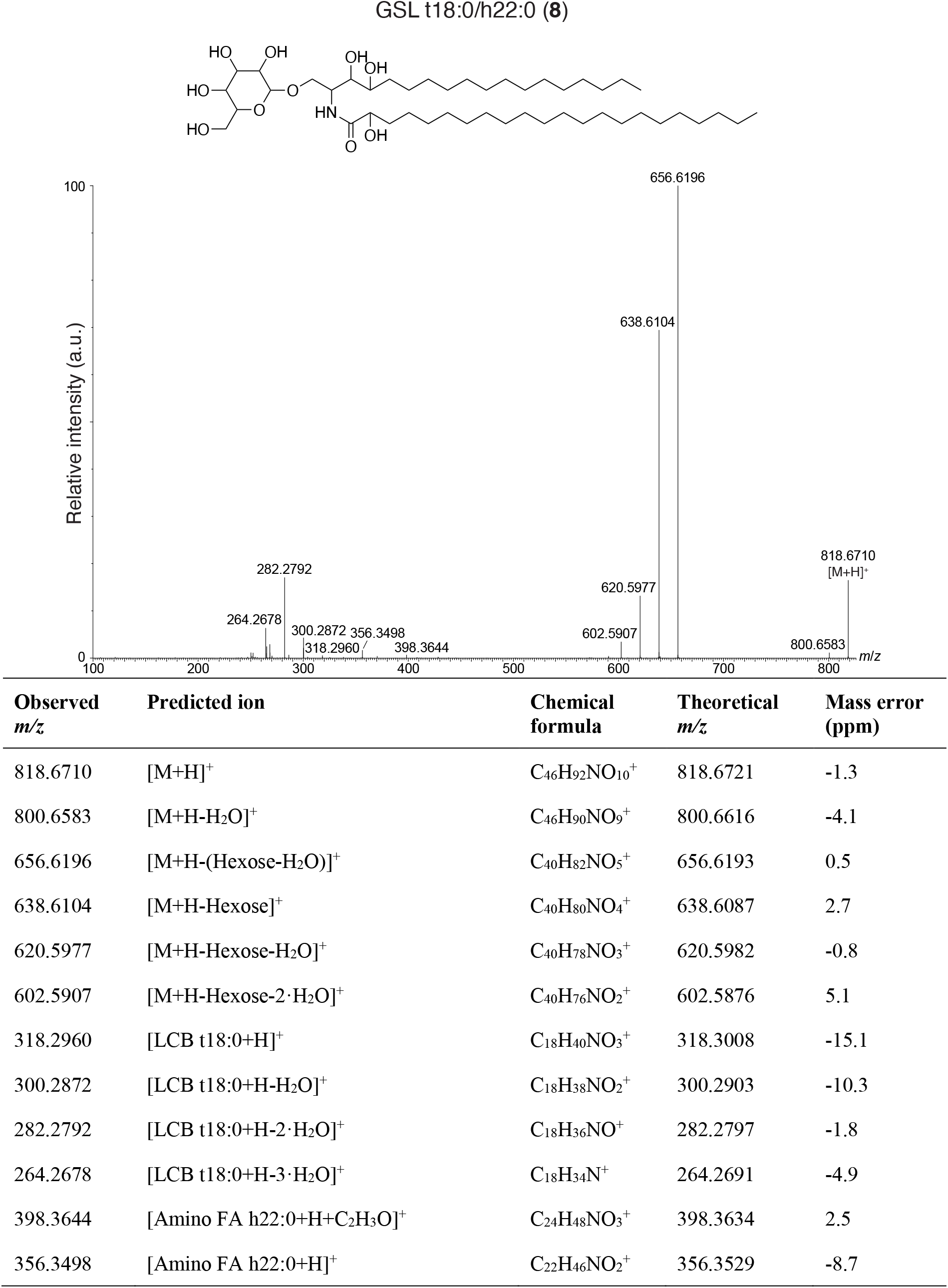
LC-MS/MS analysis of GSL t18:0/h22:0 (8). A putative structure is presented, supported by a list of fragments detected in MS/MS mode (Metabolomics Standards Initiative level 2 annotation^2^). Fragments were detected in positive ionization MS/MS mode using [M+H]^+^ = 818.6702 as the precursor ion (Table 1). The positions of the functional groups were assigned based on the most common structures in the Lipid Maps Structure Database (LMSD) ^1^.

**Figure S23:**
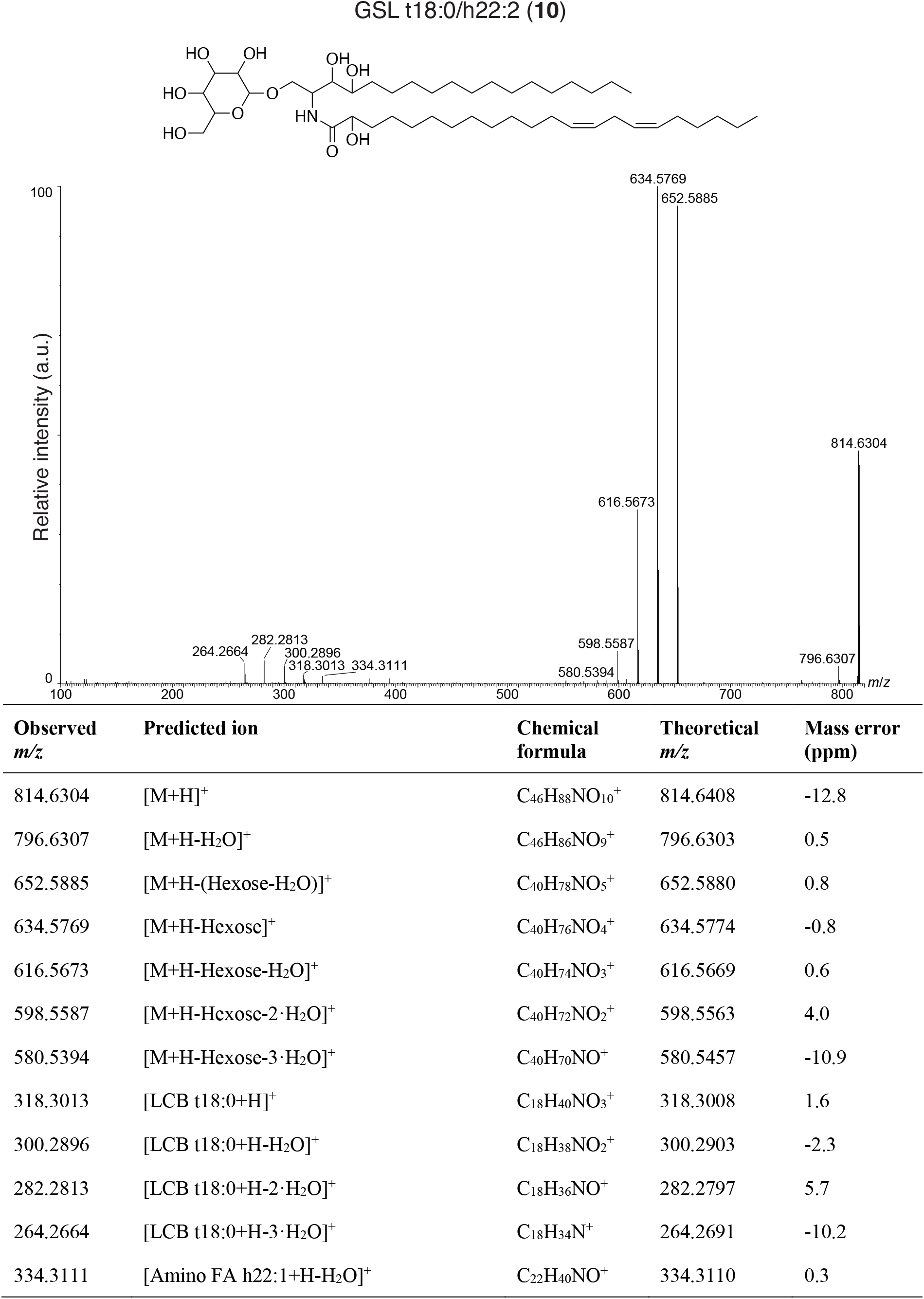
LC-MS/MS analysis of GSL t18:0/h22:2 (10). A putative structure is presented, supported by a list of fragments detected in MS/MS mode (Metabolomics Standards Initiative level 2 annotation^2^). Fragments were detected in positive ionization MS/MS mode using [M+H]^+^ = 814.6346 as the precursor ion (Table 1). The positions of the double bonds and functional groups were assigned based on the most common structures in the Lipid Maps Structure Database (LMSD) ^1^.

**Figure S24:**
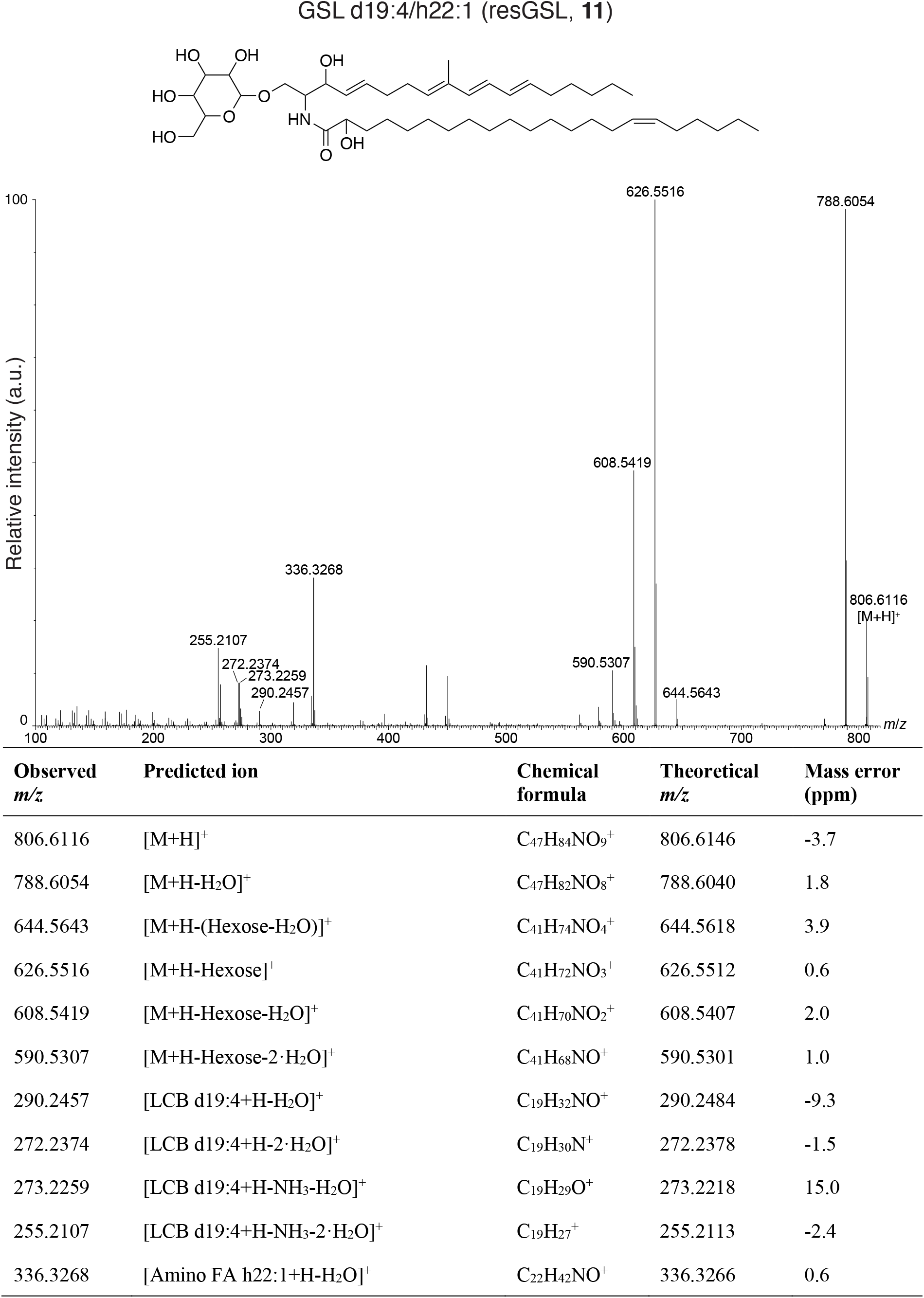
LC-MS/MS analysis of GSL d19:4/h22:1 (resGSL, 11). A putative structure is presented, supported by a list of fragments detected in MS/MS mode (Metabolomics Standards Initiative level 2 annotation^2^). Fragments were detected in positive ionization MS/MS mode using [M+H]^+^ = 806.6127 as the precursor ion (Table 1). The positions of the double bonds and functional groups were assigned based on the most common structures in the Lipid Maps Structure Database (LMSD) ^1^.

**Figure S25:**
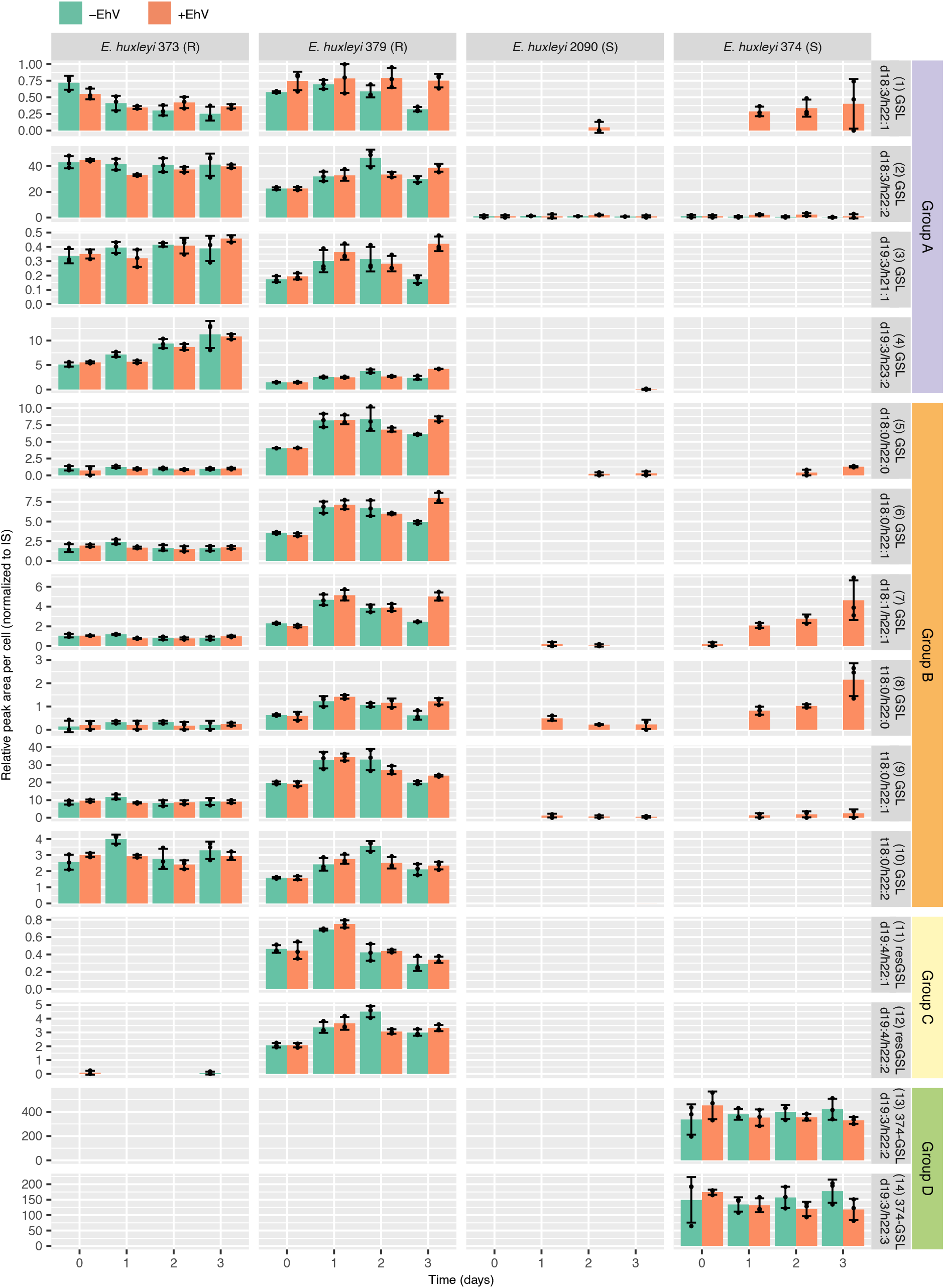
Temporal profiles of differential GSL species in two resistant (R) and two susceptible (S) *E. huxleyi* strains, with and without addition of EhV. The bar graphs show the mean relative peak area per cell ± SD (*n* = 3) of cultures without addition of EhV (–EhV) and with addition of EhV (+EhV). Peak areas were normalized to the internal standard (IS) glucosylceramide d18:1/c12:0. GSL species are numbered and ordered based on Table 1. An additional analysis with higher sensitivity was performed for some GSLs species, as shown in Fig. S19.

**Figure S26:**
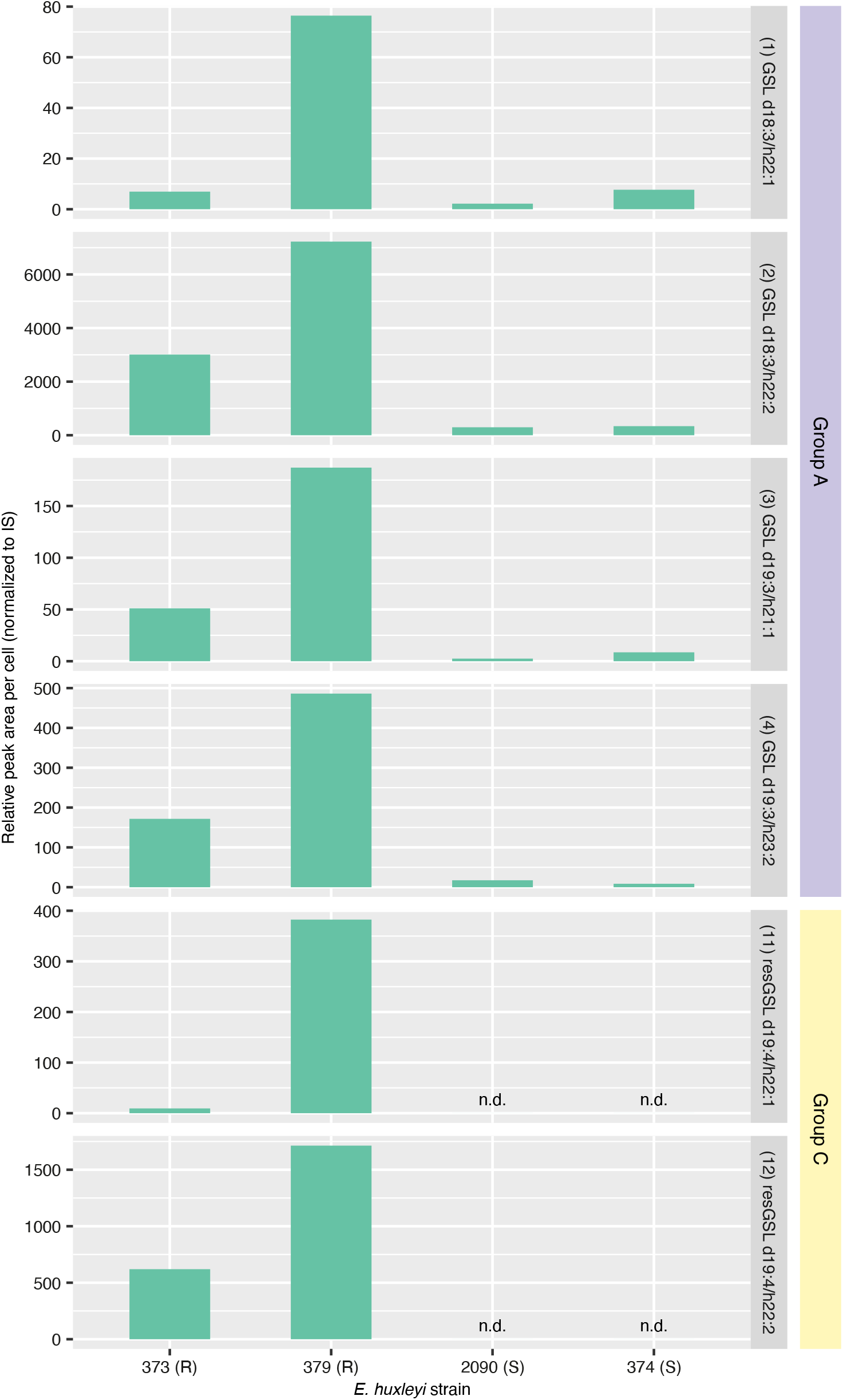
High sensitivity analysis of selected GSL species in two resistant (R) and two susceptible (S) *E. huxleyi* strains without addition of EhV. This analysis was performed to verify detection of GSL species that had low intensity in the untargeted lipidomics profiling. The bar graphs show the relative peak area per cell of each sample type (*n* = 1). Peak intensities were normalized to the IS glucosylceramide d18:1/c12:0. GSL species are numbered and ordered based on Table 1. n.d., not detected.

**Figure S27:**
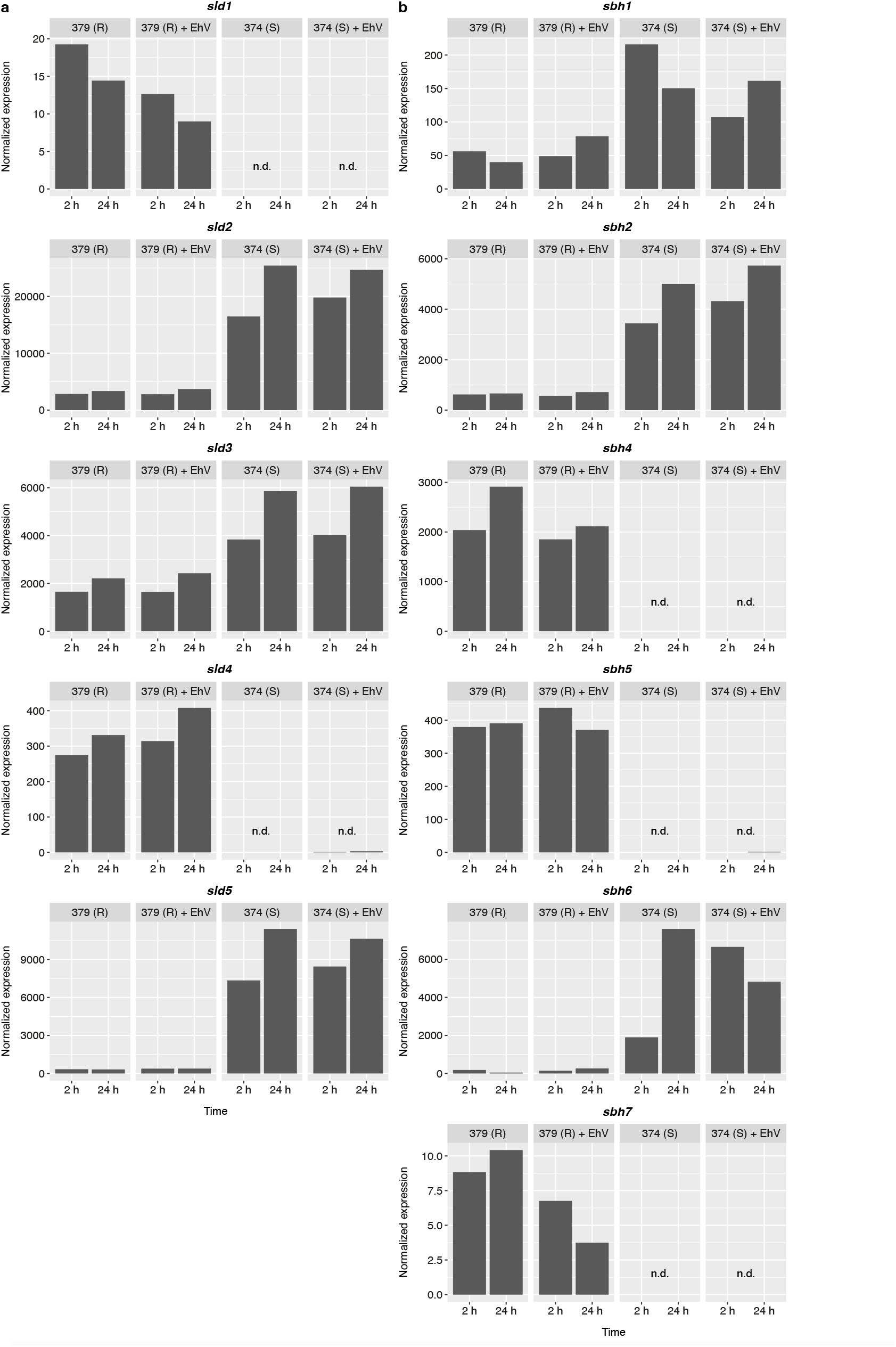
Expression of sphingolipid desaturase (*sld*) and sphingoid base hydroxylase (*sbh*) genes in the resistant *E. huxleyi* strain 379 and in the susceptible *E. huxleyi* strain 374 with and without addition of EhV. Normalized expression of (**a**) *sld* and (**b**) *sbh* genes in the resistant (R) *E. huxleyi* strain 379 and in the susceptible (S) *E. huxleyi* strain 374 with and without addition of EhV at 2 h and 24 h (*n* = 1). Expression of *sbh3* was below the limit of detection in all strains and conditions tested. Data was taken from the Marine Microbial Eukaryote Transcriptome Sequencing Project (MMETSP, *n* = 1, available at https://www.imicrobe.us, 3), samples MMETSP0994-MMETSP0997 (*E. huxleyi* 379) and MMETSP1006-MMETSP1009 (*E. huxleyi* 374). n.d., not detected.

**Figure S28:**
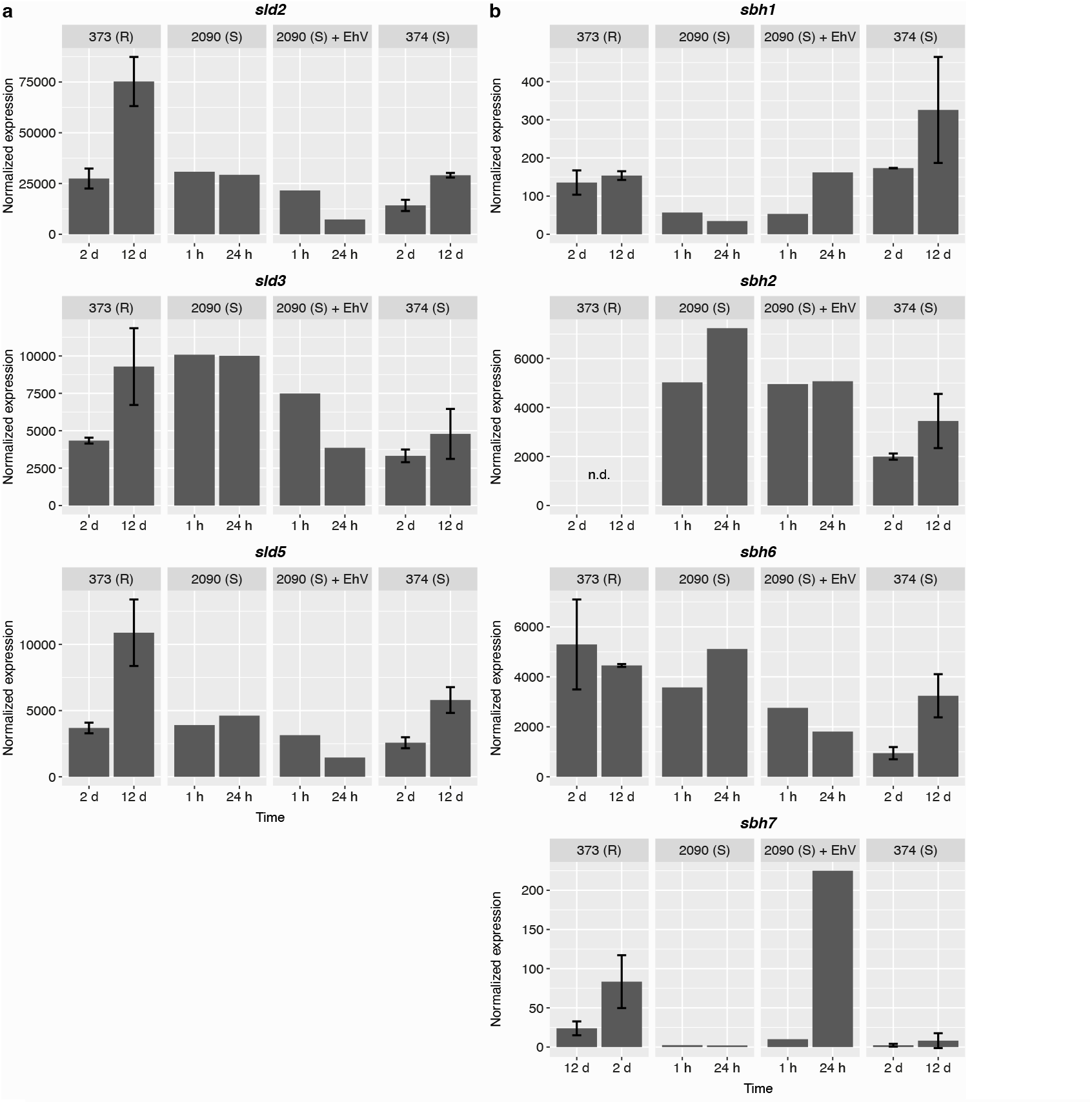
Expression of sphingolipid desaturase (*sld*) and sphingoid base hydroxylase (*sbh*) genes in *E. huxleyi* strain 373, 2090 and 374. Normalized expression of (**a**) *sld* and (**b**) *sbh* genes in the resistant (R) *E. huxleyi* strain 373 during the exponential (2 d) and stationary (12 d) growth phases, in the susceptible (S) *E. huxleyi* strain 2090 with and without addition of EhV at 1 h and 24 h, and in the susceptible *E. huxleyi* strain 374 during the exponential and stationary growth phases. Expression of *sbh3* was below limit of detection in all strains and conditions tested. Data was taken from Feldmesser *et al.* 2021^4^. Values for *E. huxleyi* strains 373 and 374 are presented as the mean ± SD (*n* = 2).

**Figure S29:**
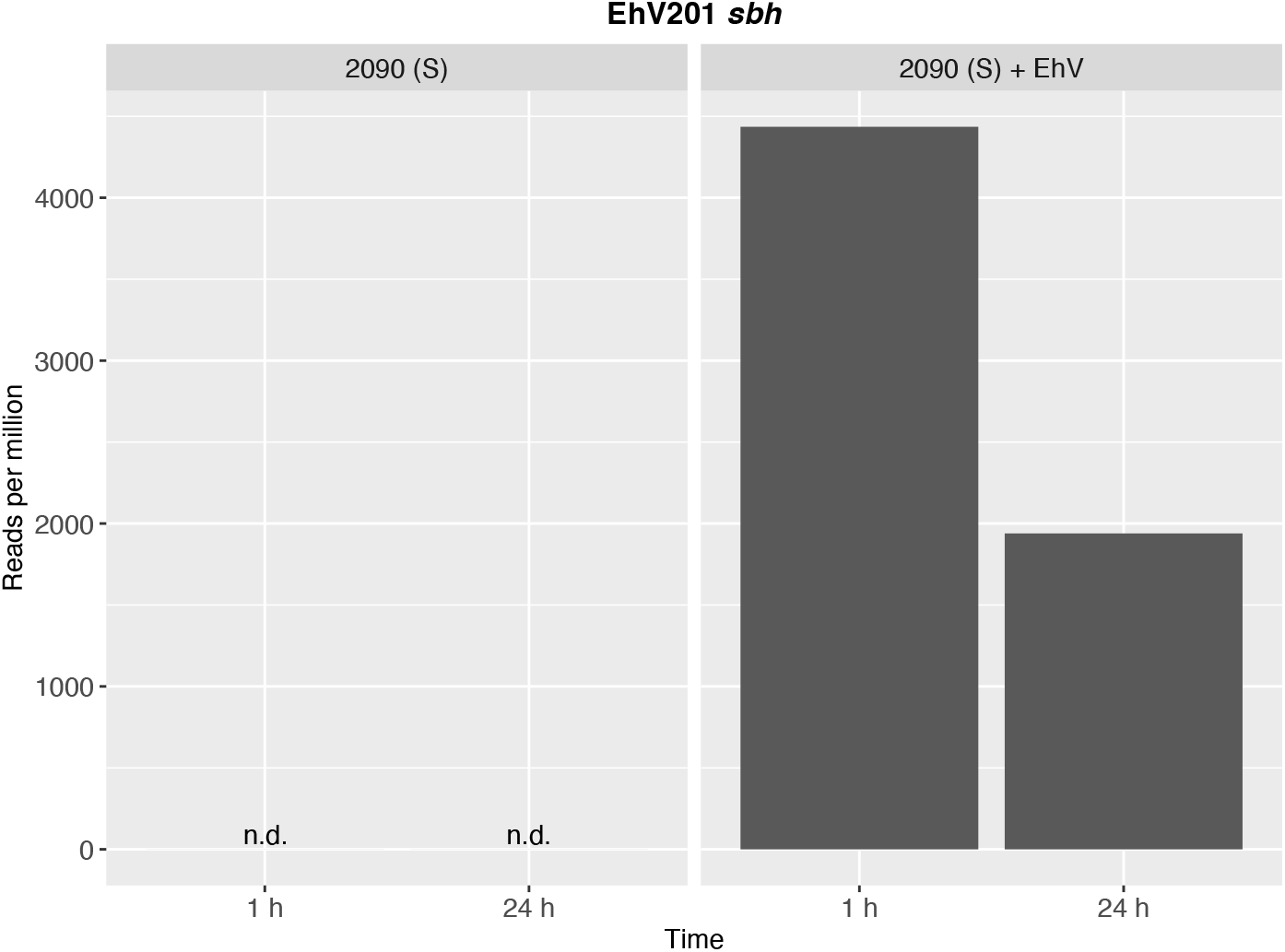
Expression of EhV201 sphingoid base hydroxylase (*sbh*) during infection of the susceptible *E. huxleyi* strain 2090. Data taken from Rosenwasser *et al.*, 2014^5^ (*n* = 1). n.d., not detected.

**Figure S30:**
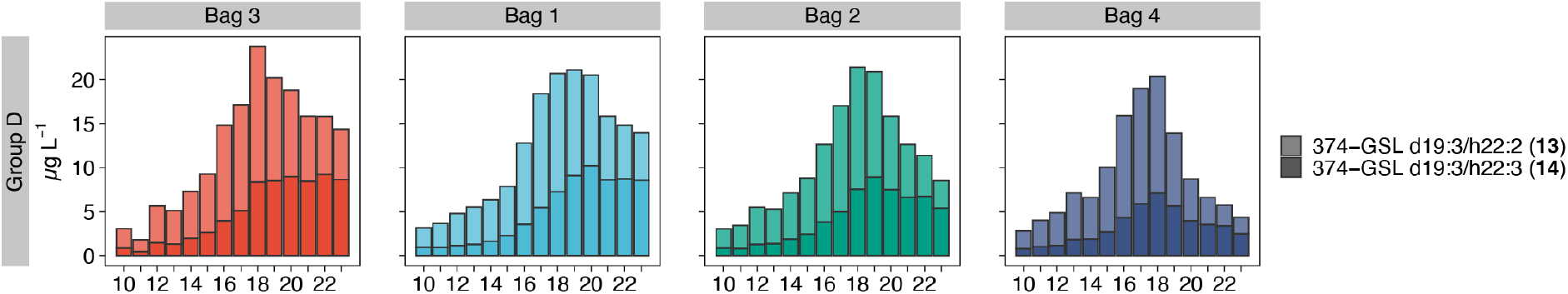
Changes in in cellular content of 374-GSL species (group D) in response to viral infection of natural *E. huxleyi* populations in a mesocosm experiment. Concentration of 374-GSL species correlates with the population dynamics in the four bags (Table S7). Bags are ordered by increasing EhV abundance, with the lowest abundance in bag 3 and the highest in bag 4, as presented in Fig. 5b.

**Figure S31:**
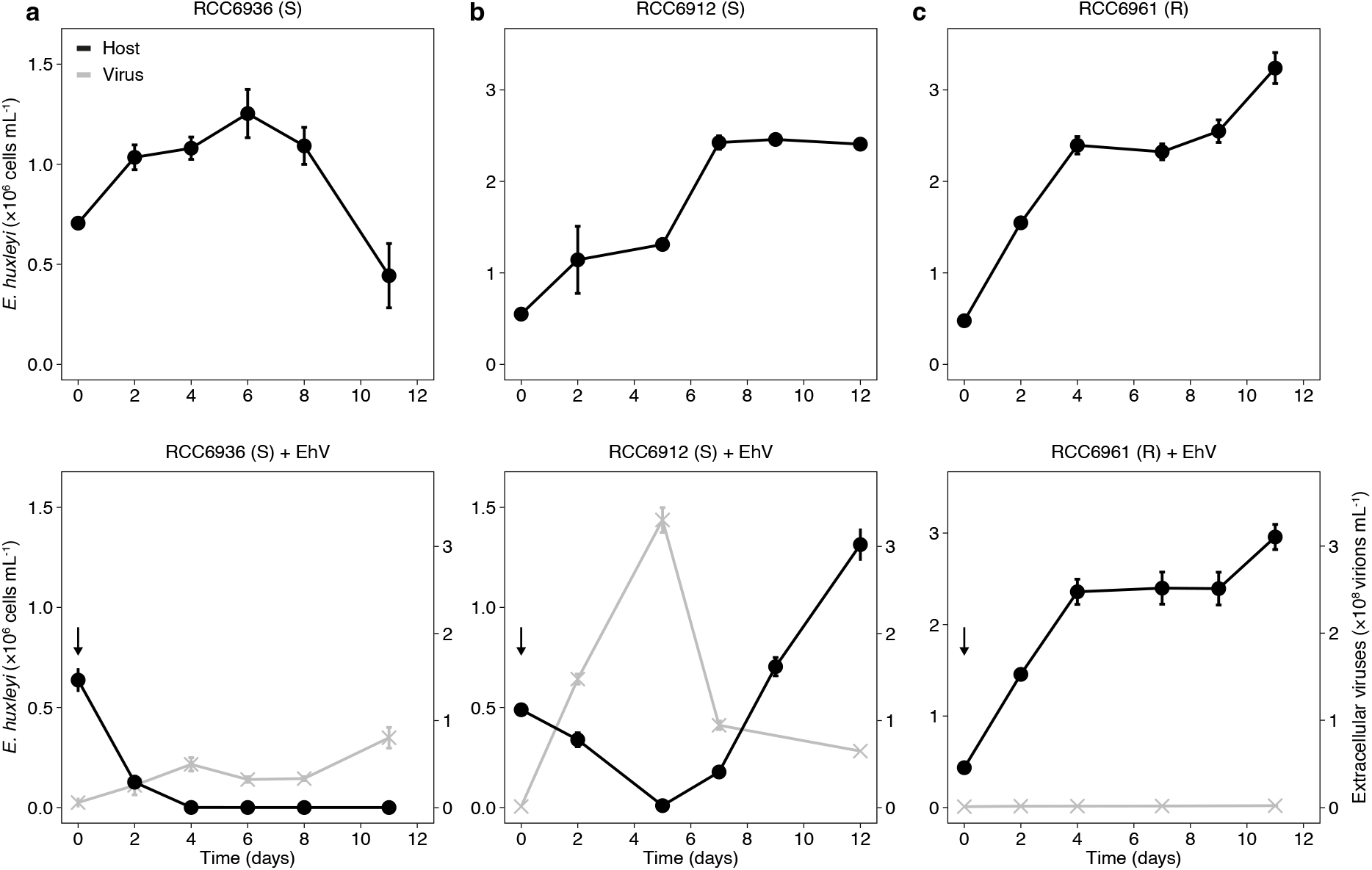
*E. huxleyi* isolates from the mesocosm experiment display different infection dynamics. (**a**) *E. huxleyi* isolate RCC6936 is susceptible (S) to EhVM1. (**b**) *E. huxleyi* isolate RCC6912 is susceptible to EhVM1, however, a resistant population recovers one week following infection. (**c**) *E. huxleyi* isolate RCC6961 is resistant (R) to EhVM1. *E. huxleyi* cell abundance (black) in cultures without (top) and with addition of EhVM1 (bottom) are presented, as well as the abundance of extracellular viruses (grey) in cultures with EhV. The black arrows indicate the addition of EhVM1 to the cultures. Values are presented as the mean ± SD (*n* = 2).

**Figure S32:**
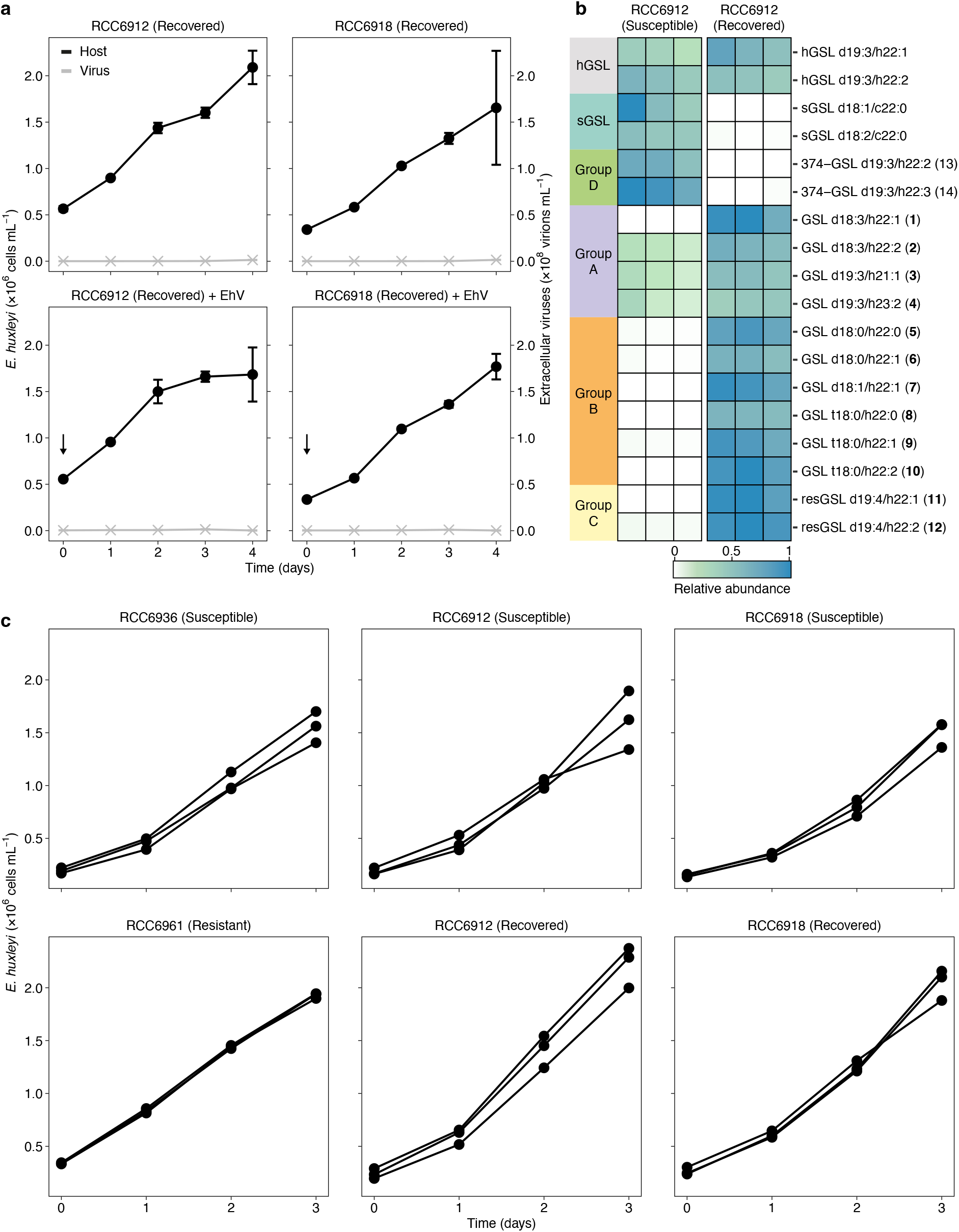
Resistance of the recovered cultures of *E. huxleyi* isolates RCC6912 and RCC6918 and the GSL composition of *E. huxleyi* isolate RCC6912. (**a**) Recovered cultures of *E. huxleyi* isolates RCC6912 and RCC6918 are resistant to the virus. *E. huxleyi* cell abundance (black) in cultures without (top) and with (bottom) addition of EhVM1 are presented, as well as abundance of extracellular viruses (grey) in cultures with EhVM1. Values are presented as the mean ± SD (*n* = 2). The black arrows indicate the addition of EhVM1 to the cultures. (**b**) GSL composition of the susceptible *E. huxleyi* isolate RCC6912 and of the cultures that recovered following infection and were resistant to the virus (*n* = 3). GSL species are grouped and numbered based on Table 1. (**c**) *E. huxleyi* cell abundance in exponentially growing cultures used for analysis of GSL composition (*n* = 3). The cultures were extracted at day 3 of the experiment.

**Figure S33:**
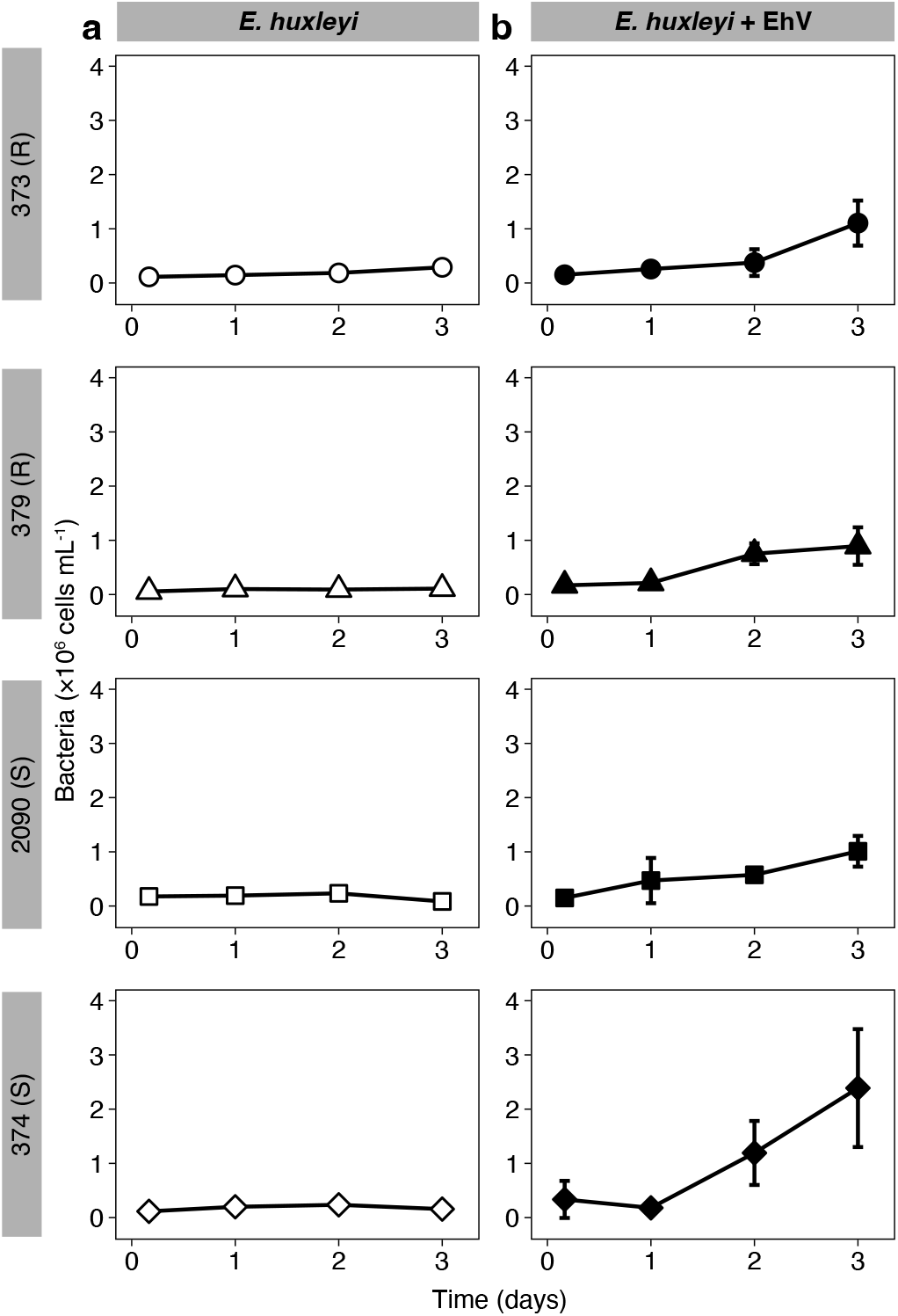
Quantification of bacteria in cultures of two resistant and two susceptible *E. huxleyi* strains, with and without addition of EhV. (**a**) Bacterial abundance during the growth of the resistant (R) *E. huxleyi* strains 373 and 379 and the susceptible (S) *E. huxleyi* strains 2090 and 374. (**b**) Bacterial abundance following the addition of EhV201. Values for (a) and (b) are presented as the mean ± SD (*n* = 3).

**Figure S34:**
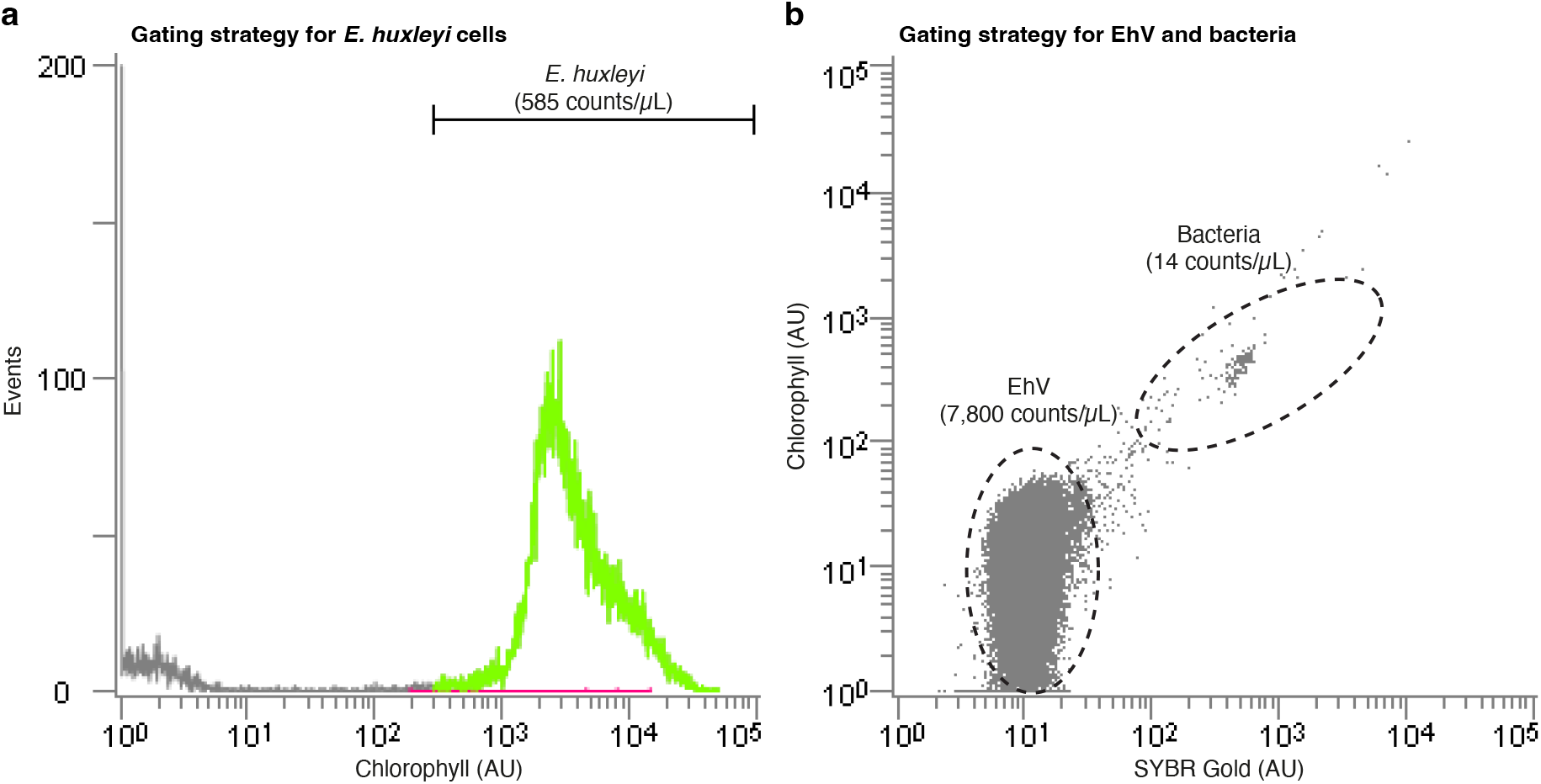
Enumeration of *E. huxleyi* cells, EhV and bacteria using flow cytometry. (a) Enumeration of *E. huxleyi* cells. Cells were identified by plotting the autofluorescence of chlorophyll (ex: 488, em: 663-737 nm). A threshold was applied based on the forward scatter signal to reduce the background noise. (**b**) Enumeration of EhV and bacteria. Flow cytometric analysis was performed with excitation at 488 nm and emission at 525 nm. A threshold was applied based on the forward scatter signal to reduce the background noise. The gates ‘EhV’ and ‘Bacteria’ were set by comparing to reference samples containing either EhV201 or bacteria.

**Figure S35:**
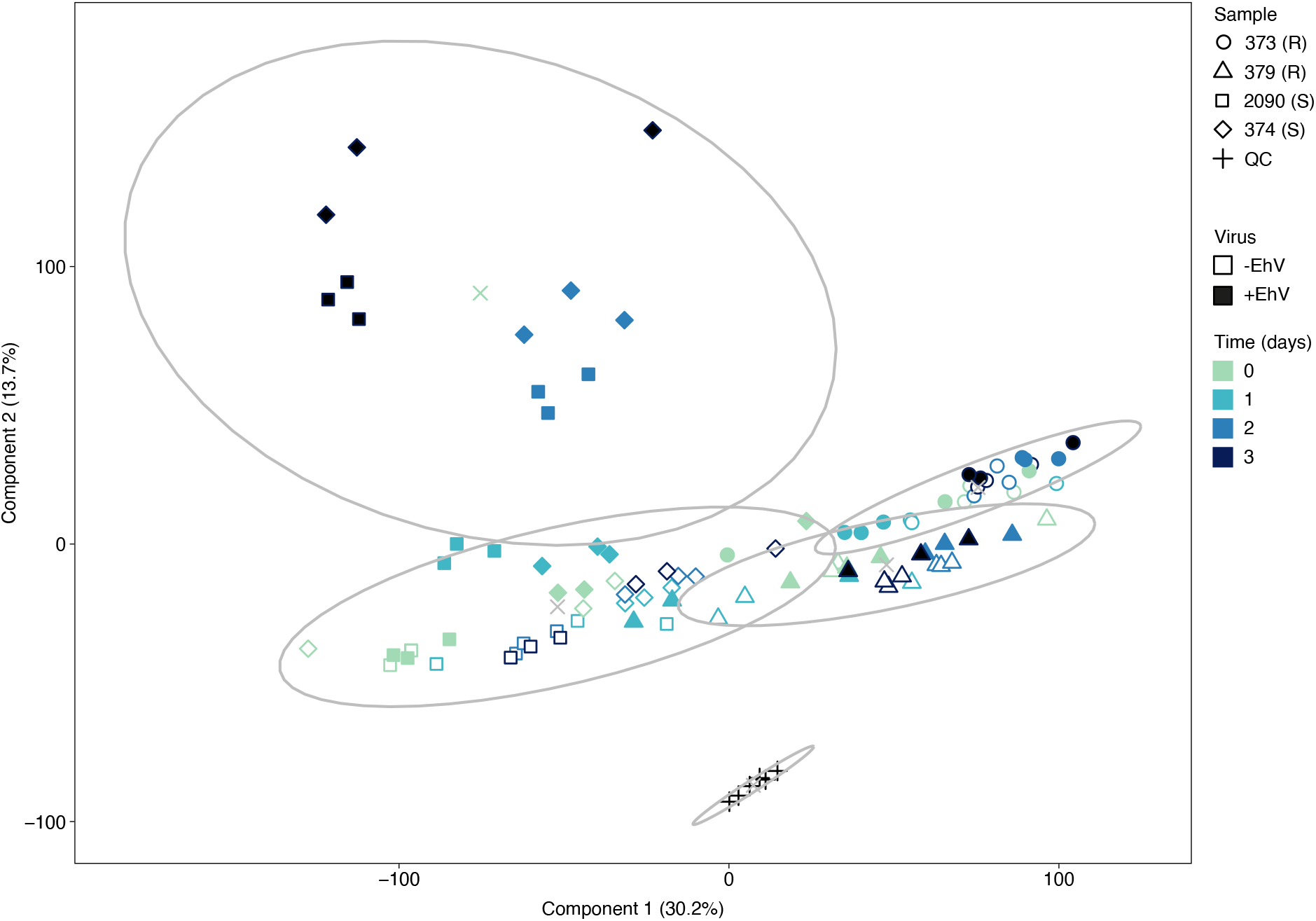
*k*-Means clustering including the pooled QC samples. Clustering of resistant and susceptible *E. huxleyi* strains with and without addition of EhV together with the pooled QC samples based on untargeted lipidomics (using 12,190 mass features) and *k*-means clustering (*k* = 5), as visualized by PCA. Percentage of explained variance is stated in parentheses. Each cluster is surrounded by an ellipse, with the mean marked by ‘×’.

**Figure S36:**
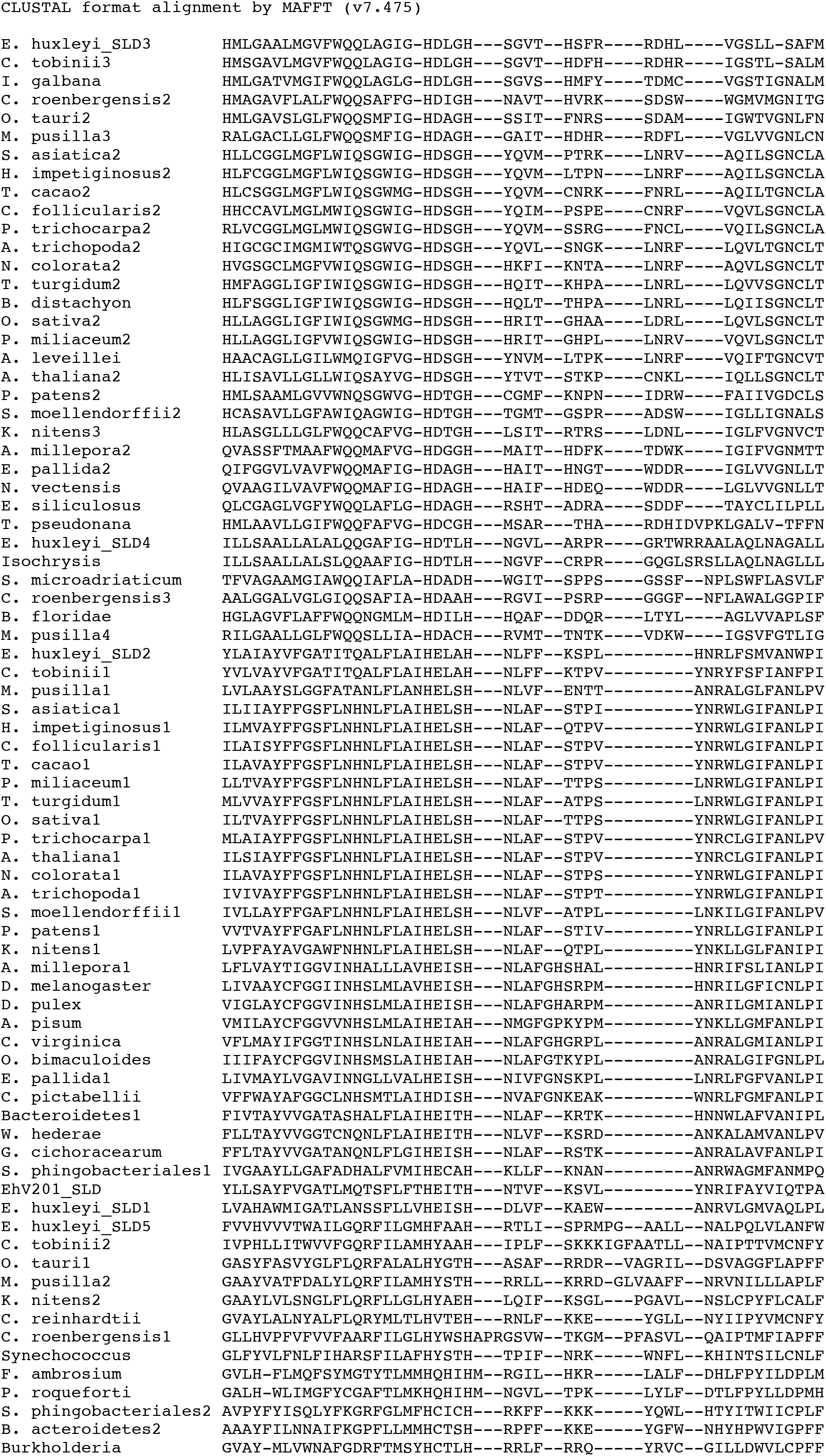
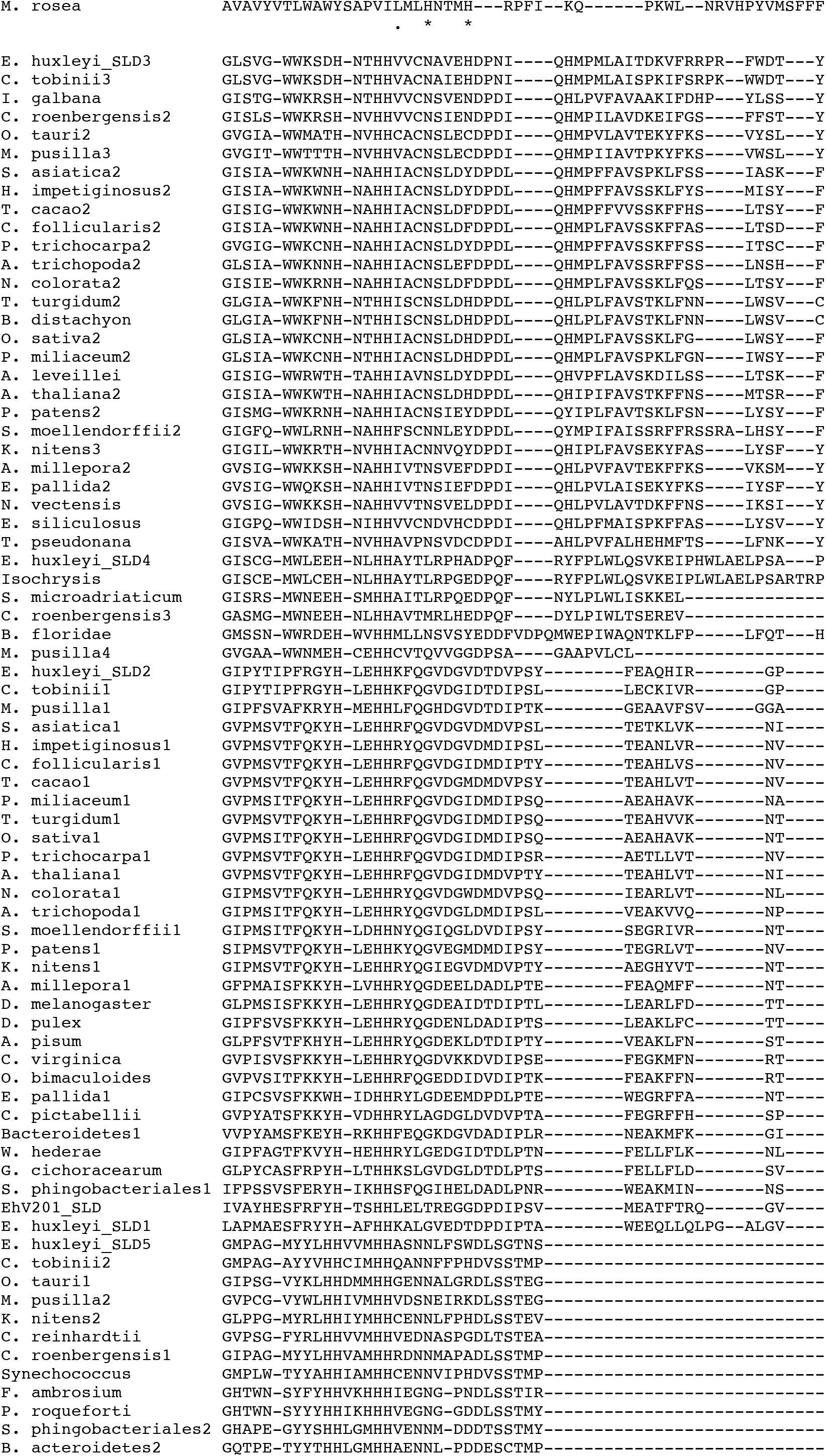
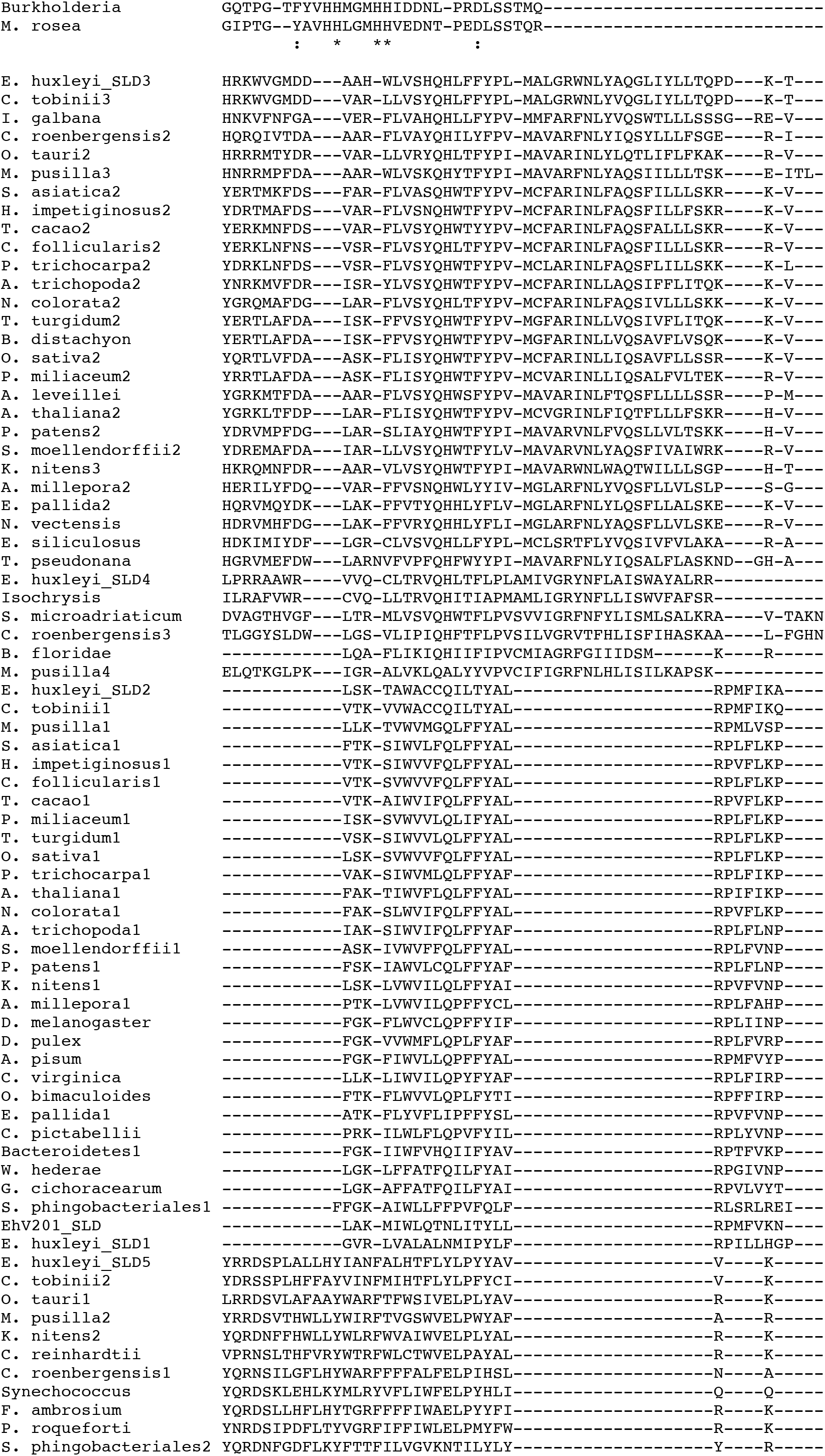
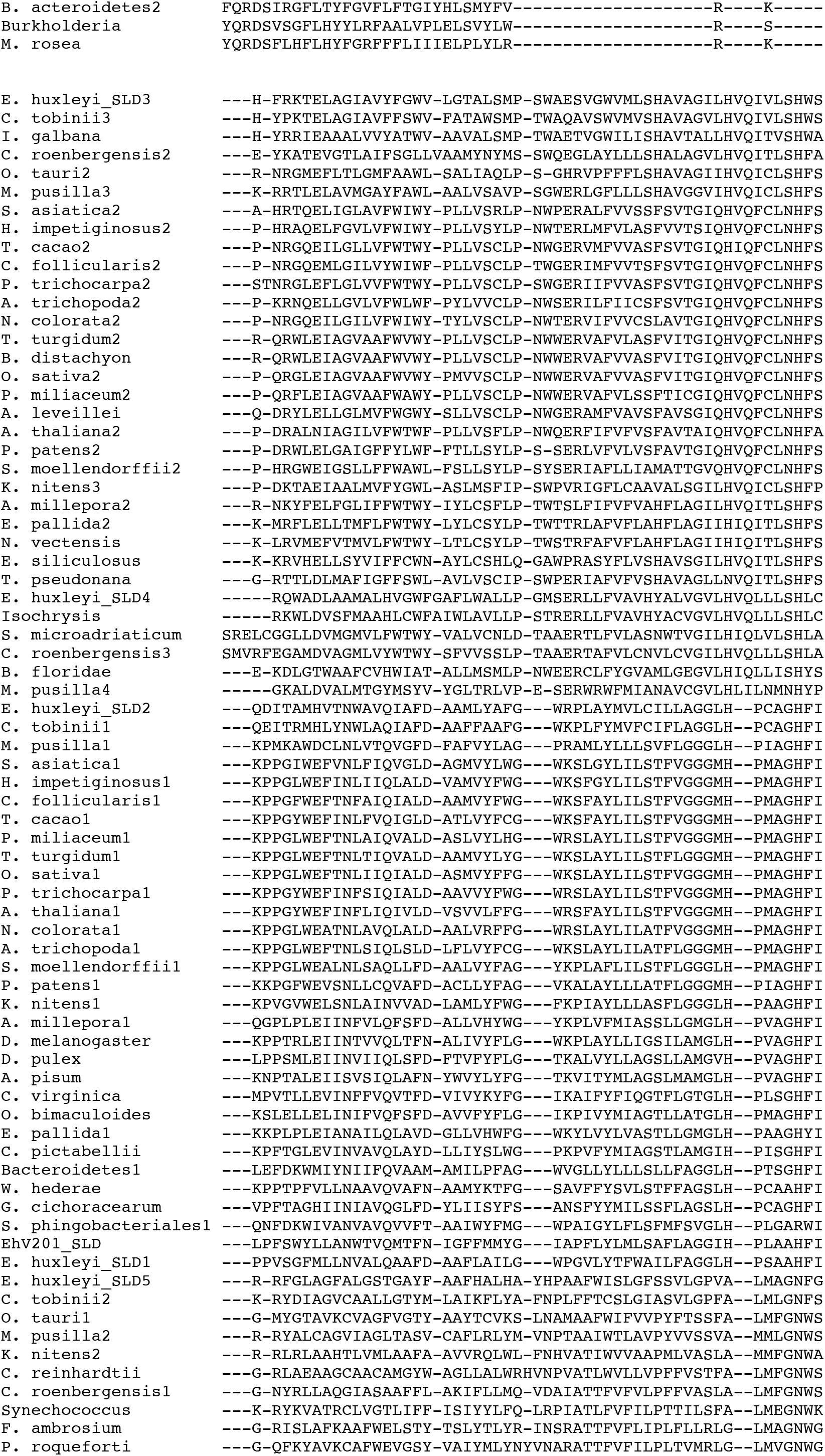
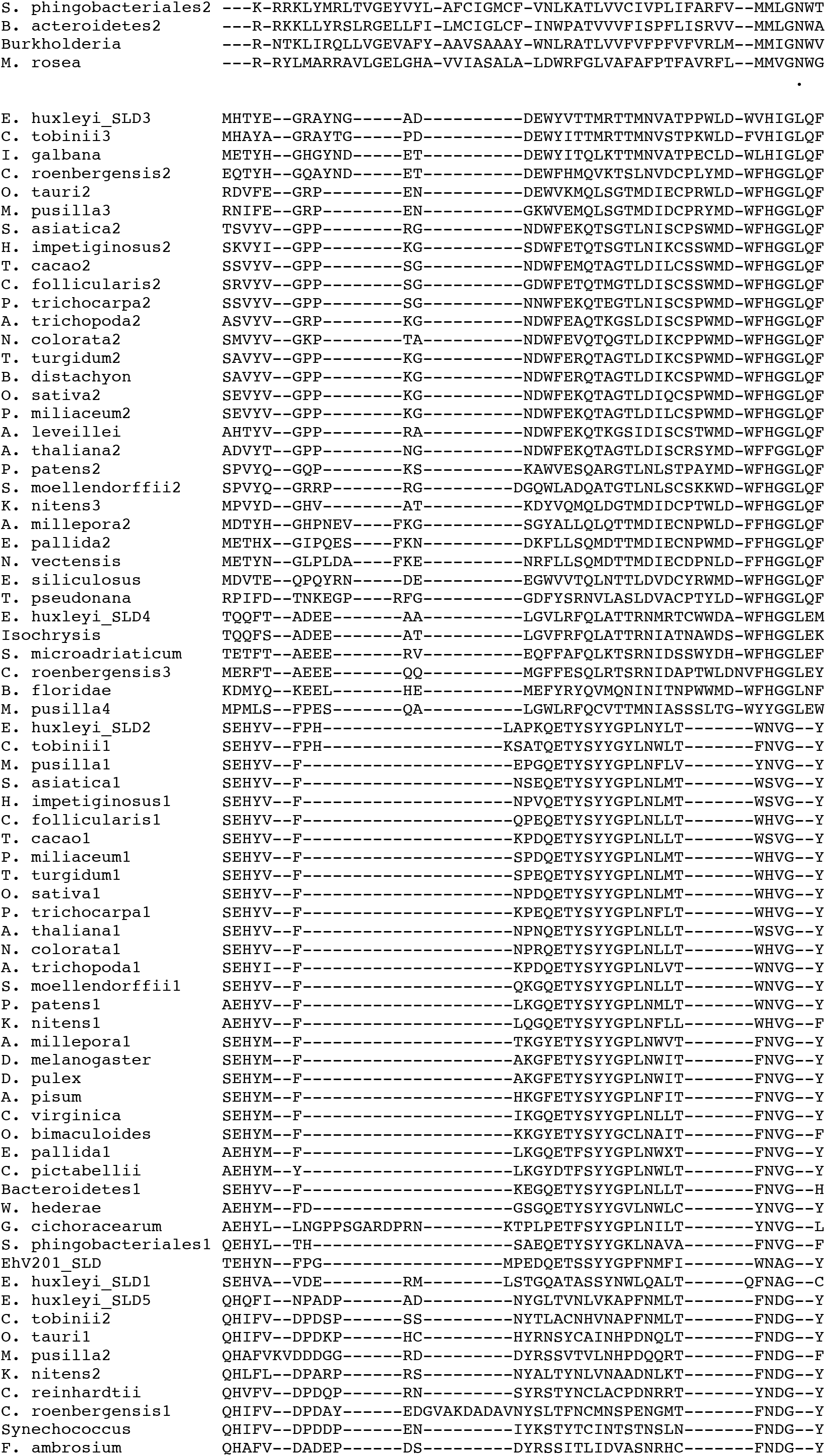
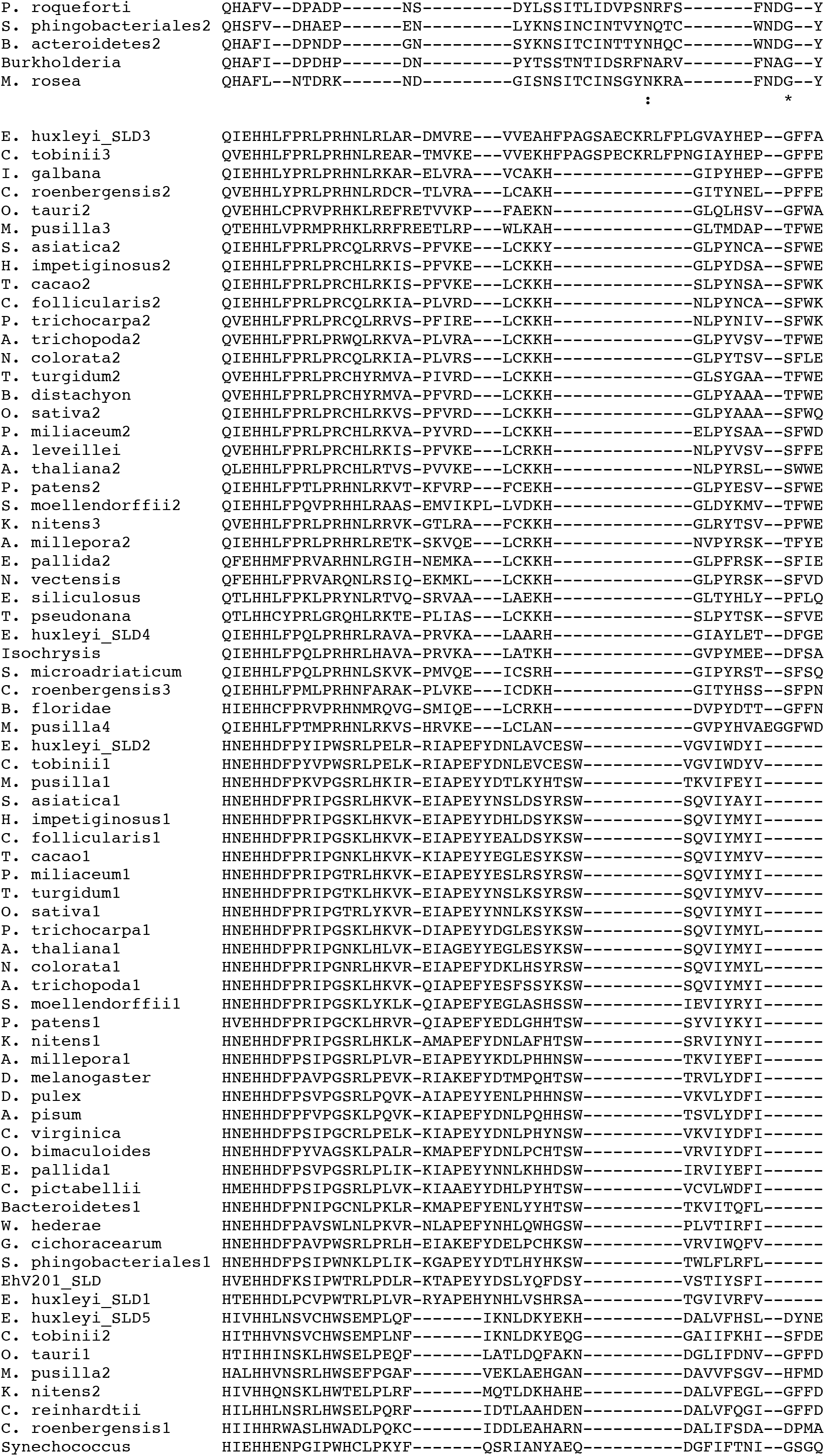
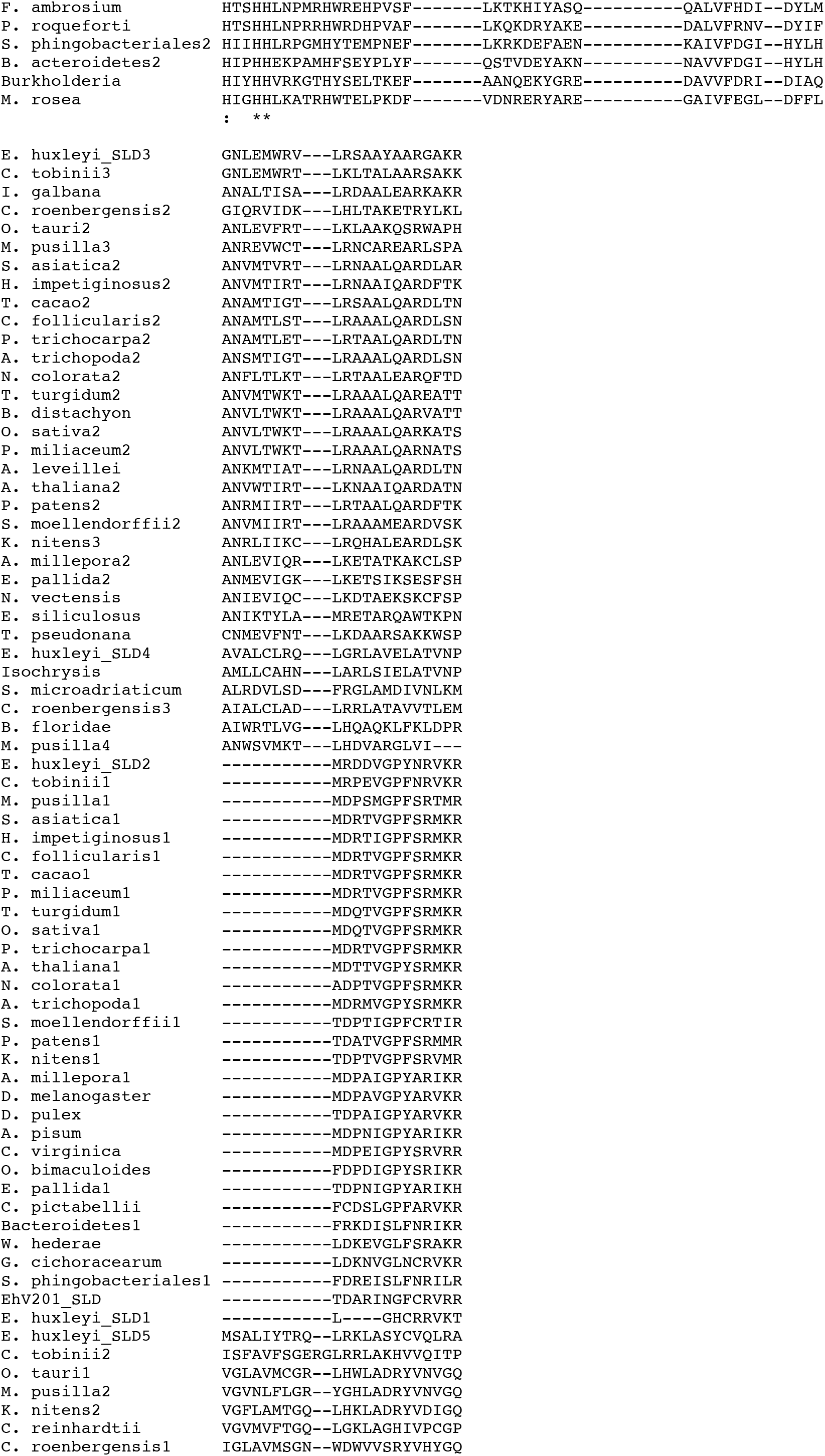
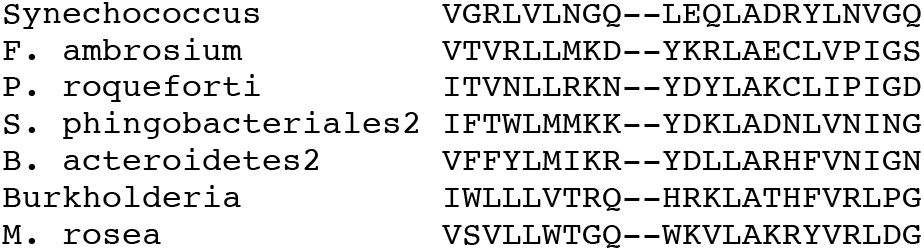
Multiple amino acid sequence alignment of the conserved domain of SLDs. As the subfamilies differ strongly even within the domain, Mafft using the L-INS-i algorithm gave the best alignment (see Methods). Details of the sequences are listed in Table S5. Consensus symbols are as follows: asterisk (*) indicates fully conserved residues, (:) indicates conservation between groups of strongly similar properties, and (.) indicates conservation between groups of weakly similar properties.

**Figure S37:**
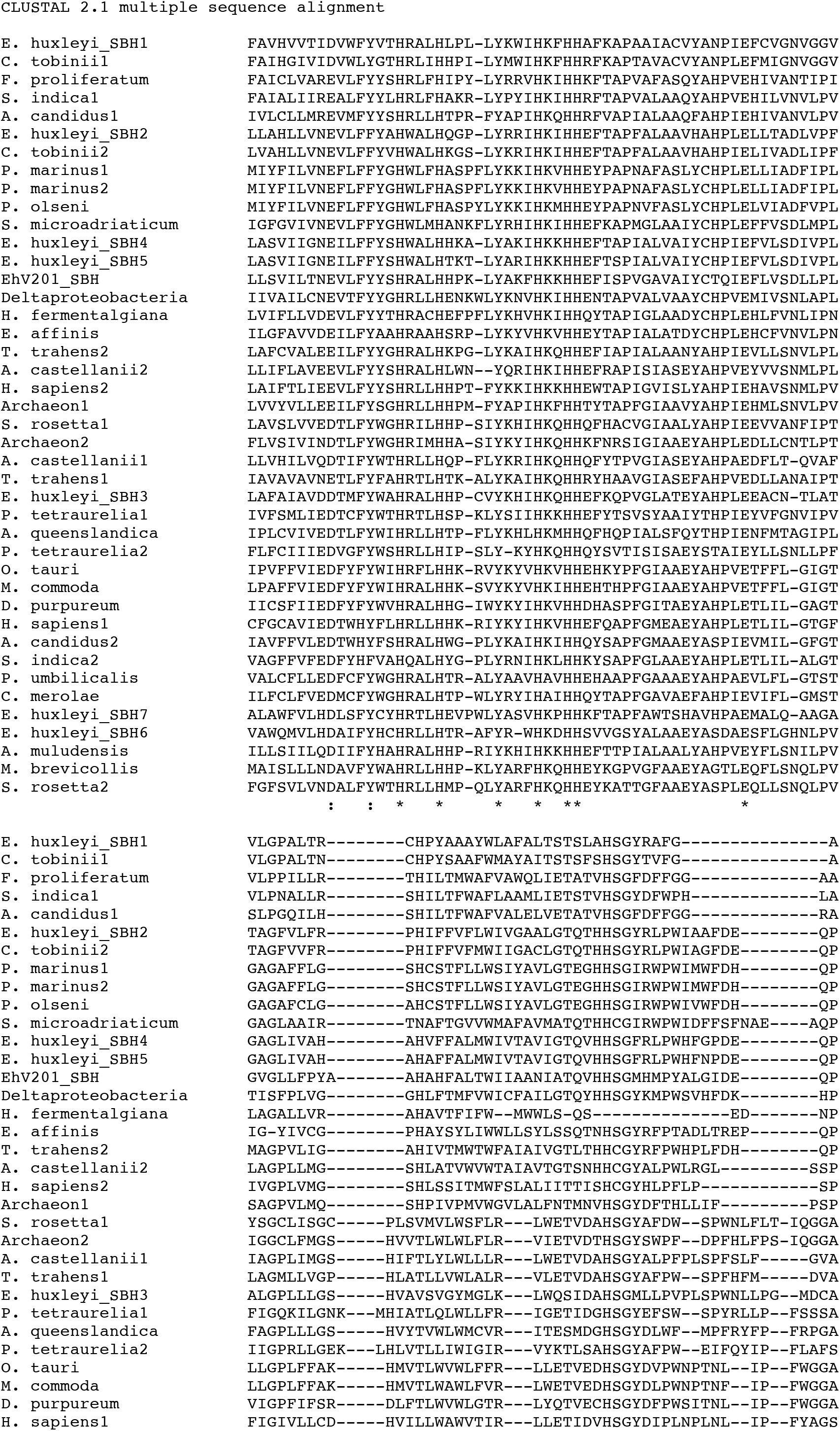
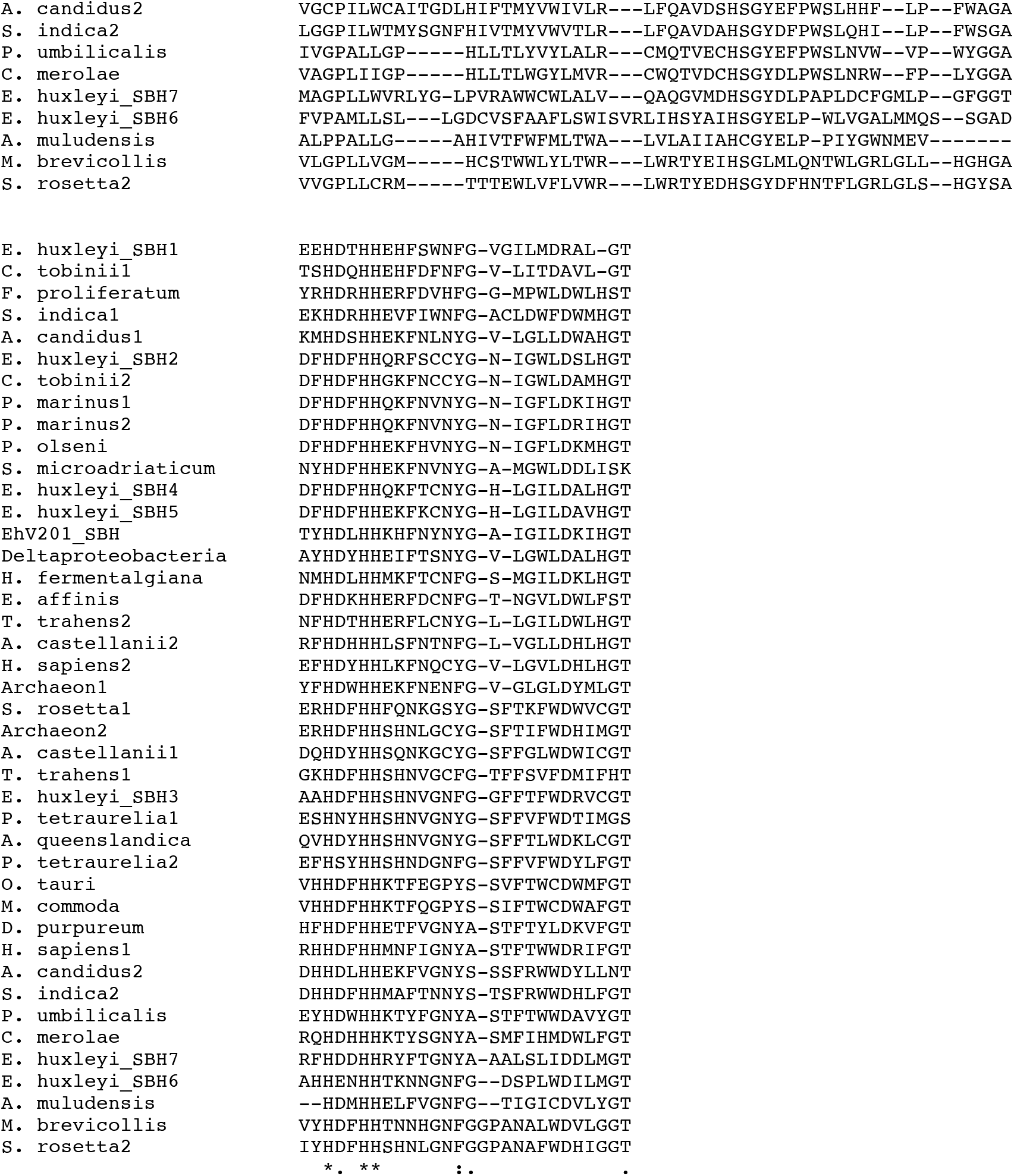
Multiple amino acid sequence alignment of the conserved domain of SBHs. ClustalW alignment of the domain is shown. Details of the sequences are listed in Table S6. Consensus symbols are as follows: asterisk (*) indicates fully conserved residues, (:) indicates conservation between groups of strongly similar properties, and (.) indicates conservation between groups of weakly similar properties.

**Figure S38:**
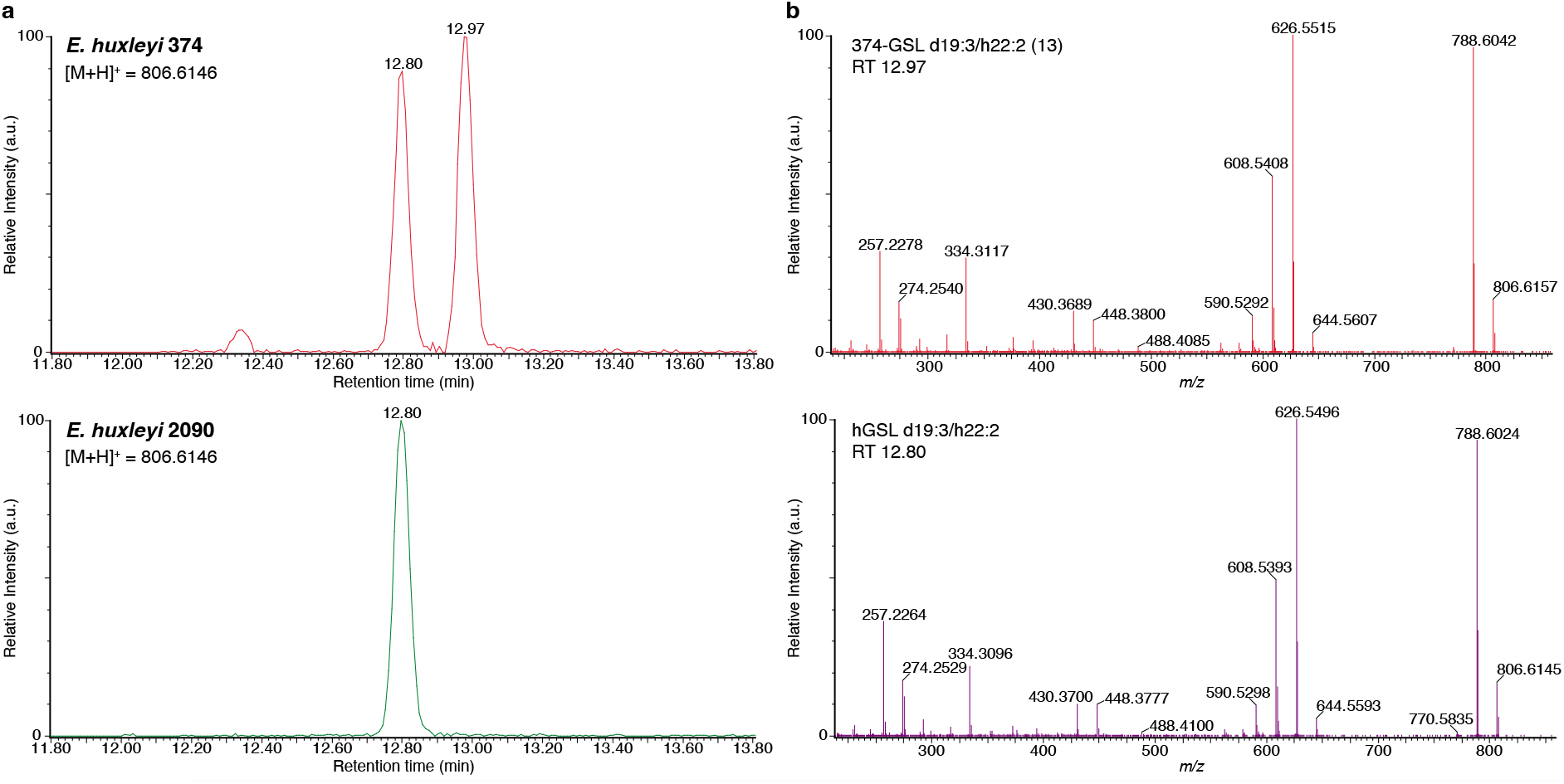
LC-MS/MS analysis of 374-GSL d19:3/h22:2 (13) and hGSL d19:3/h22:2. (**a**) Extracted ion chromatogram (EIC) of *m/z* 806.6146 in *E. huxleyi* strains 374 (top) and 2090 (bottom). hGSL d19:3/h22:2 appears in both strains (RT 12.80 min), while 374-GSL d19:3/h22:2 (**13**) appears only in *E. huxleyi* strain 374 (RT 12.97 min). (**b**) LC-MS/MS spectra of both GSLs show similar fragmentation, suggesting that they are structural isomers, either in the LCB, FA or sugar headgroup.

**Figure S39:**
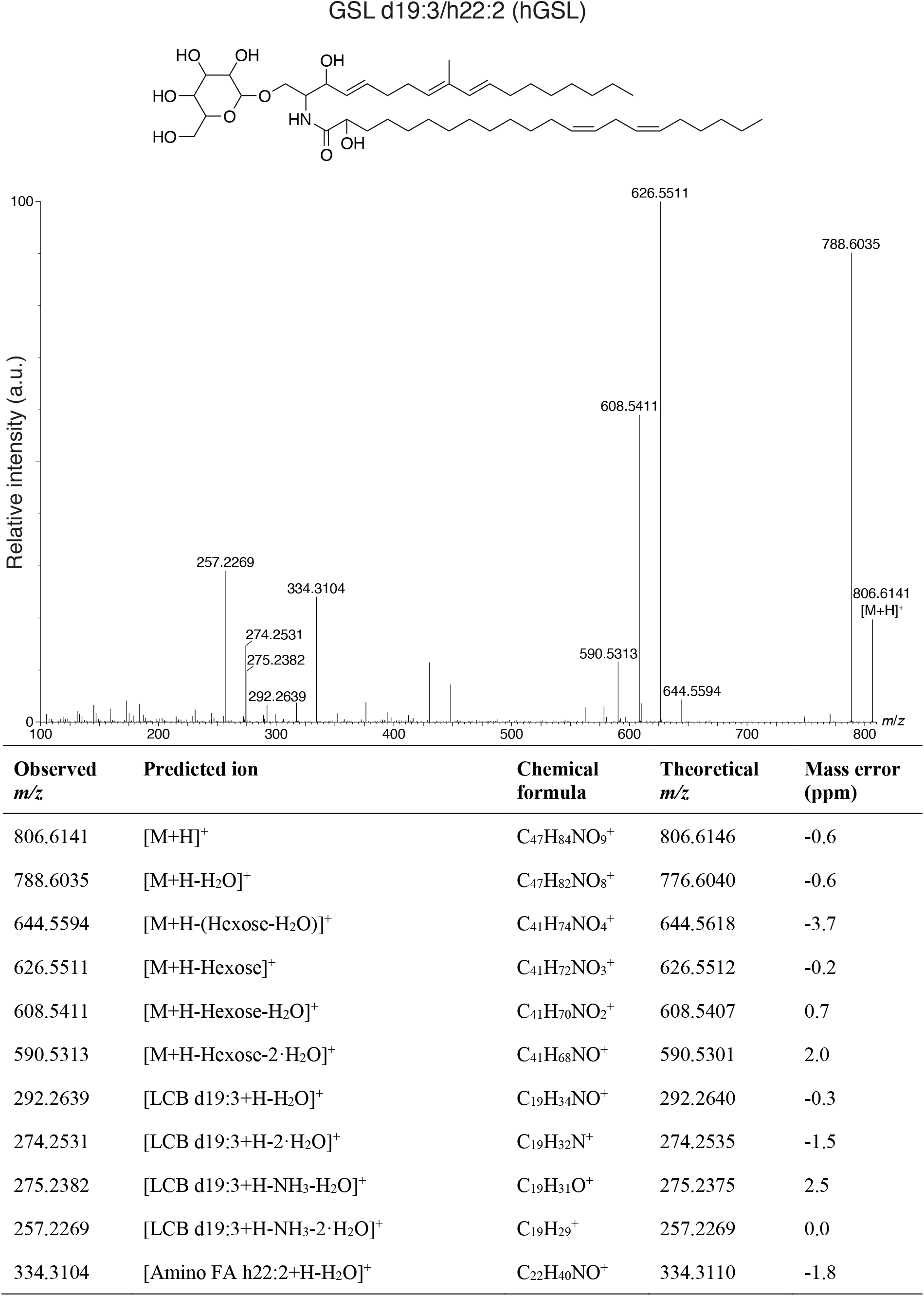
LC-MS/MS analysis of hGSL d19:3/h22:2. A putative structure is presented, as reported previously^6^. The structure is supported by a list of fragments detected in MS/MS mode (Metabolomics Standards Initiative level 2 annotation^2^). Fragments were detected in positive ionization MS/MS mode using [M+H]^+^ = 806.6146 as the precursor ion (Table S3).

**Figure S40:**
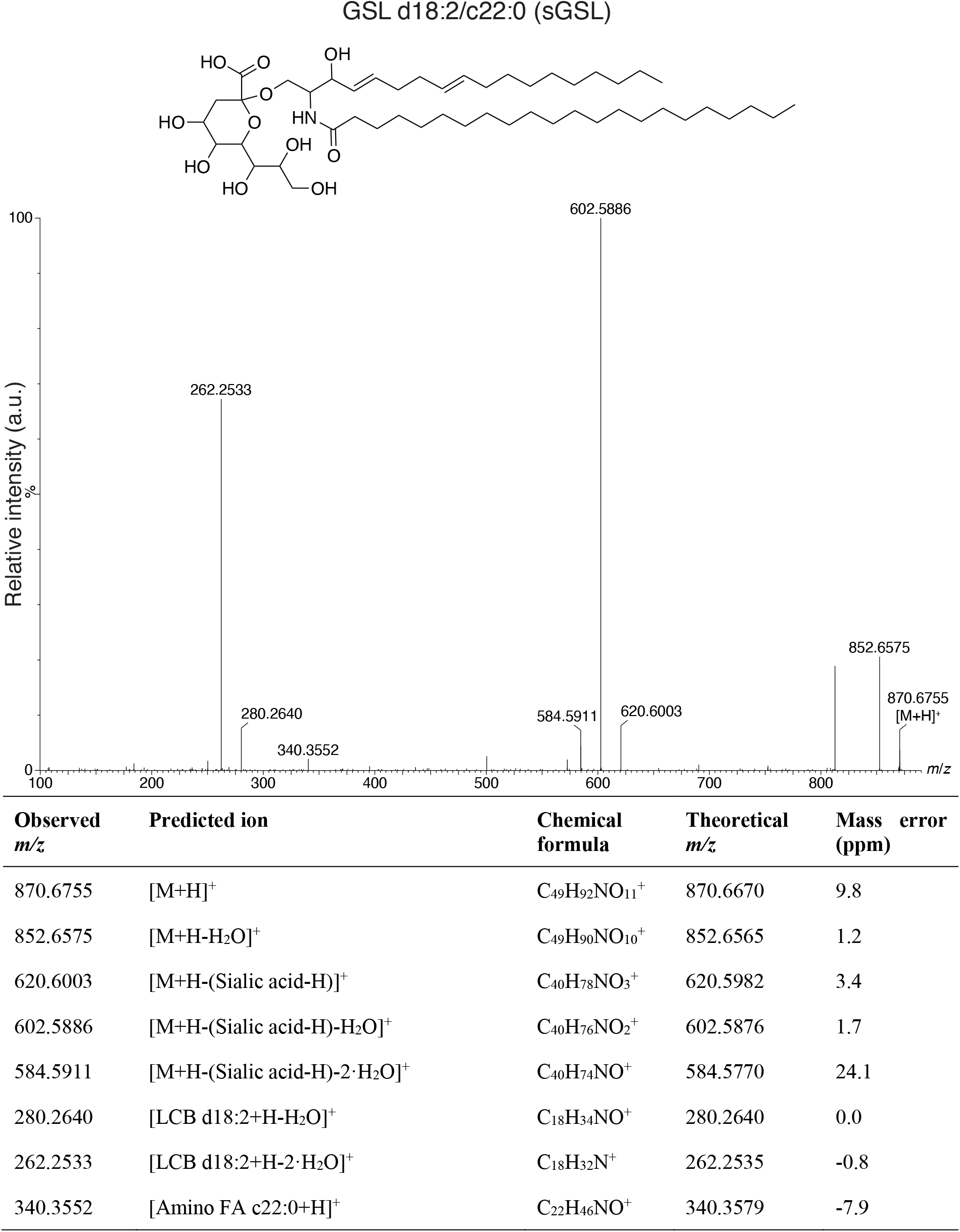
LC-MS/MS analysis of sGSL d18:2/c22:0. A putative structure is presented, as reported previously^7^. The structure is supported by a list of fragments detected in MS/MS mode (Metabolomics Standards Initiative level 2 annotation^2^). Fragments were detected in positive ionization MS/MS mode using [M+H]^+^ = 870.6670 as the precursor ion (Table S3).

**Figure S41:**
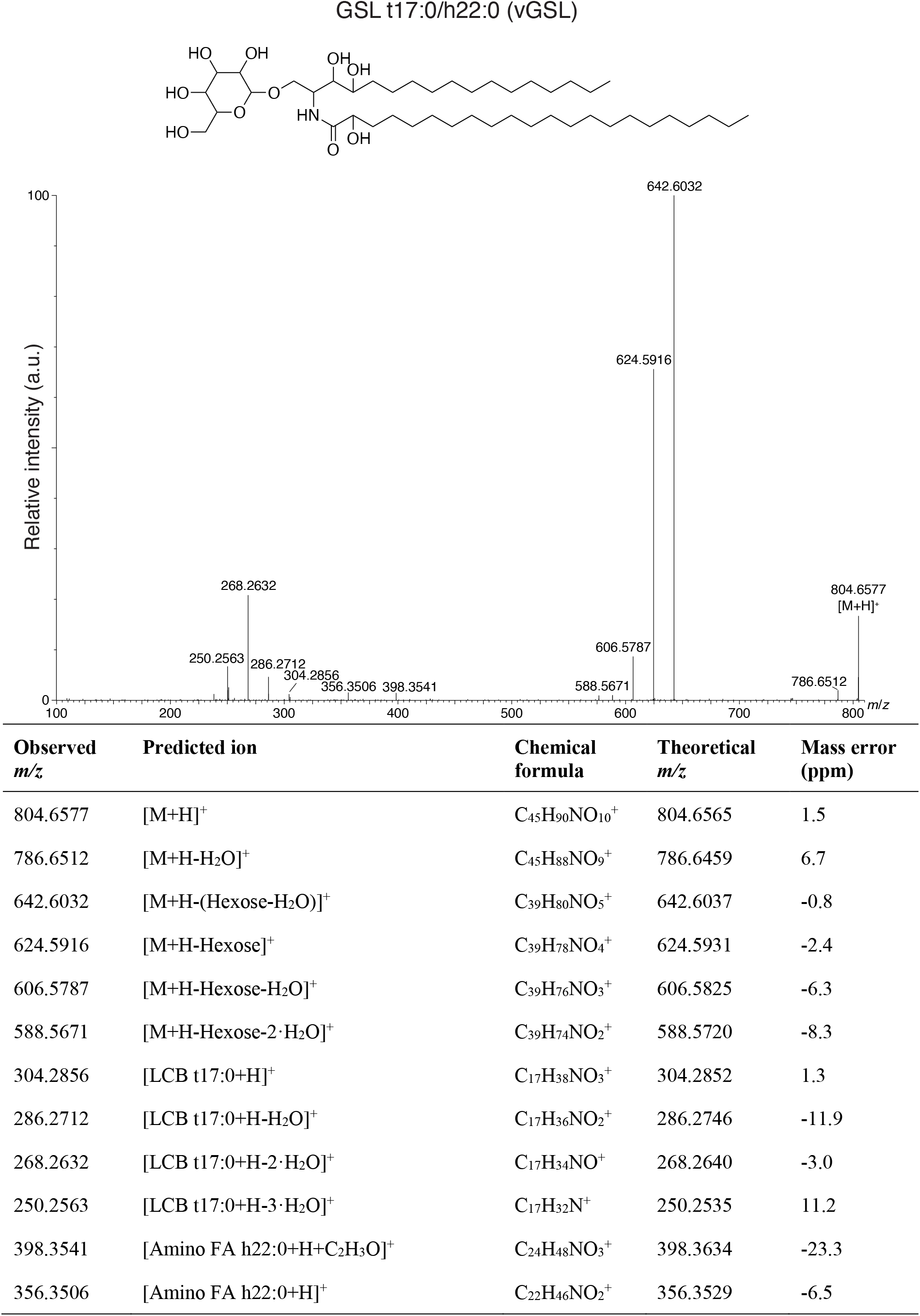
LC-MS/MS analysis of vGSL t17:0/h22:0. A putative structure is presented, as reported previously^8^. The structure is supported by a list of fragments detected in MS/MS mode (Metabolomics Standards Initiative level 2 annotation^2^). Fragments were detected in positive ionization MS/MS mode using [M+H]^+^ = 804.6565 as the precursor ion (Table S3).

**Table S1:**
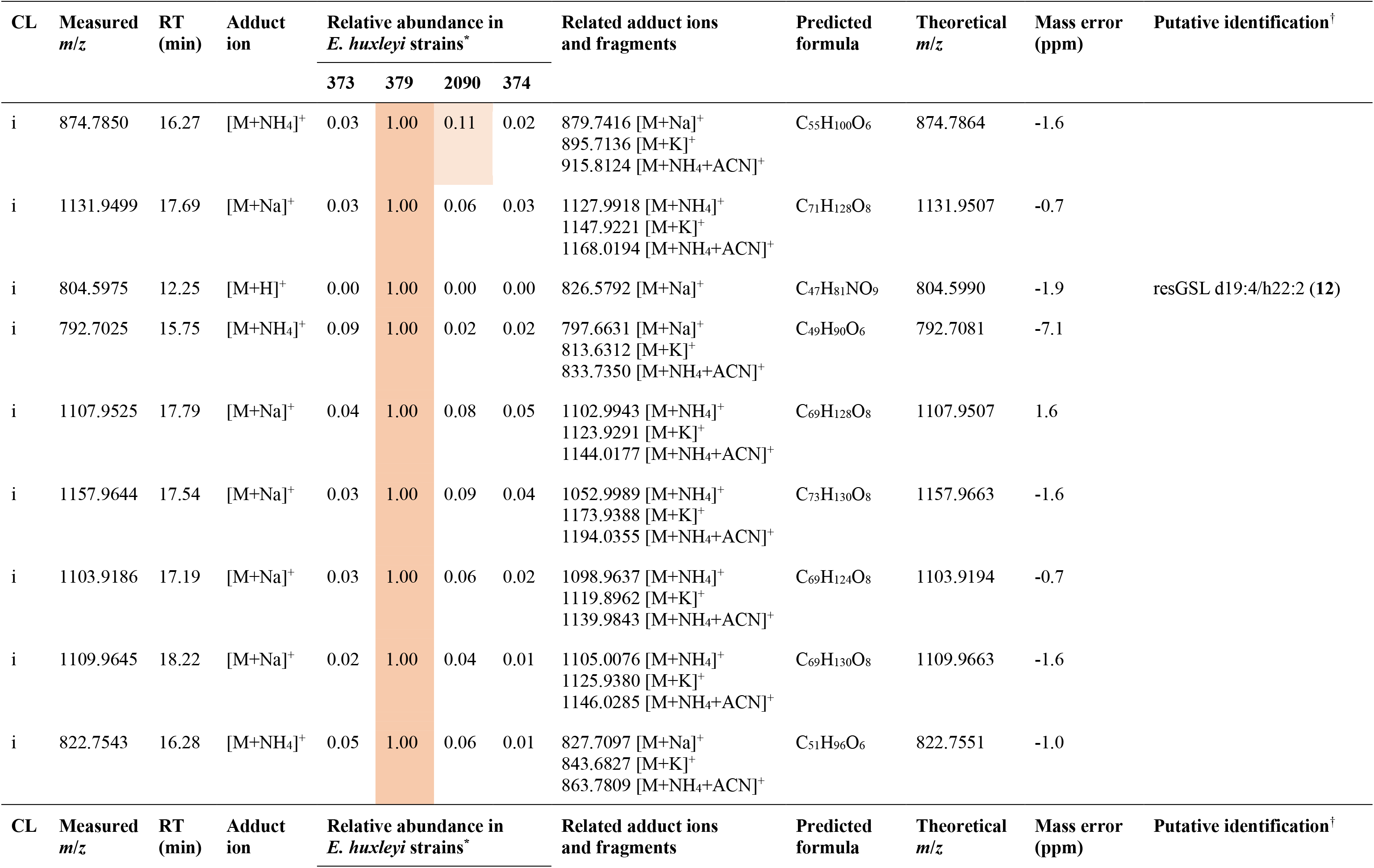
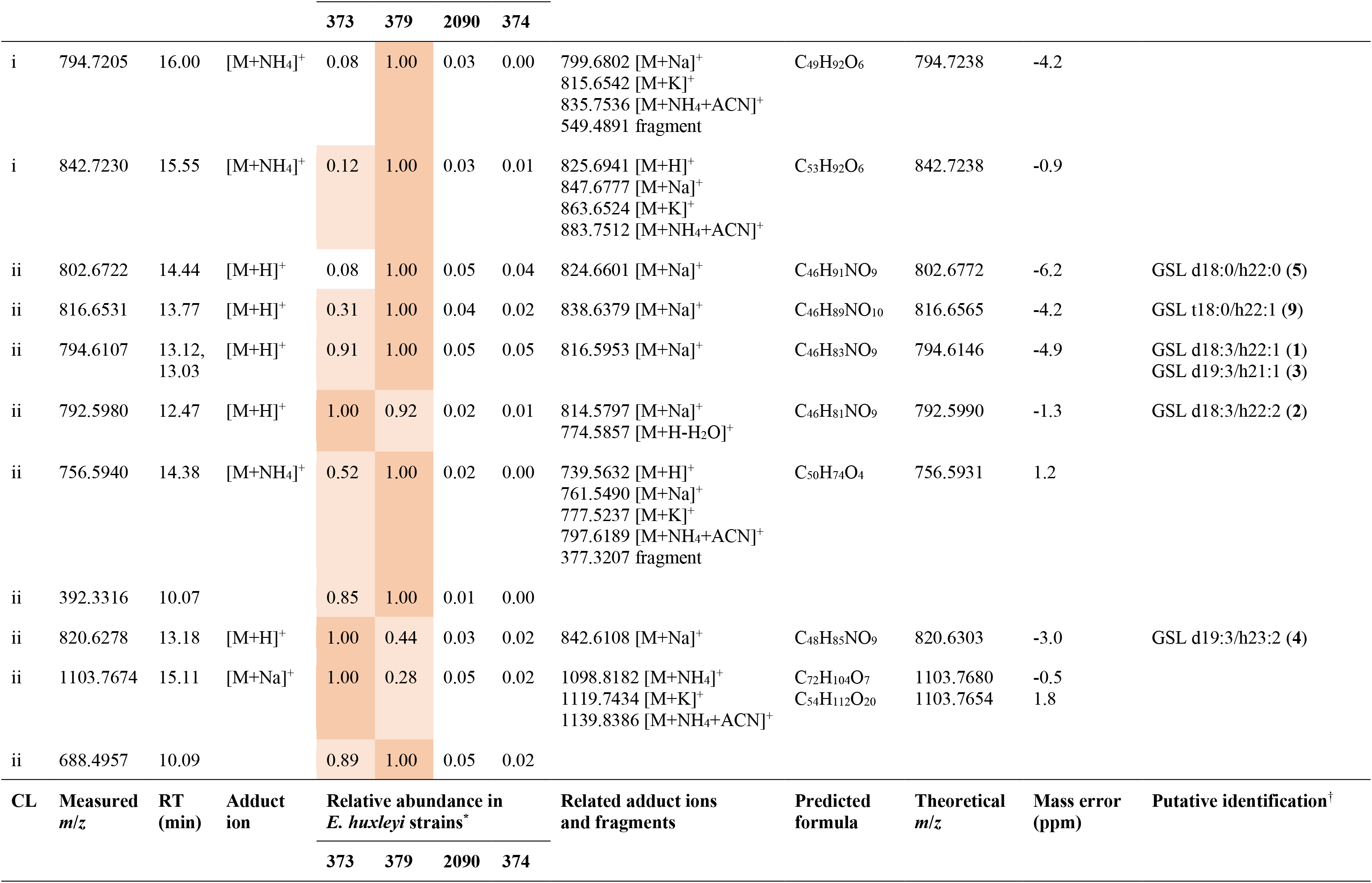
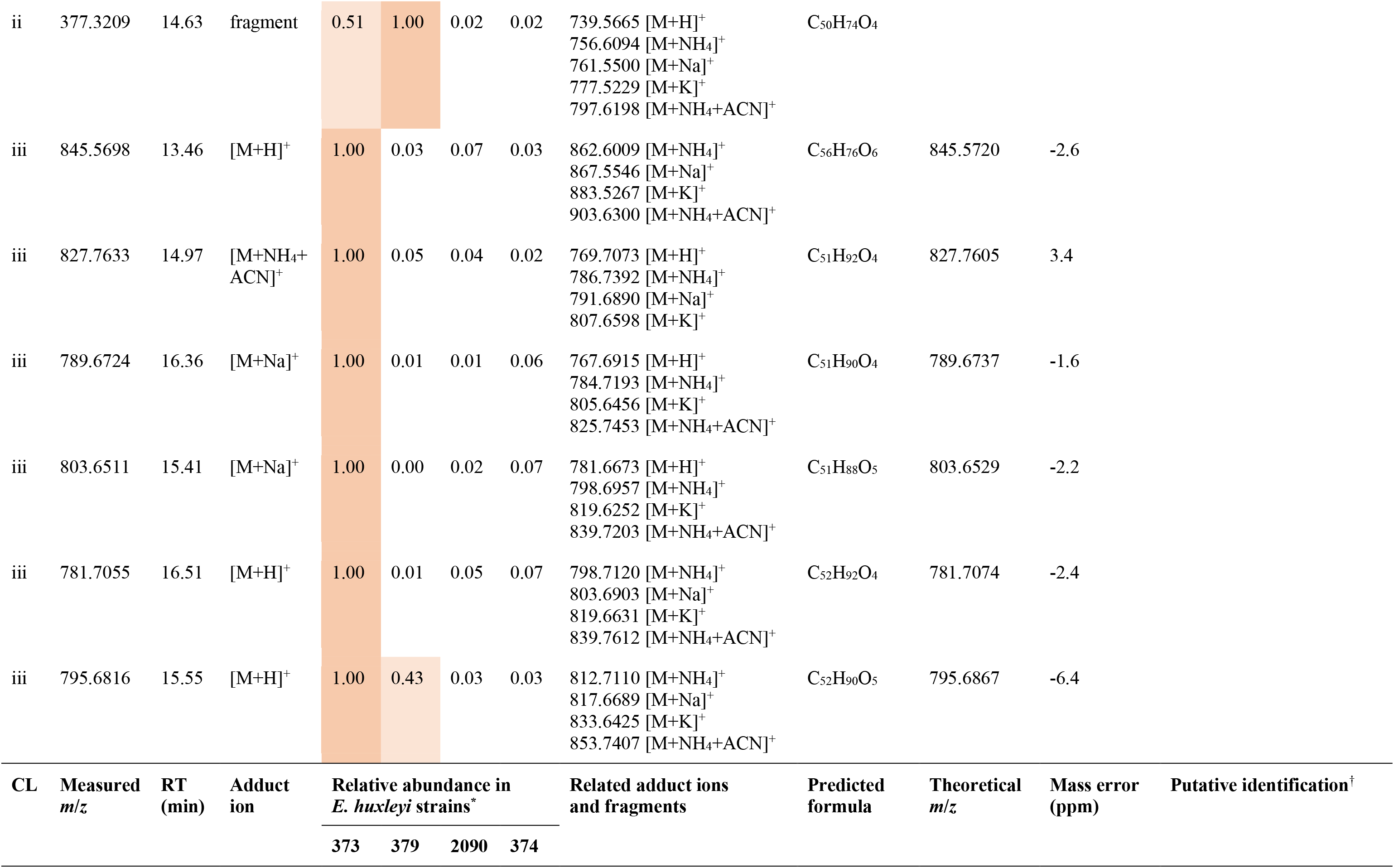
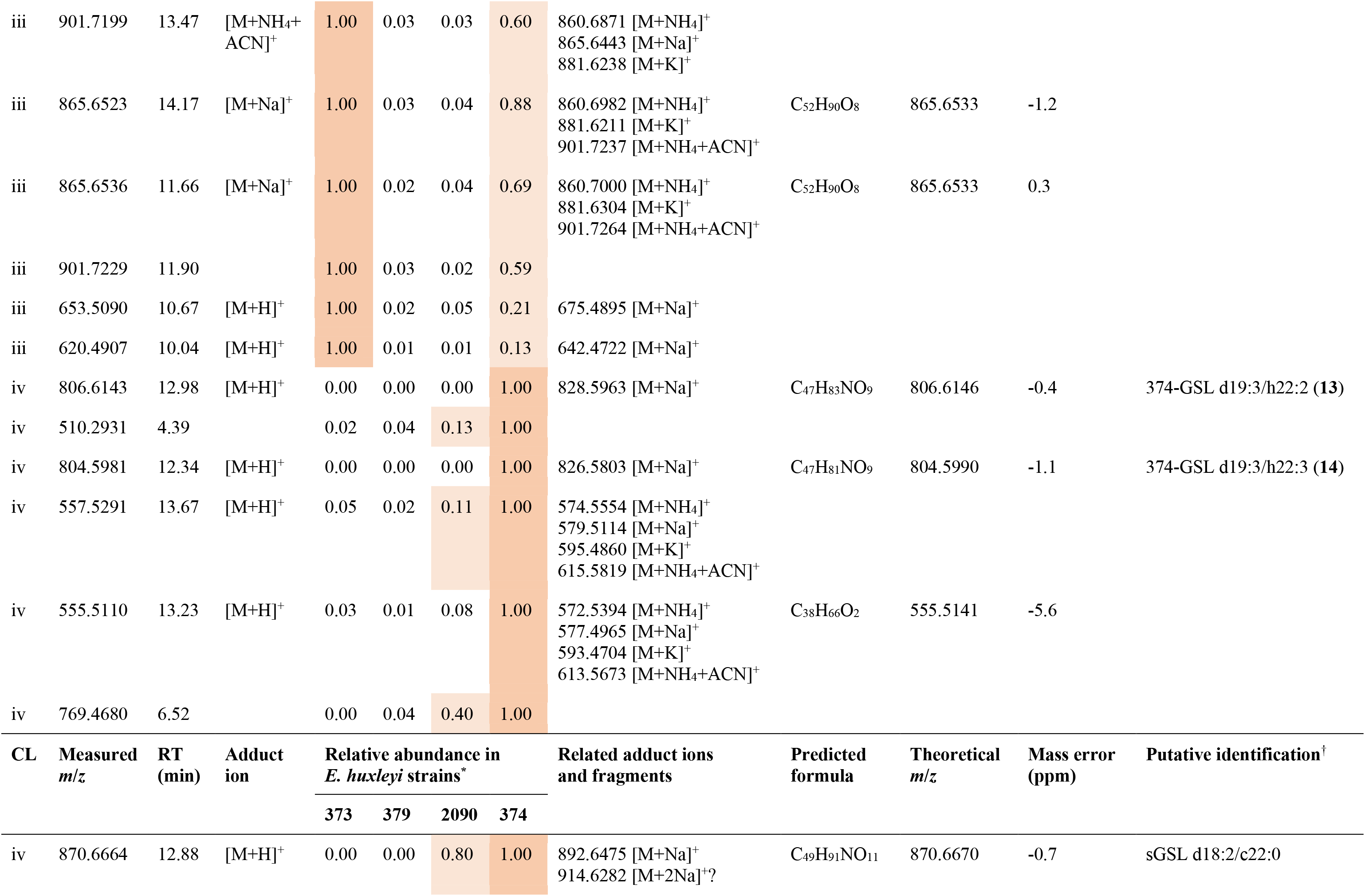
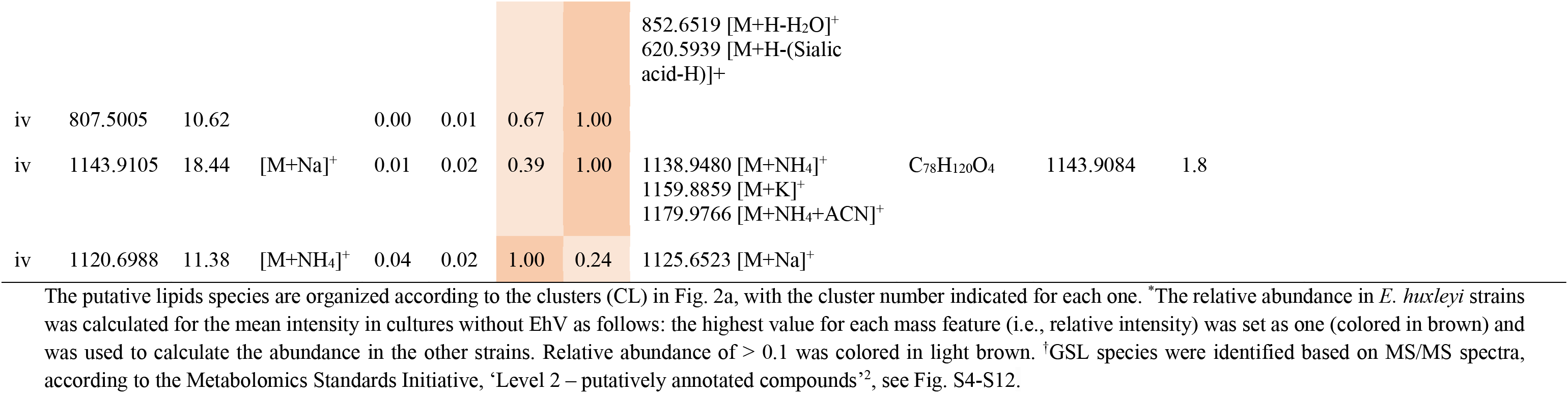
Putative lipid biomarkers for susceptible and resistant *E. huxleyi* cells.

**Table S2:** Additional putatively annotated GSLs species.

| # | GSL species<br>LCB/FA | Measured $m/z$<br>([M+H]<sup>+</sup>) | RT<br>(min) | Predicted<br>formula | Theoretical $m/z$<br>([M+H]<sup>+</sup>) | Mass error<br>(ppm) |
| --- | --- | --- | --- | --- | --- | --- |
| 6 | d18:0/h22:1 | 800.6600 | 14.22 | C<sub>46</sub>H<sub>89</sub>NO<sub>9</sub> | 800.6616 | -2.0 |
| 7 | d18:1/h22:1 | 798.6440 | 14.01 | C<sub>46</sub>H<sub>87</sub>NO<sub>9</sub> | 798.6459 | -2.4 |
| 8 | t18:0/h22:0 | 818.6702 | 14.00 | C<sub>46</sub>H<sub>91</sub>NO<sub>10</sub> | 818.6721 | -2.3 |
| 10 | t18:0/h22:2 | 814.6346 | 13.18 | C<sub>46</sub>H<sub>87</sub>NO<sub>10</sub> | 814.6408 | -7.6 |
| 11 | d19:4/h22:1<br>(resGSL) | 806.6127 | 12.92 | C<sub>47</sub>H<sub>83</sub>NO<sub>9</sub> | 806.6146 | -2.4 |

**Table S3:** GSL species identified in the *E. huxleyi*-EhV model system.

| Head group | LCB/FA composition | Chemical formula | RT (min) | Theoretical $m/z$ ($[M+H]^+$) | Name in the *E. huxleyi*-EhV system | Common name* (LIPID MAPS ID) | Occurrence | Reference |
| --- | --- | --- | --- | --- | --- | --- | --- | --- |
| Hexose | d18:0/h22:0 | C<sub>46</sub>H<sub>91</sub>NO<sub>9</sub> | 14.44 | 802.6772 | Group B (5) | HexCer(d18:0/22:0(2OH)) | Resistant cells, infected cells | This study |
| Hexose | d18:0/h22:1 | C<sub>46</sub>H<sub>89</sub>NO<sub>9</sub> | 14.22 | 800.6616 | Group B (6) | HexCer(d18:0/22:1(2OH)) | Resistant cells, infected cells | This study |
| Hexose | d18:1/h22:1 | C<sub>46</sub>H<sub>87</sub>NO<sub>9</sub> | 14.01 | 798.6459 | Group B (7) | HexCer(d18:1/22:1(2OH)) | Resistant cells, infected cells | This study |
| Sialic acid | d18:1/c22:0 | C<sub>49</sub>H<sub>93</sub>NO<sub>11</sub> | 12.88 | 872.6827 | sGSL |  | Susceptible strains, infected cells | Fulton et al., 2014 |
| Sialic acid | d18:2/c22:0 | C<sub>49</sub>H<sub>91</sub>NO<sub>11</sub> | 13.25 | 870.6670 | sGSL |  | Susceptible strains, infected cells | Fulton *et al.*, 2014 |
| Hexose | d18:3/h22:1 | C<sub>46</sub>H<sub>83</sub>NO<sub>9</sub> | 13.12 | 794.6146 | Group A (1) | HexCer(d18:3/22:1(2OH)) | All cell types | This study |
| Hexose | d18:3/h22:2 | C<sub>46</sub>H<sub>81</sub>NO<sub>9</sub> | 12.47 | 792.5990 | Group A (2) | HexCer(d18:3/22:2(2OH)) | All cell types | This study |
| Hexose | d19:3/h21:1 | C<sub>46</sub>H<sub>83</sub>NO<sub>9</sub> | 13.03 | 794.6146 | Group A (3) | HexCer(d19:3/21:1(2OH)) | All cell types | This study |
| Hexose | d19:3/h22:1 | C<sub>47</sub>H<sub>85</sub>NO<sub>9</sub> | 13.41 | 808.6303 | hGSL | HexCer(d19:3/22:1(2OH)) | All cell types | Vardi *et al.*, 2012 |
| Hexose | d19:3/h22:2 | C<sub>47</sub>H<sub>83</sub>NO<sub>9</sub> | 12.80 | 806.6146 | hGSL | HexCer(d19:3/22:2(2OH)) | All cell types | Vardi *et al.*, 2012 |
| Hexose | d19:3/h22:2 | C<sub>47</sub>H<sub>83</sub>NO<sub>9</sub> | 12.98 | 806.6146 | 374-GSL (13) | HexCer(d19:3/22:2(2OH)) | Susceptible cells, not all strains | This study |
| Hexose | d19:3/h22:3 | C<sub>47</sub>H<sub>81</sub>NO<sub>9</sub> | 12.34 | 804.5990 | 374-GSL (14) | HexCer(d19:3/22:3(2OH)) | Susceptible cells, not all strains | This study |
| Hexose | d19:3/h23:2 | C<sub>48</sub>H<sub>85</sub>NO<sub>9</sub> | 13.18 | 820.6303 | Group A (4) | HexCer(d19:3/23:2(2OH)) | All cell types | This study |
| Hexose | d19:4/h22:1 | C<sub>47</sub>H<sub>83</sub>NO<sub>9</sub> | 12.92 | 806.6146 | resGSL (11) | HexCer(d19:4/22:1(2OH)) | Resistant cells | This study |
| Hexose | d19:4/h22:2 | C<sub>47</sub>H<sub>81</sub>NO<sub>9</sub> | 12.25 | 804.5990 | resGSL (12) | HexCer(d19:4/22:2(2OH)) | Resistant cells | This study |
| Hexose | t16:0/h22:0 | C<sub>44</sub>H<sub>87</sub>NO<sub>10</sub> | 13.36 | 790.6408 | vGSL | HexCer(t16:0/22:0(2OH)) | Infected cells | Schleyer *et al.*, 2019 |
| Hexose | t17:0/h22:0 | C<sub>45</sub>H<sub>89</sub>NO<sub>10</sub> | 13.69 | 804.6565 | vGSL | HexCer(t17:0/22:0(2OH))<br>(LMSP05010197) | Infected cells | Vardi *et al.*, 2012<br>Ziv *et al.*, 2016 |
| Hexose | t17:0/h22:1 | C<sub>45</sub>H<sub>87</sub>NO<sub>10</sub> | 13.07 | 802.6408 | vGSL | HexCer(t17:0/22:1(2OH)) | Infected cells | Vardi *et al.*, 2012<br>Ziv *et al.*, 2016 |
| Hexose | t17:0/h23:0 | C<sub>46</sub>H<sub>91</sub>NO<sub>10</sub> | 14.02 | 818.6721 | vGSL | HexCer(t17:0/23:0(2OH)) | Infected cells | Vardi *et al.*, 2012<br>Ziv *et al.*, 2016 |
| Hexose | t17:0/h23:1 | C<sub>46</sub>H<sub>89</sub>NO<sub>10</sub> | 13.45 | 816.6565 | vGSL | HexCer(t17:0/23:1(2OH)) | Infected cells | Vardi *et al.*, 2012<br>Ziv *et al.*, 2016 |
| Hexose | t17:0/h24:0 | C<sub>47</sub>H<sub>93</sub>NO<sub>10</sub> | 14.33 | 832.6878 | vGSL | HexCer(t17:0/24:0(2OH))<br>(LMSP05010196) | Infected cells | Vardi *et al.*, 2012<br>Ziv *et al.*, 2016 |
| Hexose | t17:0/h24:1 | C<sub>47</sub>H<sub>91</sub>NO<sub>10</sub> | 13.80 | 830.6721 | vGSL | HexCer(t17:0/24:1(2OH)) | Infected cells | Vardi *et al.*, 2012<br>Ziv *et al.*, 2016 |
| Hexose | t18:0/h22:0 | C<sub>46</sub>H<sub>91</sub>NO<sub>10</sub> | 14.00 | 818.6721 | Group B / vGSL (8) | HexCer(t18:0/22:0(2OH)) | Resistant cells,<br>Infected cells | This study,<br>Ziv *et al.*, 2016 |
| Hexose | t18:0/h22:1 | C<sub>46</sub>H<sub>89</sub>NO<sub>10</sub> | 13.77 | 816.6565 | Group B / vGSL (9) | HexCer(t18:0/22:1(2OH)) | Resistant cells,<br>Infected cells | This study |
| Hexose | t18:0/h22:2 | C<sub>46</sub>H<sub>87</sub>NO<sub>10</sub> | 13.18 | 814.6408 | Group B / vGSL (10) | HexCer(t18:0/22:2(2OH)) | Resistant cells,<br>Infected cells | This study |

\*Common name is based on the LIPID MAPS classification system. LCB, long-chain base; FA, fatty acid; RT, retention time.

**Table S4:** Genes names and accession numbers.

| Name | Accession |
| --- | --- |
| *sld1* | MZ152812 |
| *sld2* | MZ152813, MZ152814 (KJ868223, previously called *dcd2*) |
| *sld3* | MZ152815 |
| *sld4* | MZ152816 |
| *sld5* | MZ152817, MZ152818 |
| *sbh1* | MZ152819, MZ152820 (KJ868226, previously called sphinganine hydroxylase 1) |
| *sbh2* | MZ152821 |
| *sbh3* | Predicted from the genome of *E. huxleyi* CCMP1516 |
| *sbh4* | MZ152822 |
| *sbh5* | MZ152823 |
| *sbh6* | MZ152824, MZ152825 |
| *sbh7* | MZ152826, MZ152827 |

Accession numbers in brackets are of genes that were deposited in GenBank from our earlier definitions<sup>5</sup>.

**Table S5:** Information regarding the proteins used to build the SLD phylogenetic tree.

| Name | Organism | Accession number | Description |
| --- | --- | --- | --- |
| E. huxleyi SLD1 | *Emiliana huxleyi* CCMP373 | MZ152812 | Sphingolipid desaturase 1 |
| E. huxleyi SLD2 | *Emiliana huxleyi* CCMP2090, CCMP373 | MZ152813, MZ152814 | Sphingolipid desaturase 2 |
| E. huxleyi SLD3 | *Emiliana huxleyi* CCMP2090 | MZ152815 | Sphingolipid desaturase 3 |
| E. huxleyi SLD4 | *Emiliana huxleyi* CCMP373 | MZ152816 | Sphingolipid desaturase 4 |
| E. huxleyi SLD5 | *Emiliana huxleyi* CCMP2090, CCMP373 | MZ152817, MZ152818 | Sphingolipid desaturase 5 |
| EhV201 SLD | *Emiliana huxleyi* virus 201 | AET97947.1 | Fatty acid desaturase |
| A. leveillei | *Anemone leveillei* | AAQ10732.1 | Delta-8-sphingolipid desaturase |
| A. millepora1 | *Acropora millepora* | XP_029201914.1 | Sphingolipid delta(4)-desaturase DES1-like |
| A. millepora2 | *Acropora millepora* | XP_029197704.1 | Delta(8)-fatty-acid desaturase 2-like |
| A. pisum | *Acyrtosiphon pisum* | NP_001155533.1 | Sphingolipid delta(4)-desaturase DES1-like |
| A. thaliana1<sup>†</sup> | *Arabidopsis thaliana* | NP_192402.1 | Fatty acid desaturase family protein |
| A. thaliana2 | *Arabidopsis thaliana* | OAP10850.1 | SLD2 |
| A. trichopoda1 | *Amborella trichopoda* | XP_011625523.1 | Sphingolipid delta(4)-desaturase DES1-like |
| A. trichopoda2 | *Amborella trichopoda* | XP_006847040.1 | Acyl-lipid (9-3)-desaturase |
| Bacteroidetes1 | *Bacteroidetes* bacterium SW_11_45_7 | PSR04501.1 | Fatty acid desaturase |
| Bacteroidetes2 | *Bacteroidetes* bacterium 46-16 | OJW85131.1 | Fatty acid desaturase |
| B. distachyon | *Brachypodium distachyon* | XP_003578001.2 | Delta(8)-fatty-acid desaturase 2 |
| B. floridae | *Branchiostoma floridae* | XP_002586717.1 | Hypothetical protein BRAFLDRAFT_121704 |
| Burkholderia | *Burkholderia* sp. H160 | EEA04242.1 | Conserved hypothetical protein |
| C. follicularis1 | *Cephalotus follicularis* | GAV77917.1 | FA_desaturase domain-containing protein/Lipid_DES domain-containing protein |
| C. follicularis2 | *Cephalotus follicularis* | GAV56989.1 | Cyt-b5 domain-containing protein/FA_desaturase domain-containing protein |
| C. p. bellii | *Chrysemys picta bellii* | XP_005283813.1 | Sphingolipid delta(4)-desaturase/C4-monooxygenase DES2-like |
| C. reinhardtii | *Chlamydomonas reinhardtii* | XP_001691564.1 | Predicted protein |
| C. roenbergensis1 | *Cafeteria roenbergensis* | KAA0156831.1 | Hypothetical protein FNF29_00941 |
| C. roenbergensis2 | *Cafeteria roenbergensis* | KAA0153464.1 | Hypothetical protein FNF28_06948 |
| C. roenbergensis3 | *Cafeteria roenbergensis* | KAA0174097.1 | Hypothetical protein FNF27_04483 |
| C. tobini1 | *Chrysochromulina tobini* | KOO24852.1 | Sphingolipid delta -desaturase des1-like protein |
| C. tobini2 | *Chrysochromulina tobini* | KOO20797.1 | Fatty acid desaturase |
| C. tobini3 | *Chrysochromulina tobini* | KOO29180.1 | Hypothetical protein Ctob_007971 |
| C. virginica | *Crassostrea virginica* | XP_022329012.1 | Sphingolipid delta(4)-desaturase DES1-like |
| D. melanogaster | *Drosophila melanogaster* | NP_476594.1 | Infertile crescent, isoform A |
| D. pulex | *Daphnia pulex* | EFX83396.1 | Hypothetical protein DAPPUDRAFT_48184 |
| E. pallida1 | *Exaiptasia pallida (Exaiptasia diaphana)* | XP_020891967.1 | Sphingolipid delta(4)-desaturase DES1 |
| E. pallida2 | *Exaiptasia pallida (Exaiptasia diaphana)* | XP_020912814.2, XP_020912805.1 | Delta(8)-fatty-acid desaturase |
| E. siliculosus | *Ectocarpus siliculosus* | CBN74378.1 | Fatty acid desaturase |
| F. ambrosium | *Fusarium ambrosium* | RSM00058.1 | Hypothetical protein CDV31_011915 |
| G. cichoracearum | *Golovinomyces cichoracearum* | RKF82449.1 | Sphingolipid delta-desaturase |
| H. impetiginosus1 | *Handroanthus impetiginosus* | PIN19733.1 | Fatty acid desaturase |
| H. impetiginosus2 | *Handroanthus impetiginosus* | PIN06828.1 | Delta 6-fatty acid desaturase/delta-8 sphingolipid desaturase |
| I. galbana | *Isochrysis galbana* | AEV77089.1 | Delta-6 fatty acid desaturase |
| Isochrysis | *Isochrysis sp. CCMM5001* | AFB82637.1 | Fatty acid desaturase |
| K. nitens1 | *Klebsormidium nitens* | GAQ87926.1 | Dihydrosphingosine delta-4 desaturase |
| K. nitens2 | *Klebsormidium nitens* | GAQ87984.1 | Hypothetical protein KFL_003920030 |
| K. nitens3 | *Klebsormidium nitens* | GAQ79919.1 | Sphingobase-D8 Desaturase |
| M. pusilla1 | *Micromonas pusilla CCMP1545* | XP_003064164.1 | Predicted protein |
| M. pusilla2 | *Micromonas pusilla CCMP1545* | XP_003054909.1 | Predicted protein |
| M. pusilla3 | *Micromonas pusilla CCMP1545* | XP_003063519.1 | Predicted protein |
| M. pusilla4 | *Micromonas pusilla CCMP1545* | XP_003055443.1 | Predicted protein |
| M. rosea | *Minicystis rosea* | WP_146730508.1 | Fatty acid desaturase |
| N. colorata1 | *Nymphaea colorata* | XP_031478825.1 | Sphingolipid delta(4)-desaturase DES1-like |
| N. colorata2 | *Nymphaea colorata* | XP_031504778.1 | Delta(8)-fatty-acid desaturase-like |
| N. vectensis | *Nematostella vectensis* | XP_001640617.1 | Delta(8)-fatty-acid desaturase |
| O. bimaculoides | *Octopus bimaculoides* | XP_014781463.1 | Predicted: sphingolipid delta(4)-desaturase DES1-like |
| O. sativa1<sup>†</sup> | *Oryza sativa Japonica* Group | XP_015623789.1 | Sphingolipid delta(4)-desaturase DES1-like |
| O. sativa2 | *Oryza sativa Japonica* Group | XP_015651259.1 | Delta(8)-fatty-acid desaturase 2 |
| O. tauri1 | *Ostreococcus tauri* | XP_003082334.1 | Fatty acid desaturase, type 1 |
| O. tauri2 | *Ostreococcus tauri* | OUS49176.1 | Fatty acid desaturase-domain-containing protein |
| P. miliaceum1 | *Panicum miliaceum* | RLM73192.1 | Sphingolipid delta(4)-desaturase DES1-like |
| P. miliaceum2 | *Panicum miliaceum* | RLN34870.1 | Delta(8)-fatty-acid desaturase 2-like |
| P. patens1 | *Physcomitrella patens* | XP_024361943.1 | Sphingolipid delta(4)-desaturase DES1-like |
| P. patens2 | *Physcomitrella patens* | XP_024364920.1 | Acyl-lipid (9-3)-desaturase-like |
| P. roqueforti | *Penicillium roqueforti* FM164 | CDM35784.1 | Fatty acid desaturase, type 1 |
| P. trichocarpa1 | *Populus trichocarpa* | XP_006377338.2 | Sphingolipid delta(4)-desaturase DES1-like |
| P. trichocarpa2 | *Populus trichocarpa* | XP_002308556.1 | Acyl-lipid (9-3)-desaturase |
| S. asiatica1 | *Striga asiatica* | GER36468.1 | Sphingolipid delta(4)-desaturase DES1 |
| S. asiatica2 | *Striga asiatica* | GER35419.1 | Fatty acid desaturase |
| S. microadriaticum | *Symbiodinium microadriaticum* | OLP82839.1 | Delta(8)-fatty-acid desaturase |
| S. moellendorffii1 | *Selaginella moellendorffii* | XP_002971294.1, XP_002961512.1 | Sphingolipid delta(4)-desaturase DES1-like |
| S. moellendorffii2 | *Selaginella moellendorffii* | XP_002968817.1 | Delta(8)-fatty-acid desaturase 2 |
| Sphingobacteriales1 | *Sphingobacteriales* bacterium | RYE19069.1 | Fatty acid desaturase, partial |
| Sphingobacteriales2 | *Sphingobacteriales* bacterium 48-107 | OJW43059.1 | Fatty acid desaturase |
| Synechococcus | *Synechococcus* sp. PCC 7336 | WP_156820318.1 | Fatty acid desaturase |
| T. cacao1 | *Theobroma cacao* | XP_007025663.2 | Predicted: sphingolipid delta(4)-desaturase DES1-like |
| T. cacao2 | *Theobroma cacao* | XP_007012291.1 | Predicted: acyl-lipid (9-3)-desaturase |
| T. pseudonana | *Thalassiosira pseudonana* CCMP1335 | XP_002291331.1 | Predicted protein |
| T. turgidum1 | *Triticum turgidum* subsp. durum | VAH49645.1 | Unnamed protein product |
| T. turgidum2 | *Triticum turgidum* subsp. durum | VAI17523.1 | Unnamed protein product |
| W. hederac | *Wallemia hederac* | TIA87401.1 | Hypothetical protein E3P99_03193 |

<sup>†</sup>Functionally characterized proteins.

**Table S6:** Information regarding the proteins used to build the SBH phylogenetic tree.

| Name | Organism | Accession number | Description |
| --- | --- | --- | --- |
| E. huxleyi_SBH1 | *Emiliana huxleyi* CCMP2090, CCMP373 | MZ152820, MZ152819 | Sphingoid base hydroxylase 1 |
| E. huxleyi_SBH2 | *Emiliana huxleyi* CCMP2090 | MZ152821 | Sphingoid base hydroxylase 2 |
| E. huxleyi_SBH3 | *Emiliana huxleyi* | Prediction from genome | Sphingoid base hydroxylase 3 |
| E. huxleyi_SBH4 | *Emiliana huxleyi* CCMP373 | MZ152822 | Sphingoid base hydroxylase 4 |
| E. huxleyi_SBH5 | *Emiliana huxleyi* CCMP373 | MZ152823 | Sphingoid base hydroxylase 5 |
| E. huxleyi_SBH6 | *Emiliana huxleyi* CCMP2090, CCMP373 | MZ152825, MZ152824 | Sphingoid base hydroxylase 6 |
| E. huxleyi_SBH7 | *Emiliana huxleyi* CCMP2090, CCMP373 | MZ152826, MZ152827 | Sphingoid base hydroxylase 7 |
| EhV201_SBH | *Emiliana huxleyi* virus 201 | AET97919.1 | Hypothetical protein EPVG_00031 |
| A. candidus1 | *Aspergillus candidus* | XP_024676610.1 | Putative C-4 methylsterol oxidase |
| A. candidus2 | *Aspergillus candidus* | XP_024670972.1 | Putative C-4 methyl sterol oxidase |
| A. castellanii1 | *Acanthamoeba castellanii* str. Neff | XP_004336833.1 | 4Alpha-methyl-sterol C4-methyl-oxidase |
| A. castellanii2 | *Acanthamoeba castellanii* str. Neff | XP_004336864.1 | C5orf4 protein |
| A. mulundensis | *Aspergillus mulundensis* | XP_026600416.1 | Uncharacterized protein DSM5745_09314 |
| A. queenslandica | *Amphimedon queenslandica* | XP_011404818.2 | PREDICTED: methylsterol monooxygenase 1-like |
| Archaeon1 | archaeon | RYY81668.1 | Fatty acid hydroxylase family protein |
| Archaeon2 | archaeon | RYH18502.1 | Fatty acid hydroxylase family protein |
| C. merolae | *Cyanidioschyzon merolae* strain 10D | XP_005537142.1 | Hypothetical protein, conserved |
| C. tobinii1 | *Chrysochromulina tobinii* | KOO21719.1 | c-4 Methylsterol oxidase |
| C. tobinii2 | *Chrysochromulina tobinii* | KOO23105.1 | Sterol desaturase |
| Deltaproteobacteria | *Deltaproteobacteria* bacterium | MAA78328.1 | Hypothetical protein CL916_03640 |
| D. purpureum | *Dictyostelium purpureum* | XP_003291805.1 | Hypothetical protein DICPUDRAFT_156442 |
| E. affinis | *Eurytemora affinis* | XP_023341469.1 | Fatty acid hydroxylase domain-containing protein 2-like |
| F. proliferatum | *Fusarium proliferatum* | RKL31181.1 | Hypothetical protein BFJ72_g11198 |
| H. fermentalgiana | *Hondaea fermentalgiana* | GBG26608.1 | Methylsterol monooxygenase 1-1 |
| H. sapiens1<sup>†</sup> | *Homo sapiens* | NP_006736.1 | Methylsterol monooxygenase 1 isoform 1 |
| H. sapiens2 | *Homo sapiens* | AAH04506.2 | C5orf4 protein, partial |
| M. brevicollis | *Monosiga brevicollis* MX1 | XP_001747965.1 | Hypothetical protein |
| M. commoda | *Micromonas commoda* | XP_002508762.1 | Predicted protein |
| O. tauri | *Ostreococcus tauri* | XP_003079549.2 | Fatty acid hydroxylase |
| P. marinus1 | *Perkinsus marinus* ATCC 50983 | XP_002782009.1 | Sterol desaturase, putative |
| P. marinus2 | *Perkinsus marinus* ATCC 50983 | XP_002771628.1 | Lathosterol oxidase, putative |
| P. olseni | *Perkinsus olseni* | KAF4694963.1 | Chromosome 5 4 |
| P. tetraurelia1 | *Paramecium tetraurelia* strain d4-2 | XP_001449651.1 | Hypothetical protein (macronuclear) |
| P. tetraurelia2 | *Paramecium tetraurelia* strain d4-2 | XP_001448034.1 | Hypothetical protein (macronuclear) |
| P. umbilicalis | *Porphyra umbilicalis* | OSX72141.1 | Hypothetical protein BU14_0463s0003 |
| S. cerevisiae<sup>†</sup> | *Saccharomyces cerevisiae* S288C | NP_010583.1 | Sphingosine hydroxylase |
| S. indica1 | *Serendipita indica* DSM 11827 | CCA68111.1 | Related to C-4 methyl sterol oxidase |
| S. indica2 | *Serendipita indica* DSM 11827 | CCA69868.1 | Probable ERG25-C-4 methyl sterol oxidase |
| S. microadriaticum | *Symbiodinium microadriaticum* | OLP85489.1 | Fatty acid hydroxylase domain-containing protein 2 |
| S. pombe | *Schizosaccharomyces pombe* | NP_596489.1 | Sphingosine hydroxylase Sur2 |
| S. rosetta1 | *Salpingoeca rosetta* | XP_004992472.1 | Hypothetical protein PTSG_07059 |
| S. rosetta2 | *Salpingoeca rosetta* | XP_004995906.1 | GTP binding protein 4 |
| T. trahens1 | *Thecamonas trahens* ATCC 50062 | XP_013758029.1 | 4-Alpha-methyl-sterol C4-methyl-oxidase |
| T. trahens2 | *Thecamonas trahens* ATCC 50062 | XP_013761304.1 | Sterol desaturase |

<sup>†</sup>Functionally characterized proteins.

**Table S7:** Correlations between the abundance of *E. huxleyi* and EhV and the concentration of different GSL species in four bags during a mesocosm experiment.

|  | *E. huxleyi* | hGSL<br>19:3/22:2 | sGSL<br>d18:2/c22:0 | Biomass-associated EhV | vGSL<br>t17:0/h22:0 |
| --- | --- | --- | --- | --- | --- |
| *E. huxleyi* | 1 | 0.73 | 0.72 | -0.1 | -0.01 |
| GSL d18:3/h22:1 (1) | 0.75 | 0.76 | 0.72 | 0.03 | 0.1 |
| hGSL 19:3/22:1 | 0.66 | 0.7 | 0.65 | -0.08 | -0.01 |
| hGSL 19:3/22:2 | 0.73 | 1 | 0.86 | 0.07 | 0.18 |
| GSL d18:3/h22:2 (2) | 0.65 | 0.97 | 0.81 | 0.15 | 0.26 |
| sGSL d18:2/c22:0 | 0.72 | 0.86 | 1 | 0.03 | 0.16 |
| 374-GSL d19:3/h22:2 (13) | 0.75 | 0.92 | 0.84 | -0.06 | 0.06 |
| GSL d19:3/h21:1 (3) | 0.57 | 0.92 | 0.74 | 0.22 | 0.32 |
| GSL d19:3/h23:2 (4) | 0.53 | 0.89 | 0.7 | 0.28 | 0.34 |
| sGSL d18:1/c22:0 | 0.58 | 0.86 | 0.75 | 0.17 | 0.3 |
| 374-GSL d19:3/h22:3 (14) | 0.59 | 0.88 | 0.79 | 0.13 | 0.25 |
| Biomass-associated EhV | -0.1 | 0.07 | 0.03 | 1 | 0.9 |
| vGSL t17:0/h23:1 | -0.07 | 0.11 | 0.11 | 0.95 | 0.98 |
| vGSL t17:0/h22:1 | -0.08 | 0.09 | 0.1 | 0.95 | 0.98 |
| vGSL t16:0/h22:0 | -0.06 | 0.11 | 0.12 | 0.92 | 0.99 |
| vGSL t17:0/h24:1 | -0.07 | 0.1 | 0.09 | 0.91 | 0.97 |
| GSL d18:0/h22:1 (6) | -0.03 | 0.15 | 0.14 | 0.84 | 0.96 |
| GSL d18:0/h22:0 (5) | -0.06 | 0.1 | 0.15 | 0.87 | 0.94 |
| vGSL t17:0/h23:0 | 0.01 | 0.2 | 0.18 | 0.87 | 1 |
| vGSL t17:0/h24:0 | 0 | 0.21 | 0.18 | 0.87 | 1 |
| GSL t18:0/h22:0 (8) | 0.01 | 0.19 | 0.2 | 0.85 | 0.99 |
| vGSL t17:0/h22:0 | -0.01 | 0.18 | 0.16 | 0.9 | 1 |
| GSL t18:0/h22:1 (9) | -0.03 | 0.15 | 0.15 | 0.89 | 0.99 |
| GSL t18:0/h22:2 (10) | -0.19 | -0.11 | -0.05 | 0.85 | 0.69 |

Rows and columns are organized based on hierarchal clustering (using the 'R' package 'heatmap'). Pearson correlation coefficient ( $r$ ) values are presented using a blue-red color scheme.

**Table S8:** ‘xcms’ parameters used for peak picking.

| Parameter | Value |
| --- | --- |
| fwhm | 15 |
| snthresh | 9 |
| mzdiff | 0.01 |

**Table S9:**
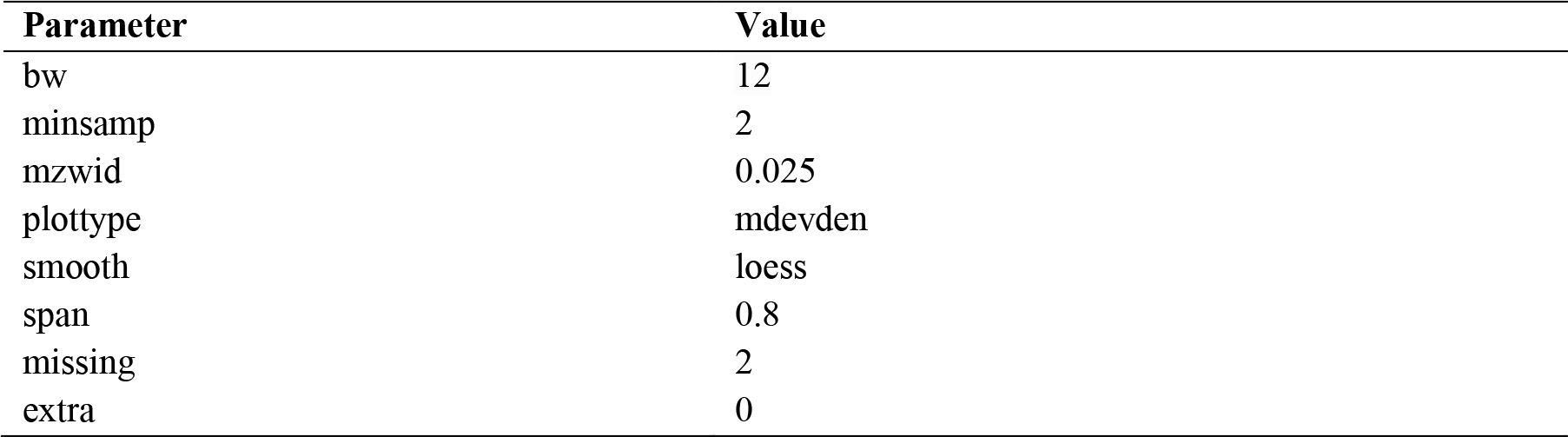
‘xcms’ parameters used for peak grouping and alignment.

**Table S10:**
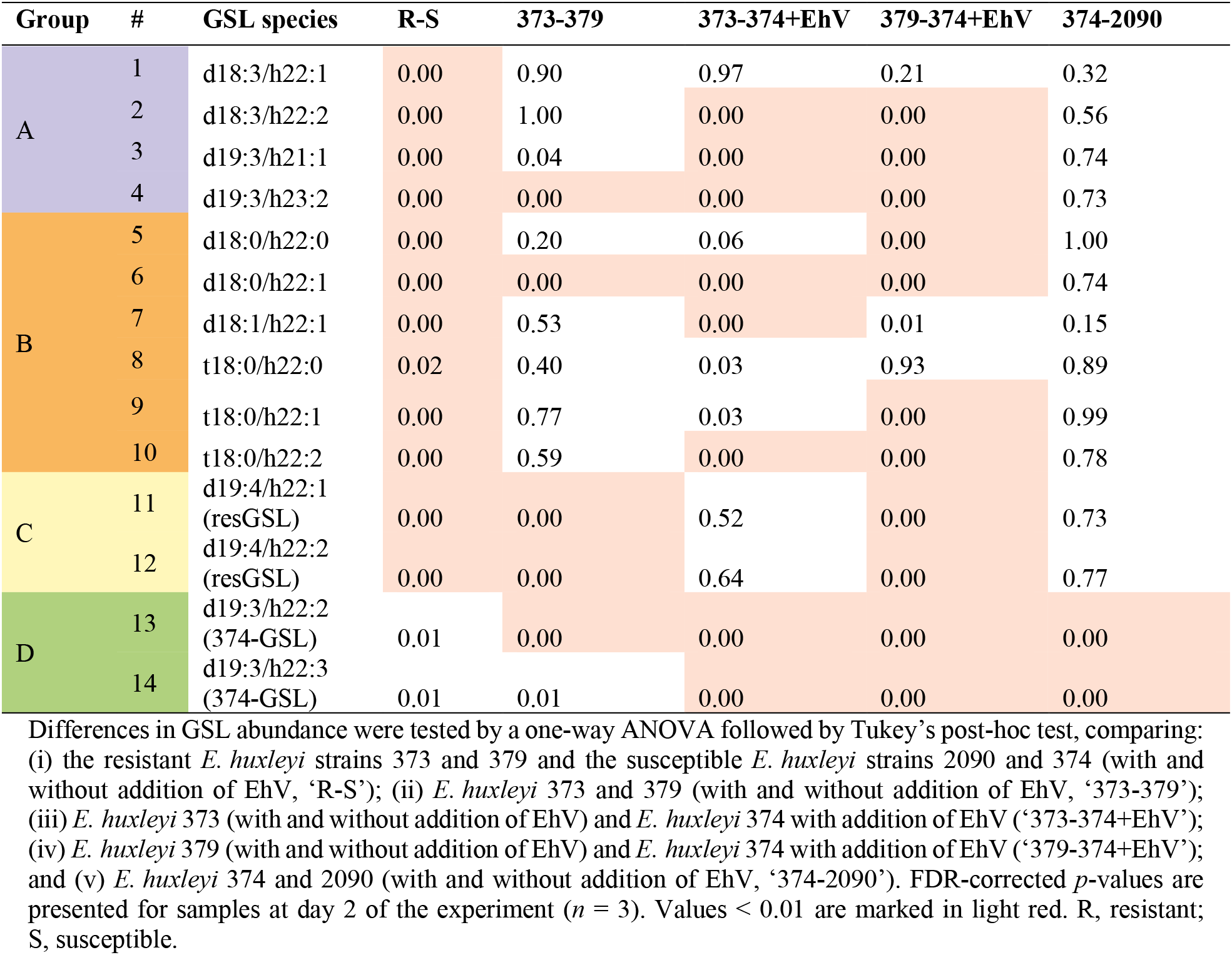
Tukey’s multiple pairwise comparison tests of GSL species in resistant and susceptible *E. huxleyi* strains.

| Group | # | GSL species | R-S | 373-379 | 373-374+EhV | 379-374+EhV | 374-2090 |
| --- | --- | --- | --- | --- | --- | --- | --- |
| A | 1 | d18:3/h22:1 | 0.00 | 0.90 | 0.97 | 0.21 | 0.32 |
|  | 2 | d18:3/h22:2 | 0.00 | 1.00 | 0.00 | 0.00 | 0.56 |
|  | 3 | d19:3/h21:1 | 0.00 | 0.04 | 0.00 | 0.00 | 0.74 |
|  | 4 | d19:3/h23:2 | 0.00 | 0.00 | 0.00 | 0.00 | 0.73 |
| B | 5 | d18:0/h22:0 | 0.00 | 0.20 | 0.06 | 0.00 | 1.00 |
|  | 6 | d18:0/h22:1 | 0.00 | 0.00 | 0.00 | 0.00 | 0.74 |
|  | 7 | d18:1/h22:1 | 0.00 | 0.53 | 0.00 | 0.01 | 0.15 |
|  | 8 | t18:0/h22:0 | 0.02 | 0.40 | 0.03 | 0.93 | 0.89 |
|  | 9 | t18:0/h22:1 | 0.00 | 0.77 | 0.03 | 0.00 | 0.99 |
|  | 10 | t18:0/h22:2 | 0.00 | 0.59 | 0.00 | 0.00 | 0.78 |
| C | 11 | d19:4/h22:1<br>(resGSL) | 0.00 | 0.00 | 0.52 | 0.00 | 0.73 |
|  | 12 | d19:4/h22:2<br>(resGSL) | 0.00 | 0.00 | 0.64 | 0.00 | 0.77 |
| D | 13 | d19:3/h22:2<br>(374-GSL) | 0.01 | 0.00 | 0.00 | 0.00 | 0.00 |
|  | 14 | d19:3/h22:3<br>(374-GSL) | 0.01 | 0.01 | 0.00 | 0.00 | 0.00 |

Differences in GSL abundance were tested by a one-way ANOVA followed by Tukey's post-hoc test, comparing: (i) the resistant *E. huxleyi* strains 373 and 379 and the susceptible *E. huxleyi* strains 2090 and 374 (with and without addition of EhV, 'R-S'); (ii) *E. huxleyi* 373 and 379 (with and without addition of EhV, '373-379'); (iii) *E. huxleyi* 373 (with and without addition of EhV) and *E. huxleyi* 374 with addition of EhV ('373-374+EhV'); (iv) *E. huxleyi* 379 (with and without addition of EhV) and *E. huxleyi* 374 with addition of EhV ('379-374+EhV'); and (v) *E. huxleyi* 374 and 2090 (with and without addition of EhV, '374-2090'). FDR-corrected *p*-values are presented for samples at day 2 of the experiment (*n* = 3). Values < 0.01 are marked in light red. R, resistant; S, susceptible.

**Table S11:** Abundance ratios of GSL species in resistant and susceptible *E. huxleyi* strains.

| Group | # | GSL species | R/S | 373/379 | 373/374+EhV | 379/374+EhV | 374/2090 |
| --- | --- | --- | --- | --- | --- | --- | --- |
| A | 1 | d18:3/h22:1 | 15 | 0.6 | 0.9 | 1.5 | 3.4 |
|  | 2 | d18:3/h22:2 | 44 | 1.0 | 18 | 19 | 0.4 |
|  | 3 | d19:3/h21:1 | 87 | 1.4 | 76 | 55 | 1.1 |
|  | 4 | d19:3/h23:2 | 1314 | 2.8 | 1649 | 580 | 1.1 |
| B | 5 | d18:0/h22:0 | 140 | 0.1 | 7.1 | 57 | 1.2 |
|  | 6 | d18:0/h22:1 | 776 | 0.2 | 289 | 1155 | 1.1 |
|  | 7 | d18:1/h22:1 | 65 | 0.2 | 0.3 | 1.4 | 13 |
|  | 8 | t18:0/h22:0 | 6.0 | 0.2 | 0.2 | 0.8 | 2.1 |
|  | 9 | t18:0/h22:1 | 540 | 0.3 | 23 | 79 | 1.5 |
|  | 10 | t18:0/h22:2 | 681 | 0.9 | 469 | 547 | 1.1 |
| C | 11 | d19:4/h22:1<br>(resGSL) | 11 | 0.0 | 0.9 | 79 | 1.1 |
|  | 12 | d19:4/h22:2<br>(resGSL) | 33 | 0.0 | 0.9 | 680 | 1.1 |
| D | 13 | d19:3/h22:2<br>(374-GSL) | 0.1 | 0.7 | 0 | 0 | 212 |
|  | 14 | d19:3/h22:3<br>(374-GSL) | 0.2 | 0.7 | 0 | 0 | 77 |

## References

1 Bergh, Ø., Børsheim, K. Y., Bratbak, G. & Heldal, M. High abundance of viruses found in aquatic environments. Nature 340, 467–468 (1989).

2 Wommack, K. E. & Colwell, R. R. Virioplankton: Viruses in aquatic ecosystems. Microbiol. Mol. Biol. Rev. 64, 69–114 (2000).

3 Weitz, J. S. et al. A multitrophic model to quantify the effects of marine viruses on microbial food webs and ecosystem processes. ISME J. 9, 1352–1364 (2015).

4 Suttle, C. A. Marine viruses — major players in the global ecosystem. Nat. Rev. Microbiol. 5, 801–812 (2007).

5 Fuhrman, J. A. Marine viruses and their biogeochemical and ecological effects. Nature 399, 541–548 (1999).

6 Wilhelm, S. W. & Suttle, C. A. Viruses and nutrient cycles in the sea: Viruses play critical roles in the structure and function of aquatic food webs. BioScience 49, 781–788 (1999).

7 Sullivan, M. B., Weitz, J. S. & Wilhelm, S. Viral ecology comes of age. Environ. Microbiol. Rep. 9, 33–35 (2017).

8 Baran, N., Goldin, S., Maidanik, I. & Lindell, D. Quantification of diverse virus populations in the environment using the polony method. Nat. Microbiol. 3, 62–72 (2018).

9 Pasulka, A. L. et al. Interrogating marine virus-host interactions and elemental transfer with BONCAT and nanoSIMS-based methods. Environ. Microbiol. 20, 671–692 (2018).

10 Vincent, F., Sheyn, U., Porat, Z., Schatz, D. & Vardi, A. Visualizing active viral infection reveals diverse cell fates in synchronized algal bloom demise. Proc. Natl. Acad. Sci. USA 118, e2021586118 (2021).

11 Allers, E. et al. Single-cell and population level viral infection dynamics revealed by phageFISH, a method to visualize intracellular and free viruses. Environ. Microbiol. 15, 2306–2318 (2013).

12 Castillo, Y. M. et al. Visualization of viral infection dynamics in a unicellular eukaryote and quantification of viral production using virus fluorescence in situ hybridization. Front. Microbiol. 11, 1559 (2020).

13 Ku, C. et al. A single-cell view on alga-virus interactions reveals sequential transcriptional programs and infection states. Sci. Adv. 6, eaba4137 (2020).

14 Piel, D. et al. Phage–host coevolution in natural populations. Nat. Microbiol. 7, 1075–1086 (2022).

15 Yau, S., et al. A viral immunity chromosome in the marine picoeukaryote, Ostreococcus tauri. PLoS Path. 12, e1005965 (2016).

16 Frada, M., Probert, I., Allen, M. J., Wilson, W. H. & de Vargas, C. The “Cheshire Cat” escape strategy of the coccolithophore Emiliania huxleyi in response to viral infection. Proc. Natl. Acad. Sci. USA 105, 15944–15949 (2008).

17 Jacobsen, A., Larsen, A., Martínez Martínez, J., Verity, P. G. & Frischer, M. E. Susceptibility of colonies and colonial cells of Phaeocystis pouchetii (Haptophyta) to viral infection. Aquat. Microb. Ecol. 48, 105–112 (2007).

18 Martínez Martínez, J., et al. New lipid envelope-containing dsDNA virus isolates infecting Micromonas pusilla reveal a separate phylogenetic group. Aquat. Microb. Ecol. 74, 17–28 (2015).

19 Tarutani, K., Nagasaki, K. & Yamaguchi, M. Viral impacts on total abundance and clonal composition of the harmful bloom-forming phytoplankton Heterosigma akashiwo. Appl. Environ. Microbiol. 66, 4916–4920 (2000).

20 Tomaru, Y., Mizumoto, H. & Nagasaki, K. Virus resistance in the toxic bloom-forming dinoflagellate Heterocapsa circularisquama to single-stranded RNA virus infection. Environ. Microbiol. 11, 2915–2923 (2009).

21 Thomas, R. et al. Acquisition and maintenance of resistance to viruses in eukaryotic phytoplankton populations. Environ. Microbiol. 13, 1412–1420 (2011).

22 Schroeder, D. C., Oke, J., Malin, G. & Wilson, W. H. Coccolithovirus (Phycodnaviridae): Characterisation of a new large dsDNA algal virus that infects Emiliana huxleyi. Arch. Virol. 147, 1685–1698 (2002).

23 Holligan, P. M. et al. A biogeochemical study of the coccolithophore, Emiliania huxleyi, in the North Atlantic. Global Biogeochem. Cycles 7, 879–900 (1993).

24 Brown, C. W. & Yoder, J. A. Coccolithophorid blooms in the global ocean. J. Geophys. Res. 99, 7467–7482 (1994).

25 Harris, R. P. Zooplankton grazing on the coccolithophore Emiliania huxleyi and its role in inorganic carbon flux. Mar. Biol. 119, 431–439 (1994).

26 Stefels, J., Steinke, M., Turner, S., Malin, G. & Belviso, S. Environmental constraints on the production and removal of the climatically active gas dimethylsulphide (DMS) and implications for ecosystem modelling. Biogeochemistry 83, 245–275 (2007).

27 Bratbak, G., Egge, J. K. & Heldal, M. Viral mortality of the marine alga Emiliania huxleyi (Haptophyceae) and termination of algal blooms. Mar. Ecol. Prog. Ser. 93, 39–48 (1993).

28 Lehahn, Y. et al. Decoupling physical from biological processes to assess the impact of viruses on a mesoscale algal bloom. Curr. Biol. 24, 2041–2046 (2014).

29 Laber, C. P. et al. Coccolithovirus facilitation of carbon export in the North Atlantic. Nat. Microbiol. 3, 537–547 (2018).

30 Wilson, W. H. et al. Isolation of viruses responsible for the demise of an Emiliania huxleyi bloom in the English Channel. J. Mar. Biol. Assoc. U.K. 82, 369–377 (2002).

31 Rosenwasser, S. et al. Rewiring host lipid metabolism by large viruses determines the fate of Emiliania huxleyi, a bloom-forming alga in the ocean. Plant Cell 26, 2689–2707 (2014).

32 Malitsky, S. et al. Viral infection of the marine alga Emiliania huxleyi triggers lipidome remodeling and induces the production of highly saturated triacylglycerol. New Phytol. 210, 88–96 (2016).

33 Schleyer, G. et al. In plaque-mass spectrometry imaging of a bloom-forming alga during viral infection reveals a metabolic shift towards odd-chain fatty acid lipids. Nat. Microbiol. 4, 527–538 (2019).

34 Evans, C., Pond, D. W. & Wilson, W. H. Changes in Emiliania huxleyi fatty acid profiles during infection with E. huxleyi virus 86: Physiological and ecological implications. Aquat. Microb. Ecol. 55, 219–228 (2009).

35 Wilson, W. H. et al. Complete genome sequence and lytic phase transcription profile of a Coccolithovirus. Science 309, 1090–1092 (2005).

36 Ziv, C. et al. Viral serine palmitoyltransferase induces metabolic switch in sphingolipid biosynthesis and is required for infection of a marine alga. Proc. Natl. Acad. Sci. USA 113, E1907–E1916 (2016).

37 Vardi, A. et al. Viral glycosphingolipids induce lytic infection and cell death in marine phytoplankton. Science 326, 861–865 (2009).

38 Fulton, J. M. et al. Novel molecular determinants of viral susceptibility and resistance in the lipidome of Emiliania huxleyi. Environ. Microbiol. 16, 1137–1149 (2014).

39 Vardi, A. et al. Host-virus dynamics and subcellular controls of cell fate in a natural coccolithophore population. Proc. Natl. Acad. Sci. USA 109, 19327–19332 (2012).

40 Feldmesser, E., Ben-Dor, S. & Vardi, A. An Emiliania huxleyi pan-transcriptome reveals basal strain specificity in gene expression patterns. Sci. Rep. 11, 20795–20795 (2021).

41 Bidle, K. D. & Kwityn, C. J. Assessing the role of caspase activity and metacaspase expression on viral susceptibility of the coccolithophore, Emiliania huxleyi (Haptophyta). J. Phycol. 48, 1079–1089 (2012).

42 Frada, M. J., et al. Morphological switch to a resistant subpopulation in response to viral infection in the bloom-forming coccolithophore Emiliania huxleyi. PLoS Path. 13, e1006775 (2017).

43 Nissimov, J. I. et al. Biochemical diversity of glycosphingolipid biosynthesis as a driver of Coccolithovirus competitive ecology. Environ. Microbiol. 21, 2182–2197 (2019).

44 Sud, M. et al. LMSD: LIPID MAPS structure database. Nucleic Acids Res. 35, D527–D532 (2007).

45 Hunter, J. E., Frada, M. J., Fredricks, H. F., Vardi, A. & Van Mooy, B. A. S. Targeted and untargeted lipidomics of Emiliania huxleyi viral infection and life cycle phases highlights molecular biomarkers of infection, susceptibility, and ploidy. Front. Mar. Sci. 2, 81 (2015).

46 Sumner, L. W. et al. Proposed minimum reporting standards for chemical analysis Chemical Analysis Working Group (CAWG) Metabolomics Standards Initiative (MSI). Metabolomics 3, 211–221 (2007).

47 Pagarete, A., Allen, M. J., Wilson, W. H., Kimmance, S. a. & de Vargas, C. Host-virus shift of the sphingolipid pathway along an Emiliania huxleyi bloom: Survival of the fattest. Environ. Microbiol. 11, 2840–2848 (2009).

48 Keeling, P. J. et al. The Marine Microbial Eukaryote Transcriptome Sequencing Project (MMETSP): Illuminating the functional diversity of eukaryotic life in the oceans through transcriptome sequencing. PLoS Biol. 12, e1001889 (2014).

49 Michaelson, L. V., Dunn, T. M. & Napier, J. A. Viral trans-dominant manipulation of algal sphingolipids. Trends Plant Sci. 15, 651–655 (2010).

50 Câmara dos Reis, M., et al. Exploring the phycosphere of Emiliania huxleyi: From bloom dynamics to microbiome assembly experiments. bioRxiv, 2022.2002.2021.481256 (2022).

51 Vincent, F., et al. Viral infection switches the balance between bacterial and eukaryotic recyclers of organic matter during algal blooms. bioRxiv (2021).

52 Kuhlisch, C. et al. Viral infection of algal blooms leaves a unique metabolic footprint on the dissolved organic matter in the ocean. Sci. Adv. 7, eabf4680 (2021).

53 Egge, J. K. & Heimdal, B. R. Blooms of phytoplankton including Emiliania huxleyi (Haptophyta). Effects of nutrient supply in different N : P ratios. Sarsia 79, 333–348 (1994).

54 Fromm, A., Schatz, D., Ben-Dor, S., Feldmesser, E. & Vardi, A. Complete genome sequence of Emiliania huxleyi virus strain M1, isolated from an induced E. huxleyi bloom in Bergen, Norway. Microbiol. Resour. Announc. 11, e00071–22 (2022).

55 Waterbury, J. B. & Valois, F. W. Resistance to co-occurring phages enables marine Synechococcus communities to coexist with cyanophages abundant in seawater. Appl. Environ. Microbiol. 59, 3393–3399 (1993).

56 Avrani, S., Wurtzel, O., Sharon, I., Sorek, R. & Lindell, D. Genomic island variability facilitates Prochlorococcus–virus coexistence. Nature 474, 604–608 (2011).

57 Kegel, J. U., John, U., Valentin, K. & Frickenhaus, S. Genome variations associated with viral susceptibility and calcification in Emiliania huxleyi. PLoS ONE 8, e80684 (2013).

58 Li, Y. et al. Sphingolipids in marine microalgae: Development and application of a mass spectrometric method for global structural characterization of ceramides and glycosphingolipids in three major phyla. Anal. Chim. Acta 986, 82–94 (2017).

59 Li, H. et al. Comparative lipid profile of four edible shellfishes by UPLC-Triple TOF-MS/MS. Food Chem. 310, 125947 (2020).

60 Wang, R. et al. Identification of ceramide 2-aminoethylphosphonate molecular species from different aquatic products by NPLC/Q-Exactive-MS. Food Chem. 304, 125425 (2020).

61 Kato, H. et al. Recognition of pathogen-derived sphingolipids in Arabidopsis. Science 376, 857–860 (2022).

62 Marquês, J. T., Marinho, H. S. & de Almeida, R. F. M. Sphingolipid hydroxylation in mammals, yeast and plants – An integrated view. Prog. Lipid Res. 71, 18–42 (2018).

63 Santos, T. C. B. et al. The long chain base unsaturation has a stronger impact on 1-deoxy(methyl)-sphingolipids biophysical properties than the structure of its C1 functional group. Biochim. Biophys. Acta 1863, 183628 (2021).

64 Ali, U., Li, H., Wang, X. & Guo, L. Emerging roles of sphingolipid signaling in plant response to biotic and abiotic stresses. Mol. Plant 11, 1328–1343 (2018).

65 Zhou, Y. et al. The sphingolipid biosynthetic enzyme Sphingolipid delta8 desaturase is important for chilling resistance of tomato. Sci. Rep. 6, 38742–38742 (2016).

66 Mamode Cassim, A., et al. Sphingolipids in plants: A guidebook on their function in membrane architecture, cellular processes, and environmental or developmental responses. FEBS Lett. 594, 3719–3738 (2020).

67 Schneider-Schaulies, J. & Schneider-Schaulies, S. Sphingolipids in viral infection. Biol. Chem. 396, 585–595 (2015).

68 Orchard, R. C., Wilen, C. B. & Virgin, H. W. Sphingolipid biosynthesis induces a conformational change in the murine norovirus receptor and facilitates viral infection. Nat. Microbiol. 3, 1109–1114 (2018).

69 Törnquist, K., Asghar, M. Y., Srinivasan, V., Korhonen, L. & Lindholm, D. Sphingolipids as modulators of SARS-CoV-2 infection. Front. Cell Dev. Biol. 9, 689854 (2021).

70 Soudani, N., Hage-Sleiman, R., Karam, W., Dbaibo, G. & Zaraket, H. Ceramide suppresses influenza a virus replication in vitro. J. Virol. 93, e00053–19 (2019).

71 Bukrinsky, M. I., Mukhamedova, N. & Sviridov, D. Lipid rafts and pathogens: The art of deception and exploitation. J. Lipid Res. 61, 601–610 (2020).

72 Mackinder, L. C. M., et al. A unicellular algal virus, Emiliania huxleyi virus 86, exploits an animal-like infection strategy. J. Gen. Virol. 90, 2306–2316 (2009).

73 Grossi, V., Raphel, D., Aubert, C. & Rontani, J.-F. The effect of growth temperature on the long-chain alkenes composition in the marine coccolithophorid Emiliania huxleyi. Phytochemistry 54, 393–399 (2000).

74 Wördenweber, R. et al. Phosphorus and nitrogen starvation reveal life-cycle specific responses in the metabolome of Emiliania huxleyi (Haptophyta). Limnol. Oceanogr. 63, 203–226 (2017).

75 Lowenstein, D. P., Mayers, K., Fredricks, H. F. & Van Mooy, B. A. S. Targeted and untargeted lipidomic analysis of haptophyte cultures reveals novel and divergent nutrient-stress adaptations. Org. Geochem. 161, 104315 (2021).

76 Del Poeta, M., Nimrichter, L., Rodrigues, M. L. & Luberto, C. Synthesis and biological properties of fungal glucosylceramide. PLoS Path. 10, e1003832 (2014).

77 Mashima, R., Okuyama, T. & Ohira, M. Biosynthesis of long chain base in sphingolipids in animals, plants and fungi. Future Sci. OA 6, FSO434 (2019).

78 Ruiz, E., Oosterhof, M., Sandaa, R.-A. A., Larsen, A. & Pagarete, A. Emerging interaction patterns in the Emiliania huxleyi-EhV system. Viruses 9, 61 (2017).

79 Thyrhaug, R., Larsen, A., Thingstad, T. F. & Bratbak, G. Stable coexistence in marine algal host-virus systems. Mar. Ecol. Prog. Ser. 254, 27–35 (2003).

80 Baumeister, T. U. H., Vallet, M., Kaftan, F., Svatoš, A. & Pohnert, G. Live single-cell metabolomics with matrix-free laser/desorption ionization mass spectrometry to address microalgal physiology. Front. Plant Sci. 10, 172 (2019).

81 Li, Z. et al. Single-cell lipidomics with high structural specificity by mass spectrometry. Nat. Commun. 12, 2869 (2021).

82 Read, B. A. et al. Pan genome of the phytoplankton Emiliania underpins its global distribution. Nature 499, 209–213 (2013).

83 Nissimov, J. I. et al. Draft genome sequence of four coccolithoviruses: Emiliania huxleyi virus EhV-88, EhV-201, EhV-207, and EhV-208. J. Virol. 86, 2896-2897 (2012).

84 Gerecht, A. C., Šupraha, L., Edvardsen, B., Probert, I. & Henderiks, J. High temperature decreases the PIC / POC ratio and increases phosphorus requirements in Coccolithus pelagicus (Haptophyta). Biogeosciences 11, 3531–3545 (2014).

85 Barak-Gavish, N. et al. Bacterial virulence against an oceanic bloom-forming phytoplankter is mediated by algal DMSP. Sci. Adv. 4, eaau5716 (2018).

86 Hummel, J. et al. Ultra performance liquid chromatography and high resolution mass spectrometry for the analysis of plant lipids. Front. Plant Sci. 2, 54 (2011).

87 R Core Team. R: A language and environment for statistical computing R foundation for statistical computing. https://r-project.org. (2020).

88 Smith, C. A., Want, E. J., O’Maille, G., Abagyan, R. & Siuzdak, G. XCMS: Processing mass spectrometry data for metabolite profiling using nonlinear peak alignment, matching, and identification. Anal. Chem. 78, 779–787 (2006).

89 Kuhl, C., Tautenhahn, R., Böttcher, C., Larson, T. R. & Neumann, S. CAMERA: An integrated strategy for compound spectra extraction and annotation of liquid chromatography/mass spectrometry data sets. Anal. Chem. 84, 283–289 (2012).

90 Alboukadel, K. & Mundt, F. factoextra: Extract and visualize the results of multivariate data analyses. R package version 1.0.5. (2017).

91 Storey, J. D., Bass, A. J., Dabney, A. & Robinson, D. qvalue: Q-value estimation for false discovery rate control. R package version 2.14.1. (2019).

92 Feldmesser, E., Rosenwasser, S., Vardi, A. & Ben-Dor, S. Improving transcriptome construction in non-model organisms: Integrating manual and automated gene definition in Emiliania huxleyi. BMC Genomics 15, 148 (2014).

93 Altschul, S. F. et al. Gapped BLAST and PSI-BLAST: A new generation of protein database search programs. Nucleic Acids Res. 25, 3389–3402 (1997).

94 Finn, R. D. et al. Pfam: The protein families database. Nucleic Acids Res. 42, D222– D230 (2014).

95 Lu, S. et al. CDD/SPARCLE: The conserved domain database in 2020. Nucleic Acids Res. 48, D265–D268 (2020).

96 Larkin, M. A. et al. Clustal W and Clustal X version 2.0. Bioinformatics 23, 2947–2948 (2007).

97 Edgar, R. C. MUSCLE: Multiple sequence alignment with high accuracy and high throughput. Nucleic Acids Res. 32, 1792–1797 (2004).

98 Katoh, K., Rozewicki, J. & Yamada, K. D. MAFFT online service: Multiple sequence alignment, interactive sequence choice and visualization. Brief. Bioinform. 20, 1160–1166 (2019).

99 Felsenstein, J. PHYLIP (Phylogeny Inference Package) version 3.6. Distributed by the author. Department of Genome Sciences, University of Washington. (2004).

100 Guindon, S. et al. New algorithms and methods to estimate maximum-likelihood phylogenies: Assessing the performance of PhyML 3.0. Syst. Biol. 59, 307–321 (2010).

101 Letunic, I. & Bork, P. Interactive Tree Of Life (iTOL) v5: An online tool for phylogenetic tree display and annotation. Nucleic Acids Res. 49, W293–W296 (2021).

102 Sheyn, U. et al. Expression profiling of host and virus during a coccolithophore bloom provides insights into the role of viral infection in promoting carbon export. ISME J. 12, 704–713 (2018).

103 Vincent, F., et al. AQUACOSM VIMS-Ehux – Core data. Dryad (2020).

## References

1 Sud, M. et al. LMSD: Lipid Maps Structure Database. Nucleic Acids Res. 35, D527–D532 (2007).

2 Sumner, L. W. et al. Proposed minimum reporting standards for chemical analysis Chemical Analysis Working Group (CAWG) Metabolomics Standards Initiative (MSI). Metabolomics 3, 211–221 (2007).

3 Keeling, P. J. et al. The Marine Microbial Eukaryote Transcriptome Sequencing Project (MMETSP): Illuminating the functional diversity of eukaryotic life in the oceans through transcriptome sequencing. PLoS Biol. 12, e1001889 (2014).

4 Feldmesser, E., Ben-Dor, S. & Vardi, A. An Emiliania huxleyi pan-transcriptome reveals basal strain specificity in gene expression patterns. Sci. Rep. 11, 20795–20795 (2021).

5 Rosenwasser, S. et al. Rewiring host lipid metabolism by large viruses determines the fate of Emiliania huxleyi, a bloom-forming alga in the ocean. Plant Cell 26, 2689–2707 (2014).

6 Vardi, A. et al. Host-virus dynamics and subcellular controls of cell fate in a natural coccolithophore population. Proc. Natl. Acad. Sci. USA 109, 19327–19332 (2012).

7 Fulton, J. M. et al. Novel molecular determinants of viral susceptibility and resistance in the lipidome of Emiliania huxleyi. Environ. Microbiol. 16, 1137–1149 (2014).

8 Ziv, C. et al. Viral serine palmitoyltransferase induces metabolic switch in sphingolipid biosynthesis and is required for infection of a marine alga. Proc. Natl. Acad. Sci. USA 113, E1907–E1916 (2016).

